# Tissue Injury and Biomaterial Treatment Modulate Tumor Growth and Response to Immunotherapy

**DOI:** 10.64898/2026.02.02.703323

**Authors:** Anna Ruta, Elise F. Gray-Gaillard, Jordan A. Garcia, Locke Davenport Huyer, Yusef Mathkour, Christopher Cherry, Michael Patatanian, Joscelyn C. Mejías, David R. Maestas, Kavita Krishnan, Peter Abraham, Matthew T. Wolf, Kellie N. Smith, Drew M. Pardoll, Jennifer H. Elisseeff

## Abstract

Immunotherapies have transformed cancer care; however, tumor intrinsic and extrinsic factors contribute to high variability in therapeutic responses. While tissue injuries can impact cancer recurrence and metastatic spread, little is known about their potential to effect immune checkpoint blockade (ICB) response. In this study, we reveal that distal traumatic muscle injury accelerated progression and impaired adjuvant ICB response of multiple murine tumors. This injury-induced accelerated tumor growth coincided with decreased intra-tumoral density and effector phenotype of tumor-reactive CD8^+^ T cells and relied on communication through a shared draining lymph node. Enhancing injury repair using a biological scaffold abrogated the injury-induced accelerated tumor growth in an interleukin-4-dependent manner and improved ICB response. In a retrospective cohort analysis of breast cancer patients undergoing ICB treatment, biological scaffold implantation following mastectomy was associated with increased overall survival. This work suggests that injury-driven immune dysfunction may contribute to cancer progression and ICB resistance, but enhancing wound healing with pro-regenerative biomaterials may offer a viable strategy for mitigating adverse cancer outcomes, particularly in the setting of adjuvant and neoadjuvant ICB.

## Introduction

The tumor microenvironment (TME) can substantially influence cancer progression and therapeutic response. While immunotherapies, such as immune checkpoint blockade (ICB) therapy, have achieved unprecedented anti-tumor responses in many aggressive solid cancers, primary and acquired resistance limit sustained efficacy to only a small patient cohort. Their therapeutic value has expanded with adjuvant, and more recently neoadjuvant, ICB treatment for high-risk, surgically resectable cancers. Unfortunately, the majority of patients with “ICB-sensitive” cancers fail to respond long-term and mechanisms of ICB resistance remain to be clearly elucidated. Growing evidence suggests that host and environmental factors such as age, body mass index, gut microbiota composition, and diet can impact the TME and tumor response to ICB therapy^1–13^. Tissue injury, which triggers a cascade of local and systemic immune responses, is another such factor that may alter the TME. While prior studies demonstrate that tissue injury can contribute to tumor incidence, recurrence, and metastatic spread^14–19^, little is known about the potential contribution of tissue damage on tumor response to ICB therapy.

The link between neoplastic lesion formation and sites of tissue inflammation is supported by findings that numerous chronic inflammatory diseases (e.g., hepatitis, idiopathic pulmonary fibrosis, pancreatitis, inflammatory bowel disease) increase the risk of cancer development within corresponding damaged tissues^20–24^. Marjolin ulcers are similarly aggressive cutaneous malignancies that arise in pre-existing scar tissue and chronic wounds^25–29^. Further, retrospective clinical studies reveal that traumatic injury before cancer diagnosis and ischemic cardiovascular insults (e.g., myocardial infarction, heart failure, stroke) after cancer diagnosis can increase the risk of cancer-related mortality^30–34^. Surgical interventions can also inflict substantial tissue damage. Diagnostic biopsies, tumor resections, and subsequent tissue reconstruction are gold-standard procedures for treating many operable solid cancers; unfortunately, postoperative locoregional recurrence and metastatic spread are common^17,35–38^. Preclinical models further connect tissue injury and inflammation with poor cancer outcomes. For example, researchers observed incidence of sarcomas at distant wound sites^39^, increased colon tumor burden following laparotomy^40^, accelerated breast cancer progression after myocardial infarction^31^, and increased incidence of immunologically-controlled dormant tumors with a subcutaneous implant injury^41^. Since surgical interventions are increasingly coupled with adjuvant, and more recently neoadjuvant, ICB therapy with promising success, it is pertinent to consider whether injury-induced immune perturbations impact the TME and tumor response to ICB therapy.

Tissue injury incites a dynamic stromal-immune response characterized by an acute inflammatory phase followed by a prolonged remodeling and resolution phase^42–45^. Activated innate immune cells trigger the early pro-inflammatory response as they recognize damage-associated molecular patterns (DAMPs), eliminate wound site bacteria, and secrete chemoattractants to bolster immune cell recruitment^46^. While innate immunity is classically regarded as a key regulator of the wound healing response, the contribution of adaptive immunity to tissue repair and in governing healing versus fibrotic outcomes is increasingly recognized. Production of interferon-gamma (IFNγ) and interleukin (IL)-17 by innate and adaptive immune cells can regulate vascularization, neurogenesis, and re-epithelialization^47–50^. The subsequent remodeling phase consists of anti-inflammatory “M2” macrophages, type-2 helper T cells (T_H_2), and regulatory T cells (Tregs), which promote wound healing and restore tissue homeostasis by stimulating angiogenesis, re-epithelialization, and fibroblast secretion of extracellular matrix (ECM) components^42,43,51^. Since tumors co-opt many wound healing hallmarks, albeit in a highly dysregulated nature, they can be considered “wounds that do not heal”^52,53^.

Adaptive immunity is central to ICB therapy, including the targeting of program cell death protein 1 (PD1) and cytotoxic T lymphocyte-associated protein 4 (CTLA4), and therapeutic efficacy may be impacted by injury-associated immune changes. Ultimately, understanding factors and mechanisms that contribute to ICB therapy resistance, including distal perturbations such as tissue injury, can help identify biomarkers predictive of responder status and guide the design of treatments to sensitize non-responders. This is particularly important given the rapid expansion of adjuvant, neoadjuvant and peri-operative (neoadjuvant/adjuvant) ICB treatment for high-risk resectable cancers with high probability for post-surgical relapse.

Here, we demonstrate that a volumetric muscle loss (VML) injury, which consists of skin and severe muscle trauma, can accelerate progression and impair the ICB therapy response of syngeneic murine tumors. The injury-induced accelerated tumor growth coincides with decreased tumor-infiltrating effector CD8^+^ T cells and relies on a shared draining lymph node (LN) between the tumor and the injury site. Treatment of the muscle defect with a pro-regenerative biologic scaffold prevented injury-induced accelerated tumor growth in a type-2 immune-dependent manner. Collectively, we show that injury-associated immune perturbation may contribute to cancer progression and adjuvant ICB therapy resistance that can be mitigated by effectively treating the distant injury site with wound healing biomaterials.

## Results

### Distal Muscle Injury Accelerates Murine Tumor Growth

To investigate whether tissue injury can impact primary tumor growth, we established a murine model that couples bilateral VML injury, consisting of a longitudinal skin incision and critical-sized excisional defect in the quadriceps muscles, with subcutaneous inoculation of syngeneic cancer cells (CT26 colon adenocarcinoma or B16F10 melanoma, 100,000 cells/mouse) on the right flank (**Fig. 1A**). We observed that concurrent VML injury significantly accelerated murine tumor growth (**Fig. 1B**). For example, VML injury 10 days prior to tumor cell inoculation significantly increased the size of CT26 tumors relative to uninjured controls (No VML: 325.8±129.9 mm^3^, VML: 849.1±662.1 mm^3^) when measured on day 19 post-inoculation, the earliest timepoint at which survival criteria was met (tumor volume >1500 mm^3^) (**Fig. 1B**, **S1A**). Similarly, VML injury 7 days prior to tumor cell inoculation consistently resulted in accelerated CT26 tumor growth, replicated in 10 out of 12 independent experiments (**Fig. S1B**). This relative sequence aligns tumor cell inoculation with the inflammatory phase of the VML injury response that peaks around 1-week post-injury and is marked by elevated immune cell infiltration into the wound site^51,54–56^. Similarly, VML injury significantly accelerated B16F10 tumor growth compared to uninjured controls (**Fig. 1C**, **S1C**).

**Fig. 1.**
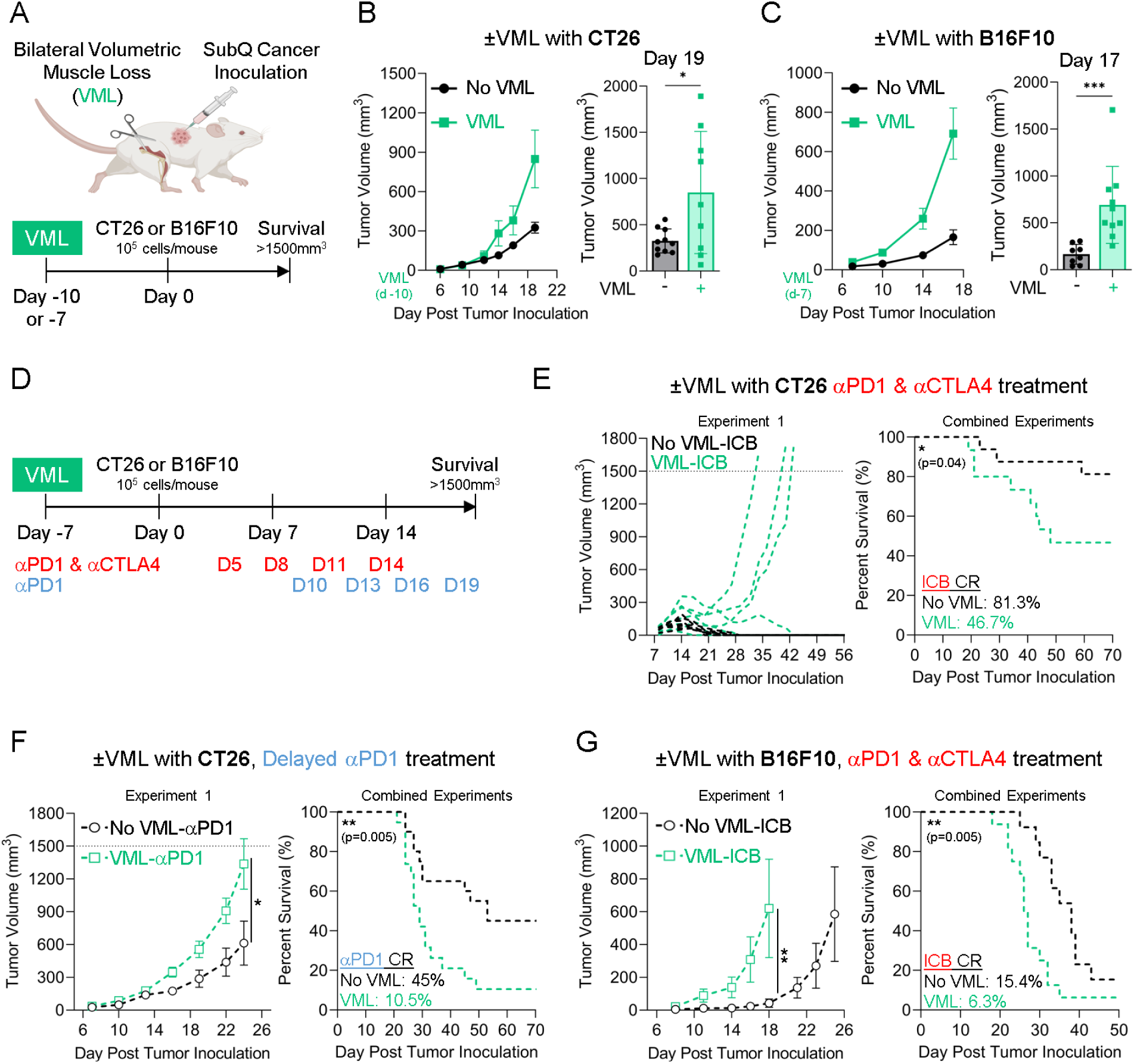
VML Injury Accelerates Murine Tumor Growth and Impairs Response to ICB Therapy. **(A)** Experimental scheme consisting of bilateral VML surgery performed on the hindlimb quadriceps muscles, followed by unilateral subcutaneous inoculation of syngeneic cancer cells (CT26 or B16F10) on the right flank. Survival criteria comprised of tumor volume >1500 mm^3^ or severe involuting ulceration. **(B)** CT26 and **(C)** B16F10 tumor growth kinetics of uninjured and VML-injured mice (n=8-10). **(D)** Experimental scheme introducing ICB therapy to the concurrent VML injury-tumor model. ICB therapy consisted of either combination αPD1/αCTLA4 (5 mg/kg each) or delayed αPD1 monotherapy (5 mg/kg) delivered via intraperitoneal injection every 3 days for 4 total doses. **(E)** CT26 tumor growth and survival curves of uninjured and VML injured mice treated with αPD1/αCTLA4 or **(F)** delayed αPD1 monotherapy. **(G)** B16F10 tumor growth and survival curves of uninjured and VML injured mice treated with αPD1/αCTLA4. **(Statistics)** Tumor growth curves: mean±SEM. Bar graphs: mean±SD (displaying earliest survival timepoint). Normally distributed data (Shapiro-Wilk test, α=0.05) was analyzed using an unpaired two-tailed student t-test (*B*); otherwise, a non-parametric two-tailed Mann-Whitney test was used (*C, F, G*). Results representative of at least 2 independent experiments (*B, C*). Survival: Kaplan-Meier curve with Log-Rank Mantel-Cox test (*E-G*). Survival results combined from 2 independent experiments with separate results presented in supplemental figure 2 (*E-G*). NS: Not significant p>0.05, * p<0.05, ** p<0.01, *** p<0.001. CR: Complete Responder.

### Muscle Injury Impairs Tumor Response to Adjuvant ICB Therapy

We extended the previously defined concurrent injury-tumor model to include ICB treatment consisting of either combination αPD1/αCTLA4 therapy (5 mg/kg each, RMP1-14 and 9H10) or delayed αPD1 monotherapy (5 mg/kg, RMP1-14) (**Fig. 1D**). Immunogenic CT26 tumors that are canonically sensitive to ICB therapy demonstrated a robust response to combination αPD1/αCTLA4 therapy in the uninjured group as expected; however, the concurrent VML injury group contained several fast-growing non-responding tumors, resulting in a decreased proportion of complete responders (No VML: 81.3%, VML: 46.7%) and significantly shortened overall survival (**Fig. 1E**, **S2A-B**). Likewise, VML injury impaired CT26 tumor response to a milder αPD1 monotherapy regimen as marked by significantly accelerated tumor growth, decreased proportion of complete responders (No VML: 45%, VML: 10.5%), shortened median survival (No VML: 53 days, VML: 29 days), and significantly decreased overall survival (**Fig. 1F**, **S2C-D**). Further, while B16F10 tumors are resistant to αPD1/αCTLA4 therapy and most ultimately reached survival criteria, the concurrent VML injury significantly accelerated the progression of ICB-treated B16F10 tumors, shortened median survival (No VML: 38 days, VML: 26.5 days), and significantly shortened overall survival (**Fig. 1G**, **S2E-F**).

### Concurrent Muscle Injury Dampens CD8^+^ T Cell-Mediated Anti-Tumor Immunity

Cytotoxic CD8^+^ T cells are potent drivers of anti-tumor immunity and serve as key targets for ICB therapy, therefore we investigated whether concurrent VML injury disrupted the effector CD8^+^ T cell response. The intra-tumoral density of CD8^+^ T cells (cell count/tumor volume) significantly decreased with concurrent VML injury, even though their proportion of total CD3^+^ tumor-infiltrating T cells did not change (**Fig. 2A**). The expression of co-inhibitory checkpoint molecules (i.e., PD1, CTLA4, LAG3, TIM3) on tumor-infiltrating CD8^+^ T cells was similar between the uninjured and VML-injured groups (**Fig. 2B**, **S3A**). Likewise, the ratio of CD8^+^ T cells to immunosuppressive FoxP3^+^ Tregs was similar between the uninjured and VML-injured groups (**Fig. S3B**).

**Fig. 2.**
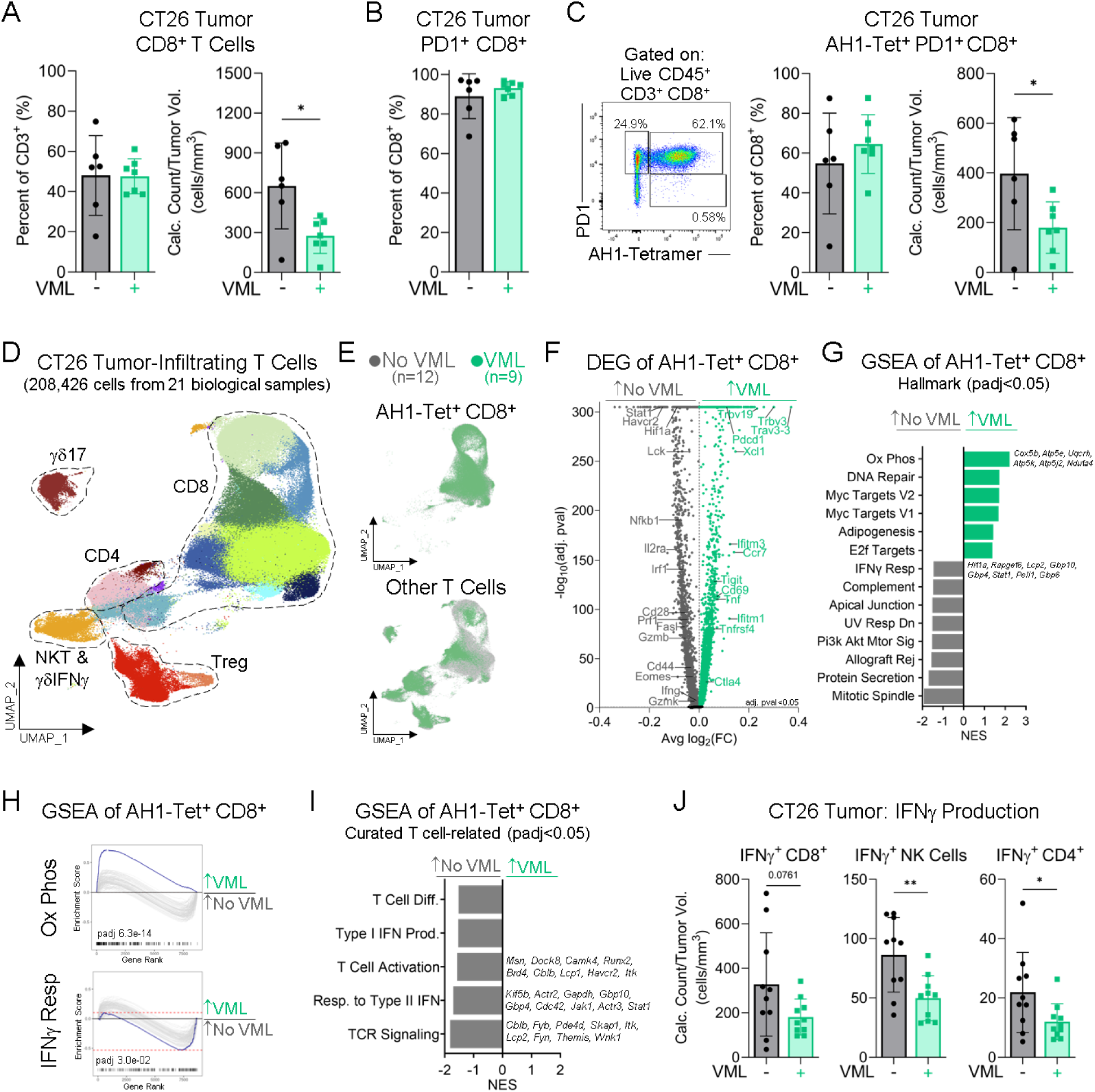
VML Injury Impacts Anti-Tumor CD8^+^ T Cell Response. **(A)** Flow cytometric profiling of CT26 tumor-infiltrating CD8^+^ T cells, **(B)** PD1 expression on CD8^+^ T cells, and **(C)** CT26 tumor-specific CD8^+^ T cells marked by AH1-loaded MHC class I tetramer in uninjured and VML injured mice. **(D)** Single-cell RNA-sequencing (scRNAseq) of CT26 tumor-infiltrating T cells from uninjured and VML injured mice. UMAP depicting main T cell clusters (e.g., CD8, CD4, Treg, unconventional). **(E)** T cell UMAP separated by tumor-reactive AH1-Tetramer^+^ CD8^+^ T cells (top) and other T cells (bottom) colored by cells from uninjured (gray) and VML injured (green) mice. **(F)** Differential gene expression and **(G, H)** GSEA using Hallmark or **(I)** curated list of T cell-related pathways with top leading edge genes comparing tumor-reactive AH1-Tetramer^+^ CD8^+^ T cells of uninjured and VML injured mice. **(J)** Intracellular cytokine staining following ex vivo stimulation of IFNγ-producing effector cell types in CT26 tumors of uninjured and VML injured mice. **(Statistics)** Bar graphs: mean±SD. Normally distributed data (Shapiro-Wilk test, α=0.05) was analyzed using an unpaired two-tailed student t-test (*A-C, J*). Results representative of at least 2 independent experiments (*A-C*). NS: Not significant p>0.05, * p<0.05, ** p<0.01. UMAP: Uniform manifold approximation and projection. GSEA: Gene set enrichment analysis.

Tumor-infiltrating T cells are a heterogeneous population composed of tumor-specific and bystander T cells. We identified and characterized the CT26 tumor-specific CD8^+^ T cells by using a class I tetramer loaded with the immunodominant tumor antigen gp70_423-431_ (AH1)^57^. The AH1-Tet^+^ CD8^+^ T cells made up a substantial portion of the CT26 tumor-infiltrating CD8^+^ T cells and overwhelmingly expressed PD1 (**Fig. 2C**). Similar to the overall CD8^+^ T cell results, concurrent VML injury did not change the proportion of AH1-Tet^+^ PD1^+^ CD8^+^ T cells, but significantly decreased their intra-tumoral density, which suggests a diminished effector response (**Fig. 2C**, **S3C**). For in-depth transcriptomic profiling, we isolated CT26 tumor-infiltrating T cells, including AH1-Tet^+^ tumor-reactive CD8^+^ T cells, using fluorescence-activated cell sorting (FACS) and performed single-cell RNA-sequencing (scRNAseq). We sequenced 208,426 high-quality cells from 21 independent biological replicates (No VML: 12, VML: 9) and effectively captured the main T cells subsets including CD8 T cells, conventional CD4 T cells, Tregs, and unconventional T cells (i.e. γδ and NKT) (**Fig. 2D-E, S4A-B**). The CD8 T cells overwhelmingly expressed co-inhibitory receptors (*Ctla4, Lag3, Pdcd1*) (**Fig. S4C**) as well as effector molecules (*Gzmb, Ifng, Prf1, Tnf*) (**Fig. S4D**) indicative of an exhausted effector phenotype. Sub-cluster analysis revealed 10 distinct CD8 T cell subsets, including multiple cytotoxic effector (*Gzmd/e/f, Nkg7, Prf1, Lgals3*), IFN-stimulated (*Isg15, Isg20, Ifit1, Bst2*), early effector (*Ccr2, Itgb7, Ly6c2, Ifitm1/2, Gzma, Ccl5*)^58,59^, naïve/memory-like (*Ccr7, Il7r, Tcf7*)^58,59^, and cycling (*Mki67, Top2a, Ube2c*) subsets (**Fig. S4E**). We confirmed that the tumor-reactive CD8^+^ T cells had increased expression of alpha and beta T cell receptor (TCR) variable regions previously reported for AH1-Tet^+^ CD8^+^ T cell clones in CT26 tumors (*Trav3-3, Trav7, Trav21-dv12, Trbv13-2, Trbv19, Trbv31*)^60–62^ in comparison to the other CD8^+^ T cells (**Fig. S4F**).

To investigate how VML injury affects anti-tumor T cell immunity, we performed differential gene expression analysis on AH1-Tet^+^ CD8^+^ T cells from uninjured and VML injured mice (**Fig. 2F**). The tumor-reactive CD8^+^ T cells from uninjured mice had higher expression of T cell activation (*Stat1, Cd28, Lck, Il2ra, Cd44*) and cytotoxic (*Prf1, Gzmb, Fasl, Ifng*) genes, while those for injured mice had increased expression of co-inhibitory receptors (*Pdcd1, Ctla4, Tigit*) and other activation genes that do not necessarily contribute to effector activity (*Cd69, Xcl1*). Accordingly, gene set enrichment analysis (GSEA) using Hallmark pathways revealed significant enrichment in IFNγ response by tumor-reactive CD8^+^ T cells from uninjured relative to injured mice (**Fig. 2G-H**), suggestive of a heightened anti-tumor effector phenotype that may contribute to the observed slower tumor growth and greater ICB therapy response. VML injury may also promote metabolic reprogramming of tumor-reactive CD8^+^ T cells as indicated by a significant shift from PI3K/AKT/mTOR signaling in uninjured mice, which promotes T cell glycolysis and activation^63,64^, to oxidative phosphorylation in VML injured mice (**Fig. 2G-H**). By using a curated group of T cell-related gene sets, we further observed that tumor-reactive CD8^+^ T cells from uninjured mice were significantly enriched in TCR signaling, T cell activation, and IFN pathways (**Fig. 2I**). Complementary intracellular cytokine staining following *ex vivo* stimulation revealed that concurrent VML injury decreased intra-tumoral density of IFNγ^+^ effector sub-populations, including CD8^+^ T cells, CD4^+^ T cells, and natural killer (NK) cells (**Fig. 2J**). While VML injury was associated with a diminished intra-tumoral CD8^+^ T cell effector response, CT26 tumor growth kinetics were still accelerated in injured mice relative to uninjured controls with systemic CD8^+^ T cell depletion (αCD8β, 5 mg/kg, 53-5.8) (**Fig. S5**), indicating that CD8^+^ T cell-independent mechanisms may also contribute to injury-induced accelerated tumor growth.

### A Shared Draining Lymph Node is Critical for Muscle Injury-Induced Accelerated Tumor Growth

T cell priming and activation within tumor-draining lymph nodes (tdLNs) is critical for controlling cancer progression and mounting potent immunotherapy responses^65–69^. Flow cytometric immunophenotyping revealed that concurrent VML injury significantly decreased CD8^+^ T cell effector memory (CD44^hi^CD62L^-^) differentiation in CT26 tdLNs (No VML: 2.59±0.56%, VML: 1.91±0.31%) and in αPD1/αCTLA4 treated B16F10 tdLNs (**Fig. 3A, S6A**). CD8^+^ T cells residing in CT26 tdLNs also significantly decreased surface expression of activation markers IL2Rα (CD25) and natural killer group 2 member D (NKG2D) with concurrent VML injury (**Fig. 3B-C**). While NKG2D is most often associated with NK cells, its expression is induced on activated murine CD8^+^ T cells where it can serve as a co-stimulatory receptor with signaling potentiating proliferation, effector function, and memory formation^70,71^. Overall, we observed a decrease in tdLN effector CD8^+^ T cell activation. However, the frequency and calculated counts of CT26 tumor-specific AH1-Tet^+^ CD8^+^ T cells within the tdLNs remained unchanged between uninjured and VML-injured mice (**Fig. S6B**).

**Fig. 3.**
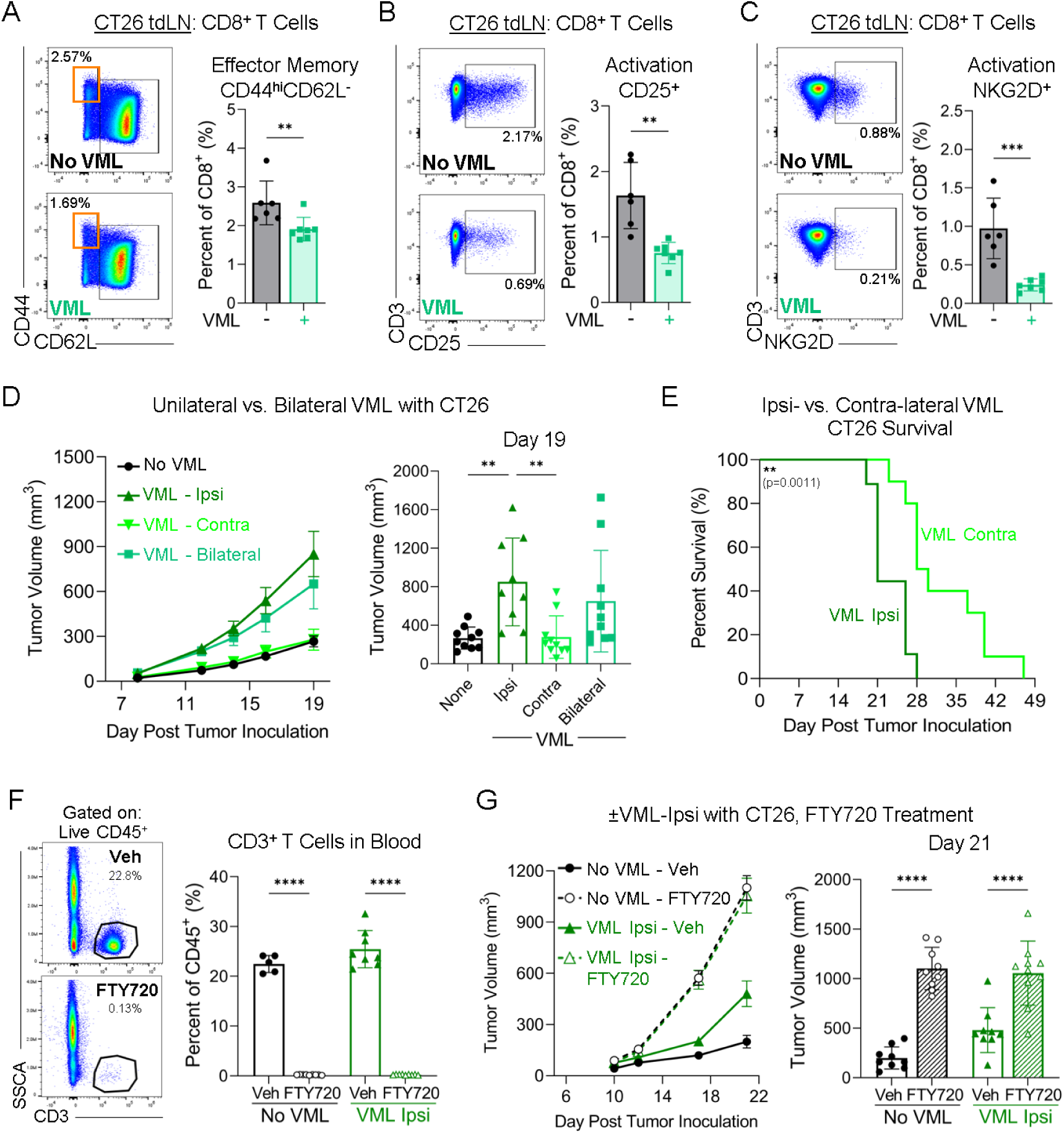
VML Injury Impacts Distant Tumor Growth by Engaging a Shared Tumor-Draining Lymph Node (tdLN). **(A)** Flow cytometric profiling of CD8^+^ T cells in CT26 tdLN, including effector memory differentiation (CD44^hi^CD62L^-^) and **(B, C)** activation (CD25^+^, NKG2D^+^) in uninjured and VML-injured mice. **(D)** The bilateral VML injury (square) was modified to a unilateral procedure performed on either the ipsilateral (triangle) or contralateral (inverted triangle) quadriceps muscle relative to the tumor-bearing flank. CT26 tumor growth kinetics of uninjured and VML-injured (bi-, ipsi-, and contra-lateral) mice (n=9-10). **(E)** Survival curve (volume >1500mm^3^ or severe involuting ulceration) of CT26 tumor-bearing mice with ipsi-versus contra-lateral VML injury. **(F)** Inhibition of lymphocyte LN egress using FTY720 HCl (25 μg/mouse). Representative flow cytometry plots and quantification of CD3^+^ T cells in peripheral blood (collected at survival endpoint) to confirm successful sequestration of lymphocytes in LN/spleen. **(G)** CT26 tumor growth kinetics of uninjured and ipsilateral VML-injured mice treated with vehicle (solid) or FTY720 HCl (dashed) (n=9-10). **(Statistics)** Tumor growth curves: mean±SEM. Bar graphs: mean±SD (*D, E* display earliest survival timepoint). For 2 groups, normally distributed data (Shapiro-Wilk test, α=0.05) was analyzed using an unpaired two-tailed student t-test (*B, C*); otherwise, a non-parametric two-tailed Mann-Whitney test was used (*A*). For >2 groups, data was analyzed using an ordinary one-way (*D*) or two-way ANOVA (*F, G*) with Tukey’s multiple comparisons test (only relevant comparisons shown). Survival: Kaplan-Meier curve with Log-Rank Mantel-Cox test (*E*). Results representative of at least 2 independent experiments (*D, G*). NS: Not significant p>0.05, * p<0.05, ** p<0.01, *** p<0.001, **** p<0.0001.

In the bilateral VML injury model, the right inguinal LN served as the primary draining LN for both the subcutaneous tumor (right flank) and muscle injury site (right hindlimb). To precisely discern whether VML injury accelerated tumor growth by modulating the tdLN, we modified our model to either an ipsilateral (right hindlimb) or contralateral (left hindlimb) VML injury relative to the right tumor-bearing flank (**Fig. S7A**). We exclusively observed accelerated CT26 tumor growth in the ipsilateral VML injury group that had a shared draining LN between the tumor and injury site; the contralateral VML injury group exhibited comparable tumor growth to the uninjured control group (**Fig. 3D**, **S7B-D**). The ipsilateral injury significantly shortened survival in comparison to contralateral injury (Median Survival, Ipsi-VML: 21 days, Contra-VML: 29 days), suggesting critical involvement of a shared draining LN (**Fig. 3E**). Additionally, we implemented an alternate skin excision injury model (dorsal 6 mm biopsy punch aligned with forelimbs) in which primary LN drainage from the skin wound (axial and brachial LN) is discrete from that of the tumor and found that the concurrent skin excision injury did not affect CT26 tumor growth or response to αPD1 monotherapy (**Fig. S8**).

To functionally test whether T cells in the shared tdLN contribute to the injury-induced accelerated tumor growth, we blocked T cell egress from LNs of uninjured and ipsilateral VML-injured mice with Fingolimod (FTY720 HCl, 25 μg/mouse), which disrupts lymphocyte expression of sphingosine 1-phosphate receptor resulting in LN sequestration (**Fig. S9A**). We confirmed successful CD3^+^ T cell depletion from peripheral circulation by flow cytometric analysis (**Fig. 3F**). Within the uninjured control cohort, FTY720 treatment significantly increased the CT26 tumor growth compared to vehicle (sterile saline) (**Fig. 3G**). Notably, FTY720 treatment abrogated the observed VML injury-induced accelerated CT26 tumor progression with comparable tumor growth kinetics in FTY720-treated uninjured and ipsilateral VML groups (**Fig. 3G**, **S9B-C**), demonstrating that LNs and T cells play an essential role in mediating communication between the tumor and muscle injury site.

### Treatment of Muscle Injury with Pro-Regenerative Biomaterial Mitigates Accelerated Tumor Growth

Since we observed that VML injury can increase primary tumor progression and impair ICB therapy response, we investigated whether treatment of the injury with pro-regenerative biologics can improve cancer outcomes. At the time of bilateral VML injury, we either implanted porcine-derived decellularized ECM scaffold (10 mg/defect) or locally administered helminth-derived regenerative soluble egg antigen solution (rSEA, 1.2 mg/mL with 75 μL/defect)^54,72^ into the defect site. Treatment of the VML injury with ECM scaffold abrogated the accelerated CT26 tumor growth with untreated VML injury, returning tumor growth kinetics to the uninjured baseline (**Fig. 4A**, **S10A**). Summary data compiled from 10 independent experiments demonstrated that mean CT26 tumor volumes of the untreated injury group were significantly greater than those of the ECM-treated injury group (**Fig. S10B**). Relative to the uninjured group, the fold change in day 14 tumor volume of the untreated injury group was significantly greater than of the ECM-treated injury group (**Fig. 4B**). Similarly, ECM treatment of VML injury significantly decreased B16F10 tumor progression compared to untreated VML injury (**Fig. 4C**, **S10C**). Further, VML injury treatment with an alternate type-2 immune stimulating pro-regenerative therapeutic, *Schistosoma mansoni* rSEA^72^, significantly reduced CT26 tumor growth kinetics compared to untreated VML injury (**Fig. 4D**, **S10D**). In the context of ICB therapy response, ECM treatment of VML injury delayed progression of CT26 tumors treated with αPD1, extending median (VML: 29 days, VML+ECM: 43 days) and overall survival (**Fig. 4E**, **S10E-F**). However, B16F10 tumor response to αPD1/αCTLA4 was not improved by ECM treatment in injured mice (**Fig. 4F**, **S10G-H**), demonstrating varied effects based on specific tumor type. Taken together, our results suggest that leveraging pro-regenerative biologics to promote tissue repair may serve as a feasible strategy for mitigating injury-induced accelerated tumor progression.

**Fig. 4.**
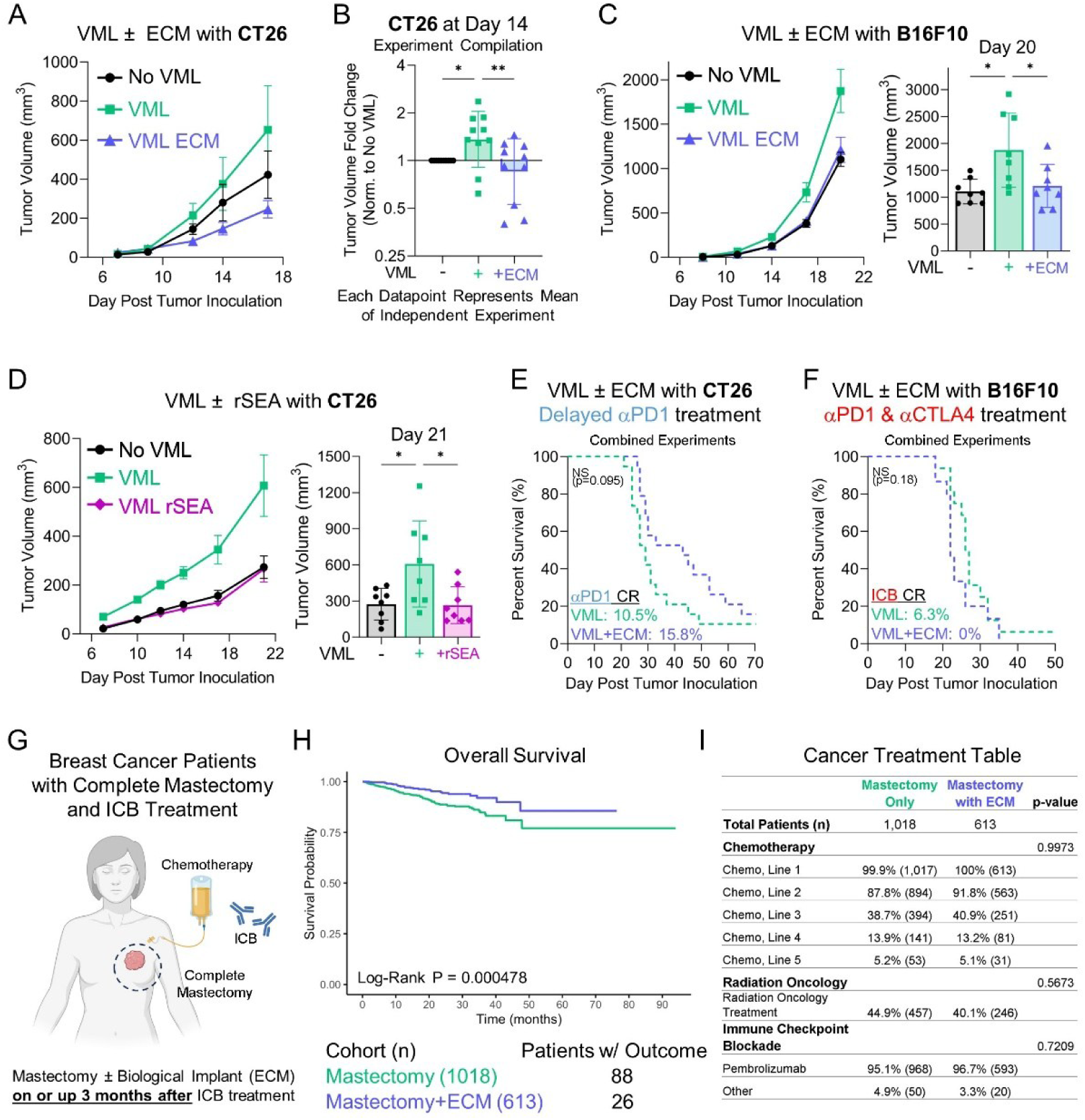
Treatment of Injury with Pro-Regenerative Biologics Impacts Cancer Progression. **(A)** CT26 tumor growth kinetics of uninjured, untreated VML-injured, and extracellular matrix (ECM) scaffold-treated VML-injured mice (n=6-8). **(B)** Experiment compilation of CT26 tumor volume fold changes (normalized to the uninjured group within each experiment) of untreated and ECM-treated VML-injured groups at day 14 post-inoculation (n=10 independent experiments). **(C)** B16F10 tumor growth kinetics of uninjured, untreated VML-injured, and ECM-treated VML-injured mice (n=8). **(D)** CT26 tumor growth kinetics of uninjured, untreated VML-injured, and *Schistosoma mansoni* regenerative soluble egg antigen (rSEA)-treated VML-injured mice (n=8). **(E)** Survival curves of untreated and ECM-treated VML-injured mice bearing CT26 tumors treated with delayed αPD1 monotherapy or **(F)** bearing B16F10 tumors treated with αPD1/αCTLA4. **(G)** Schematic of patient selection criteria for retrospective cohort study using TriNetX database. **(H)** Overall survival curve for breast cancer patients treated with ICB therapy and complete mastectomy (CPT 19303), with versus without biological implant placement (CPT 15777), on or up to 3 months after ICB therapy. **(I)** Summary table of distribution of common cancer treatments (e.g., chemotherapy, radiation) between both patient cohorts. **(Statistics)** Tumor growth curves: mean±SEM. Bar graphs: mean±SD (earliest survival timepoint or experiment endpoint). Experiment Compilation: Each datapoint represents an independent experiment, with fold change represented as geometric mean±geometric SD on a Log2 scale (*B*). Data was analyzed using an ordinary one-way ANOVA with Tukey’s multiple comparisons test (*A-D*). Survival: Kaplan-Meier curve with Log-Rank Mantel-Cox test (*E, F, I*). Survival results combined from 2 independent experiments with separate results presented in supplemental figures (*E, F*). Survival experiments were conducted in tandem with uninjured controls, thereby untreated VML data are the same as in Fig 1F and 1G (*E, F*). Categorical data in clinical cancer treatment table was analyzed using Fisher’s exact test (*I*). NS: Not significant p>0.05, * p<0.05, ** p<0.01. CR: Complete Responder.

### Biological Scaffold Placement Post Mastectomy is Associated with Improved Outcomes in ICB-Treated Breast Cancer Patients

Similar to the biomaterial used in our preclinical studies, decellularized ECM scaffolds are used clinically to fill soft tissue defects following tumor resection and during tissue reconstruction^73–79^. In the case of breast reconstruction, acellular dermal matrices (e.g., AlloDerm) provide structural support for temporary tissue expanders and permanent implants. The recent approval of Pembrolizumab (αPD1) treatment in early-stage triple-negative breast cancer (TNBC) with chemotherapy (KEYNOTE-522)^80,81^ combined with the frequent use of ECM-based scaffolds for breast reconstruction enables clinical investigation of possible synergy between the immunotherapy and reconstructive materials. We conducted a retrospective clinical cohort study using TriNetX, a multi-institutional research network of deidentified patient data, to assess whether ECM placement post mastectomy can impact overall survival of breast cancer patients treated with ICB therapy. Our patient inclusion criteria consisted of malignant neoplasm of the breast, treatment with common ICB therapies including Pembrolizumab, and complete simple mastectomy on or up to 3 months after ICB treatment (**Fig. 4G**). Patients (n=1631) were then stratified into two cohorts based on presence (n=613) or absence (n=1018) of a biological implant for soft tissue reinforcement placed on or up to 3 months after ICB therapy. Most patients were female (99.8%) and race was evenly distributed between the two cohorts (**Fig. S11**). Patient age was significantly higher in the mastectomy only cohort (Mastectomy: 55±13.8, Mastectomy+ECM: 46.5±11.7) (**Fig. S11**). In comparison to ICB-treated patients with complete mastectomies, ICB-treated patients with biological scaffold placement and complete mastectomies exhibited a significant increase in overall survival probability (Mastectomy: 77.06%, Mastectomy+ECM: 85.65%, Log-Rank P-Value=0.000478) (**Fig. 4H**). The distribution of other cancer treatments (e.g., chemotherapy, radiation, pembrolizumab) was comparable between the two cohorts (**Fig. 4I**). The occurrence and type of reconstructive procedures (e.g., free flap, temporary tissue expander, permanent synthetic breast implant) was not compared between cohorts or controlled for in this analysis.

### Biomaterial Treatment of Muscle Injury Promotes Type-2 Immune Signature in TME

Type-2 immunity is commonly associated with allergic pathologies and parasitic infections, however it also plays a critical role in wound healing and tissue repair^82^. Pro-regenerative therapies, such as decellularized ECM scaffolds and rSEA, enhance muscle repair by inducing a type-2 immune response mediated by CD4^+^ T_H_2 cells, innate lymphoid type-2 cells (ILC2), eosinophils, CD206^+^ “M2” macrophages, CD301b^+^ regenerative macrophages, and immunomodulatory dendritic cells^54,56,72,83–86^. To investigate whether ECM treatment of VML injury promotes a corresponding type-2 immune signature in the CT26 TME, we utilized an IL4-enhanced green fluorescent protein (eGFP) reporter mouse strain (4Get, C.129-Il4^tm1Lky^/J) that reports the transcriptional activity of the *Il4* locus. We found a significantly higher percentage of CT26 tumor-infiltrating CD4^+^ T cells that produced IL4-eGFP^+^ (T_H_2) in mice with ECM-treated VML injuries than in untreated injury controls (VML: 3.84±1.26%, VML+ECM: 8.41±1.78%) (**Fig. 5A**).

**Fig. 5.**
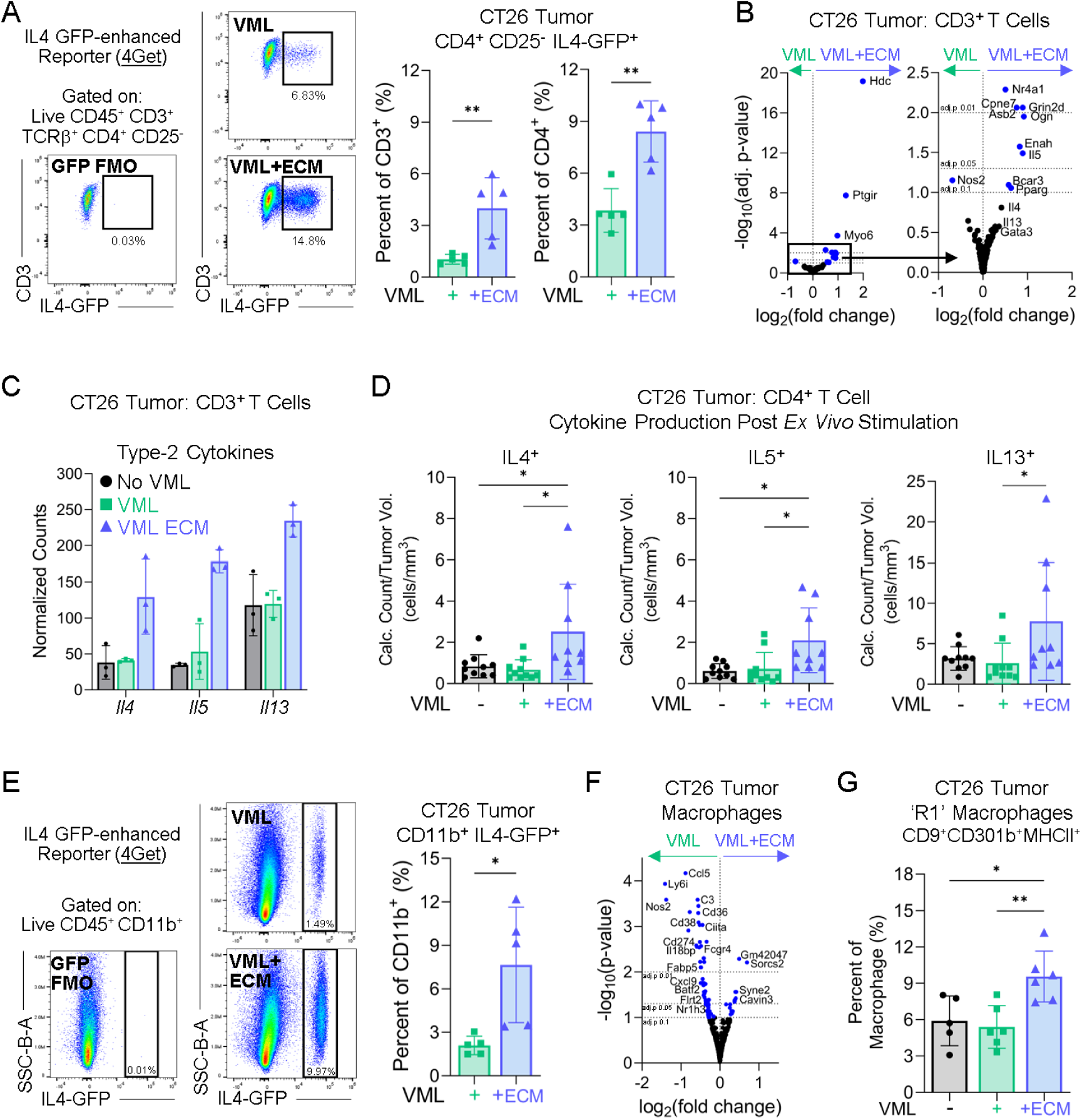
Biomaterial Injury Treatment Promotes Type-2 Immune Signature in CT26 Tumor Microenvironment. **(A)** Frequency of CT26 tumor-infiltrating IL4-eGFP^+^ CD4^+^ T cells (TH2) in VML-injured (untreated and ECM-treated) IL4-eGFP reporter (4Get) mice. CD4^+^CD25^+^ (Tregs) were removed before TH2 gating (flow plot % are out of parent CD4^+^CD25^-^, which differs from bar graph depicting % out of all CD4^+^). **(B)** Differential gene expression and **(C)** normalized RNA transcript counts per million for type-2 cytokine genes from bulk RNAseq of CT26 tumor-infiltrating CD3^+^ T cells from uninjured and VML injured (untreated and ECM-treated) mice. **(D)** Intra-tumoral density of type-2 cytokine (IL4, IL5, IL13)-producing CT26 tumor-infiltrating CD4^+^ T cells from uninjured and VML injured (untreated and ECM-treated) mice following *ex vivo* cell stimulation. **(E)** Frequency of CT26 tumor-infiltrating IL4-eGFP^+^ CD11b^+^ myeloid cells in VML injured (untreated and ECM-treated) 4Get mice. (F) Differential gene expression from bulk RNAseq of CT26 tumor-associated F480^+^ macrophages from untreated and ECM-treated VML injured mice. **(G)** Polarization of CT26 tumor-associated macrophages towards regenerative R1 (CD9^+^CD301b^+^MHCII^+^) phenotype in uninjured and VML injured (untreated and ECM-treated) mice. **(Statistics)** Bar graphs: mean±SD. For 2 groups, normally distributed data (Shapiro-Wilk test, α=0.05) was analyzed using an unpaired, two-tailed student t-test (*A, E*). For >2 groups, data was analyzed using an ordinary one-way ANOVA with Tukey’s multiple comparisons test (*D, G*). Bulk RNAseq: Negative-binomial distribution using DeSeq2 with FDR correction for multiple testing (adjusted p-value thresholds indicated on plots) (*B, F*). No statistical analysis was performed on normalized transcript counts (*C*). NS: Not significant p>0.05, * p<0.05, ** p<0.01.

ECM injury treatment increased type-2 immune gene expression in CT26 tumor-infiltrating T cells based on bulk RNAseq profiling (**Fig. 5B**). The most significantly increased gene encoded for histidine decarboxylase (*Hdc*), an enzyme involved in histamine synthesis and highly associated with type-2 mast cell allergy responses^87^. Likewise, we observed higher expression of *Abs2* (specifically expressed in T_H_2 with regulation by GATA3)^88,89^, *Il5* (a prototypical T_H_2 cytokine important for eosinophil stimulation and immunoglobulin IgE production), and *Pparg* (an anti-inflammatory factor associated with M2 macrophages and cytokine production by T_H_2)^90^. Other canonical T_H_2 marker genes (*Il4*, *Il13*, *Gata3)* were also increased with ECM treatment, albeit to a lesser extent (**Fig. 5B-C**). To assess cytokine production by tumor-infiltrating T cell at the protein level, we performed intracellular cytokine staining following *ex vivo* stimulation. IFNγ was the highest expressed of the measured cytokines by CD4^+^ T cells, but the intra-tumoral density of IFNγ^+^ (T_H_1), IL17a^+^ (T_H_17), and IL10^+^ (Treg) CD4^+^ T cells did not significantly change with ECM injury treatment (**Fig. S12A-B**). In contrast, while the proportion of type-2 cytokine (IL4, IL5, and IL13) producing CD4^+^ T cells (T_H_2) remained constant, their intra-tumoral density significantly increased with ECM treatment of VML injury (**Fig. 5D**, **S12C**).

To further analyze the impact of injury treatment on the TME, we evaluated the innate immune compartment and observed a significantly higher percentage of IL4-eGFP^+^ CD11b^+^ myeloid cells in CT26 tumors of 4Get mice with ECM-treated VML injuries compared to untreated injuries (VML: 2.09±0.64%, VML+ECM: 7.65±3.98%) (**Fig. 5E**). The proportions of various myeloid subsets (e.g., eosinophils, monocytes, and macrophages) within CT26 tumors did not significantly change with concurrent injury or injury treatment (**Fig. S13A**). Flow cytometric profiling of CT26 tumor-associated macrophages (TAMs: CD45^+^SiglecF^-^Ly6C^mid^F4/80^+^) revealed no significant differences in polarization towards classical M1 (CD86^+^CD206^-^) or alternative M2 (CD86^-^CD206^+^) phenotypes with injury treatment (**Fig. S13B**). However, complementary bulk RNAseq analysis of FACS-isolated CT26 TAMs revealed a significant downregulation of several M1 polarization-associated genes (*Nos2*, *Cd38*, *Cxcl9*, *Gbp4*, *Gbp5*)^91–95^ with ECM treatment of VML injury (**Fig. 5F**). TAMs from the untreated VML condition also had higher expression of *Ccl5* and *Cd274* (encodes programmed death ligand 1 or PDL1). Interestingly, Ccl5 has been implicated in facilitating colorectal cancer immune escape by decreasing CD8^+^ T cell infiltration and promoting PDL1 upregulation that drives T cell exhaustion with reduced cytotoxic behavior^96,97^. We extended our TAM characterization to consider fibrotic and regenerative subsets previously defined in the context of muscle injury with biomaterial implants^86^. The frequency of fibrotic F1 (CD9^-^CD301b^-^MHCII^+^, linked to type-1 inflammation) and fibrotic F2 (CD9^hi^CD301b^-^ MHCII^-^, linked to type-3 inflammation) macrophage subsets did not significantly change (**Fig. S13C**). In contrast, the regenerative R1 (CD9^+^CD301b^+^MHCII^+^) macrophage subset, defined as immune-modulatory with glycolytic metabolism and enriched with ECM implantation at the injury site^86^, was significantly increased in CT26 tumors of mice with ECM-treated VML injuries (**Fig. 5F**).

Collectively, our data suggest that features of the type-2 immune response, which develops locally with ECM implantation at the muscle injury site and promotes wound healing, also appear in the distant TME and may impact cancer outcomes.

### Tumor-Associated LN and Adipose Develop Type-2 Phenotype with Biomaterial Injury Treatment

Immunomodulation by ECM injury treatment appears to exceed the local TME and extend regionally to important tumor-associated tissues, including the tdLN and surrounding adipose. First, we observed that ECM injury treatment significantly increased the number of lymphocytes, both B220^+^ B cells and CD3^+^ T cells, in the tdLN (**Fig. 6A**). The number of Tregs (CD4^+^FoxP3^+^CD25^+^) per tdLN remained unchanged, leading to a significantly lower percentage of Tregs out of total expanded CD4^+^ T cells (**Fig. 6B**, **S14A-B**). Conversely, CD8^+^ T cell counts per tdLN increased with ECM injury treatment, resulting in a significantly higher CD8^+^ T cell to Treg ratio that may potentially contribute to the observed delay in CT26 tumor growth (**Fig. 6C**, **S14C**). We observed no differences in the frequency or number of CT26 tumor-specific AH1-Tet^+^ CD8^+^ T cells in the tdLN between injury treatment conditions (**Fig. S14D**).

**Fig. 6.**
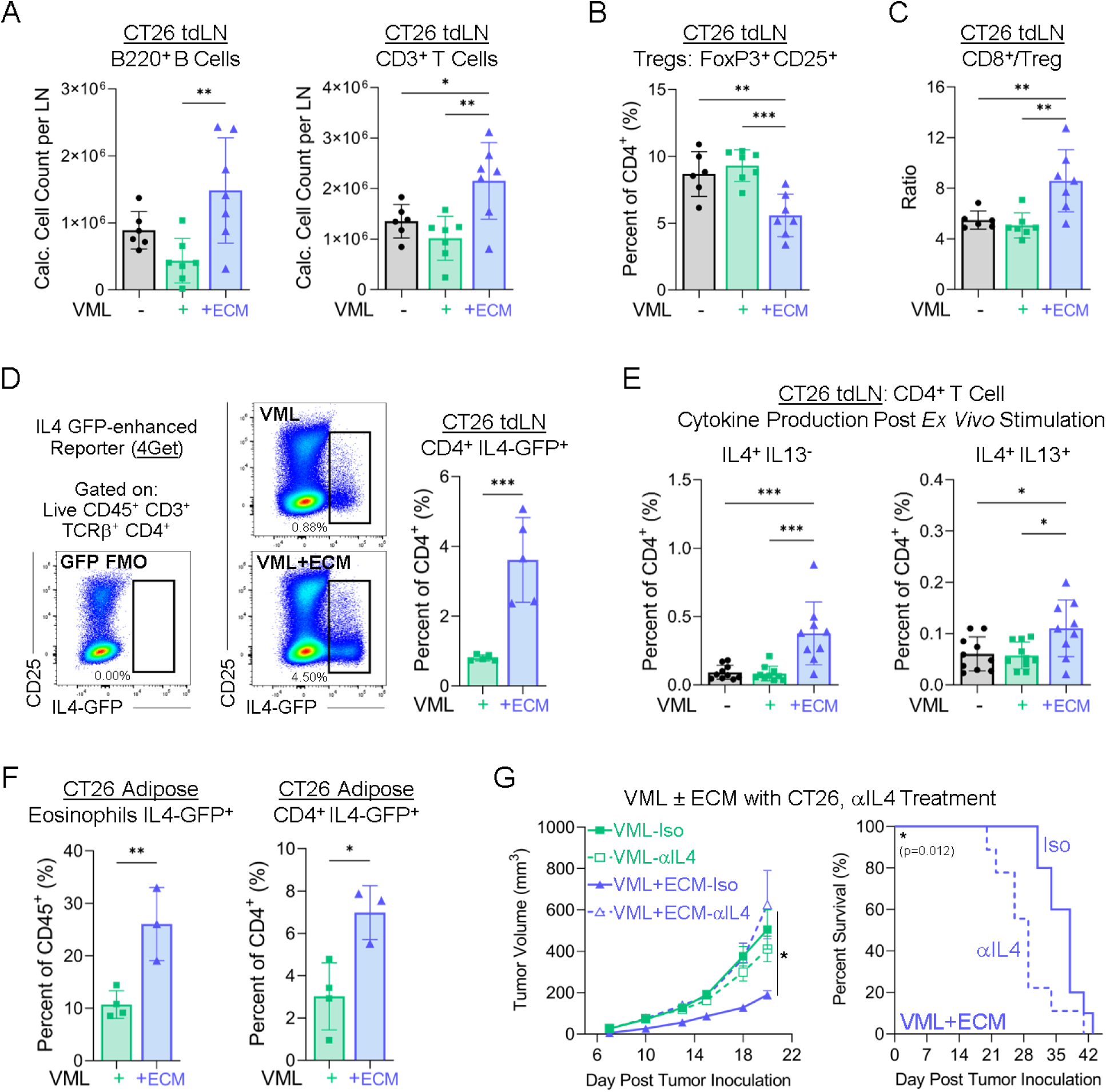
Biomaterial Injury Treatment-Induced Type-2 Immunity Also Develops in Tumor-Associated Tissues and Contributes to Delayed Tumor Growth. **(A)** Flow cytometric profiling of B and T lymphocytes, **(b)** Tregs (CD4^+^FoxP3^+^CD25^+^), and **(C)** CD8^+^/Treg ratio in CT26 tumor-draining lymph nodes (tdLNs) of uninjured and VML-injured (untreated and ECM-treated) mice. **(D)** Frequency of IL4-eGFP^+^ CD4^+^ T cells (TH2) in CT26 tdLNs of VML-injured (untreated and ECM-treated) 4Get mice. **(E)** Type-2 cytokine (IL4 and IL13) production by CD4^+^ T cells in CT26 tdLNs of uninjured and VML-injured (untreated and ECM-treated) mice following *ex vivo* cell stimulation. **(F)** Frequency of IL4-eGFP^+^ SiglecF^+^ eosinophils and CD4^+^ T cells in CT26 tumor-encapsulating adipose of VML-injured (untreated and ECM-treated) 4Get mice. **(G)** IL4 neutralization: CT26 tumor growth and survival curves of VML-injured (untreated and ECM-treated) mice treated with isotype (solid) or αIL4 antibody (1 mg/mouse initial dose followed by 0.5 mg/mouse maintenance) (dashed) (n=9-10). **(Statistics)** Tumor growth curves: mean±SEM. Bar graphs: mean±SD. For 2 groups, normally distributed data (Shapiro-Wilk test, α=0.05) was analyzed using an unpaired, two-tailed student t-test (*D, F*). For >2 groups, data was analyzed using an ordinary one-way (*A-C, E*) or two-way (*G - only D20*) ANOVA with Tukey’s multiple comparisons test. Survival: Kaplan-Meier curve with Log-Rank Mantel-Cox test (*G*). NS: Not significant p>0.05, * p<0.05, ** p<0.01, *** p<0.001.

As within the local TME, ECM injury treatment promoted type-2 immune skewing within the tdLN and surrounding adipose. Using the 4Get mouse strain, we found a significantly higher frequency of CD4^+^ T cells within the tdLN expressing IL4-eGFP^+^ (T_H_2) when the VML injury was treated with ECM scaffold compared to untreated injury controls (VML: 0.80±0.09%, VML+ECM: 3.61±1.21%) (**Fig. 6D**). Intracellular cytokine staining following *ex vivo* stimulation validated that ECM injury treatment exclusively increased type-2 cytokine production by CD4^+^ T cells within the tdLN. We observed a significant increase in the percentage of IL4 single-positive (IL4^+^IL13^-^) and IL4/IL13 double-positive (IL4^+^IL13^+^) CD4^+^ T cells with ECM injury treatment (IL4^-^IL13^+^ and IL5^+^ were unchanged) (**Fig. 6E**, **S14E-F**). The frequency of tdLN CD4^+^ T cells producing other cytokines, including IFNγ (T_H_1), IL17a (T_H_17), and IL10, was unaffected by injury treatment (**Fig. S14F**). Similarly, the inguinal fat pad that encapsulates the subcutaneously-implanted CT26 tumor and tdLN exhibited a significant increase in IL4-eGFP^+^ cells, both CD11b^+^SiglecF^+^ eosinophils and CD4^+^ T cells (T_H_2), with ECM injury treatment (**Fig. 6F**). In all, the type-2 immune phenotype critical for promoting muscle injury healing with the ECM scaffold also arises in the distant tumor and its surrounding tissues, including the tdLN and encapsulating adipose.

### Disruption of Biomaterial-Induced IL4 Signaling Impacts Tumor Growth Kinetics

Lastly, we assessed whether type-2 immune activation functionally contributes to the delayed tumor growth kinetics we observe with ECM injury treatment. We disrupted type-2 signaling by administering an IL4 neutralizing antibody (αIL4, 11B11) starting with a high dose (1 mg/mouse) two days before CT26 tumor inoculation and maintained by lower doses (0.5 mg/mouse) upon tumor inoculation and subsequently every 5 days for the duration of the study (**Fig. S15A**). Within the untreated VML cohort, IL4 neutralization did not affect CT26 tumor growth kinetics compared to isotype controls (**Fig. 6G**, **S15B-D**). In contrast, IL4 neutralization significantly accelerated CT26 tumor progression and shortened survival within the ECM-treated VML cohort (Median Survival in VML+ECM, Iso: 38 days, αIL4: 29 days) (**Fig. 6G**, **S15B-D**). Most importantly, the significant prolong survival observed with ECM treatment relative to untreated injury between the isotype treated groups (Median Survival in Isotype, VML: 29 days, VML+ECM: 38 days) was eliminated when IL4 signaling was neutralized (Median Survival in αIL4, VML: 30 days, VML+ECM: 29 days). These findings suggest that ECM treatment of concurrent muscle injury mitigates injury-induced accelerated tumor growth through, at least in part, an IL4-dependent mechanism.

## Discussion

Tissue injury (e.g., physical trauma, chronic inflammatory diseases, surgical intervention) is an extrinsic factor that can contribute to cancer progression, recurrence, and metastatic spread. While numerous preclinical injury models demonstrate this association^31,39–41,98–100^, most do not recapitulate the substantial skin and muscle trauma that accompanies tumor resections and subsequent tissue reconstructions. To address this gap, we implemented a traumatic VML injury model coupled with subcutaneous inoculation of syngeneic cancer cells. While preclinical tumor resection models are arguably more clinically relevant, their implementation specifically to study tissue injury is poorly suited; if primary tumor resection were to serve as the injury, the uninjured control group forgoing resection would be inevitably confounded by prolonged primary tumor burden. We consistently demonstrated that VML injury occurring 1-week before cancer cell inoculation accelerates CT26 and B16F10 tumor progression. In this experiment scheme, the tumor inoculation aligned with the acute immunological peak of the VML injury response. This model is particularly pertinent to a minimal residual disease setting, such as neoadjuvant or adjuvant ICB with surgical tumor resection.

Motivated by the large number of patients whose tumors unfortunately do not respond to ICB therapy for reasons still largely unknown, we extended our investigation to assess whether tissue injury can contribute to tumor ICB resistance. The relevance of tissue injury as a host factor affecting ICB therapy efficacy is further supported by the numerous approved and ongoing clinical trials coupling surgical tumor resection with (neo)adjuvant ICB therapy^101–112^. We demonstrated that concurrent VML injury can impair tumor sensitivity to adjuvant αPD1 and combination αPD1/αCTLA4 therapy, resulting in faster tumor progression, shortened survival, and decreased proportion of complete responders. Notably, we observed considerable variability in the effect of VML injury on tumor ICB therapy response (e.g., fast-growing non-responders, initial responders with subsequent tumor escape, delayed responders, immediate complete responders). These heterogenous results warrant future analyses to discern why injury drives tumor ICB resistance in some mice but not others.

While we explored ICB therapy in the adjuvant (delivered after surgery) setting, future work can apply this concurrent VML injury-tumor model in the context of neoadjuvant ICB therapy to better recapitulate emerging trends in clinical cancer care. The potential for tissue injury to affect neoadjuvant ICB therapy may support response-driven surgical de-escalation for patients whose tumors exhibit clinical complete response or major pathological response (MPR). This work could complement related clinical trials, including ones assessing whether LN dissection can be safely omitted in patients with nodal melanoma achieving MPR following neoadjuvant αPD1/αCTLA4 (PRADO extension cohort of OpACIN-neo trial, NCT02977052)^113^ or considering organ-sparing alternative treatments in patients with muscle-invasive bladder cancer following neoadjuvant ICB therapy (NCT05200988 and NCT03558087)^114,115^. Overall, understanding how tissue injury can influence cancer outcomes may help inform personalized treatment selection (e.g., resection, organ preservation, alternative non-surgical intervention, active surveillance) based on pathological response to neoadjuvant ICB therapy.

Our finding that concurrent VML injury reduces tumor response to ICB therapy indicates that we have established a useful preclinical model to elucidate, and subsequently target, mechanisms contributing to ICB resistance. Cytotoxic CD8^+^ T cells are vital effector cells in orchestrating anti-tumor immunity and serve as targets for ICB therapy. Therefore, we first investigated whether tissue injury disrupts the protective CD8^+^ T cell response. Prior studies found that surgical trauma could induce CD8^+^ T cell exhaustion and dysfunction, marked by upregulated expression of co-inhibitory checkpoint molecules, reduced effector cytokine production, and decreased proliferation that ultimately impair tumor control^116–125^. Similarly, we observed that concurrent VML injury significantly reduced the intra-tumoral density of CD8^+^ T cells, including CT26 tumor-specific CD8^+^ T cells. Additionally, scRNAseq profiling revealed that CT26 tumor-specific CD8^+^ T cells of uninjured mice are enriched in T cell activation, TCR signaling, and effector IFN pathways. These results suggest that injury may promote a more tumor-permissive TME by dampening the effector CD8^+^ T cell response within CT26 tumors.

Beyond the effector T cell response directly within the TME, we found that VML injury decreased CD8^+^ T cell activation and effector memory differentiation within the tdLN. Involvement of the tdLN proved to be indispensable for driving VML injury-induced accelerated tumor progression. We only observed faster CT26 tumor growth when the VML injury shared a primary draining LN with the tumor (i.e., bilaterial or ipsilateral VML) and it was abrogated when lymphocyte egress from LNs was pharmacologically inhibited. Further characterization of the tdLN myeloid compartment, focusing on dendritic cell tumor-antigen uptake, antigen-presentation, and co-stimulatory molecule expression is needed to discern whether tissue injury interferes with CD8^+^ T cell priming.

While our results show that VML injury alters the CD8^+^ T cell response within the local TME and tdLN, injury-induced accelerated CT26 tumor growth persisted with systemic CD8^+^ T cell depletion. This finding suggests the contribution of CD8^+^ T cell-independent factors. Previously reported alternative, but potentially synergistic, immune-related mechanisms that link tissue injury to induction of a pro-tumorigenic TME include NK cell dysfunction^126–142^, systemic inflammation mediated by neutrophils and pro-inflammatory moieties^15,41,143–159^, and immunosuppression exerted by Tregs and myeloid-derived suppressor cells (MDSCs)^31,40,160–163^. VML injury may also modulate critical TME stromal components such as cancer-associated fibroblasts and endothelial cells. Additionally, since tissue injury cascade and tumor progression are both very energy-intensive biological processes, it’s possible that competition for metabolic resources also influences cancer outcomes.

Proposed therapeutic interventions to mitigate the risk of post-surgical cancer recurrence and metastasis consist mainly of reducing postoperative systemic inflammation with nonsteroidal anti-inflammatory drugs (NSAIDs)^41,145–152^, minimizing infection risks with antibiotics^153,155,156^, and overcoming effector cell dysfunction with immunostimulatory regimens (e.g., vaccines, ICB therapy, IL2, innate agonists)^119–121,123–126,130,131^. Here, we propose an alternate strategy to abrogate injury-induced accelerated cancer progression that entailed treating the injury site with a pro-healing decellularized ECM scaffold. Previously, our group demonstrated that co-implantation of the ECM scaffold with B16F10 tumor cells delayed tumor growth and enhanced ICB therapy response^164^. Recently, researchers combined the ECM scaffold with tumor-specific peptide antigens and immune-stimulating adjuvants to formulate a cancer vaccine platform that effectively promoted murine tumor regression and induced long-lasting immunological memory^165^. In this study, we leveraged the ECM scaffold’s pro-regenerative capacities to aid in concurrent muscle injury repair. Clinically, FDA-approved decellularized ECM scaffolds are used for wound healing and, in certain contexts, to fill defects following tumor resection^73–79^; however, their application in a preclinical setting consisting of coincident tumor and injury insults has not been extensively characterized. We observed that treating the distal VML defect with an ECM scaffold consistently negated the injury-induced accelerated CT26 and B16F10 tumor progression. Additionally, CT26 tumors treated with αPD1 exhibited slower progression when an ECM scaffold was implanted into the VML defect, resulting in a small but favorable trend in prolonged survival.

The potential for ECM scaffold-mediated repair of surgery-associated tissued injury to influence cancer outcomes is supported by our retrospective analysis of ICB-treated breast cancer patients, which revealed a clinical association between biological scaffold placement post mastectomy and overall survival probability. ICB therapy is used in the peri-operative setting for TNBC (ER^-^,PR^-^,HER2^-^), which has high recurrence rates even after surgery with curative intent. Of note, since metastasis of primary breast cancers is thought to progress locally through tissue extension and tdLN, this cohort may provide a relevant human counterpart for our murine system, revealing the importance of a common LN drainage of the tissue injury and micrometastatic tumor deposits that are the source of relapse and mortality after surgery and adjuvant ICB therapy. While promising, limitations in TriNetX query design and restricted availability of critical patient information (e.g., cancer stage, precise ICB regimen, type of biological scaffold, occurrence and type of reconstructive procedure, type of synthetic breast implant) warrant comprehensive follow-up studies and ultimately prospective clinical trials to further investigate the potential synergy between biological scaffold application to surgical sites and cancer immunotherapy.

The decellularized ECM scaffold enhances muscle repair by inducing a potent type-2 immune response characterized by local infiltration and activation of T_H_2 cells, eosinophils, CD206^+^ M2 macrophages, and CD301b^+^ regenerative macrophages^54,83,85,86^. When we treated the distal VML injury with an ECM scaffold, we detected traces of the corresponding type-2 immune signature within the tumor and its associated tissues (tdLN and encapsulating adipose). We demonstrated that this ECM scaffold-induced type-2 immunity functionally contributes to the delayed CT26 tumor progression, as antibody-mediated IL4 neutralization reverted CT26 tumor kinetics to the accelerated rate observed in the untreated VML group and significantly shortened survival. Ultimately, while the primary intention of ECM scaffold implantation in a distal injury defect was to enhance wound healing, we found that the scaffold-associated type-2 inflammation may also influence anti-tumor immunity.

Type-2 immunity is predominantly regarded as pro-tumorigenic and often associated with unfavorable cancer outcomes; however, contrary evidence of it serving anti-tumorigenic roles is also reported^166–169^. For example, exogenous administration of IL4 and genetically engineered IL4-secreting tumor cell vaccines demonstrate that IL4 can effectively orchestrate tumor rejection^164,170–179^. The anti-tumorigenic effect of IL4 is closely associated with the induction of cytotoxic degranulation by eosinophils and neutrophils^180–182^, impediment of tumor neovascularization^183^, activation of NK cells^184–186^, and induction of immunological memory that is protective against subsequent tumor re-challenge^176,187^. Further, researchers recently showed that follicular helper T cells in tdLNs respond to αPD1 therapy by producing IL4, which was crucial in promoting proliferation and activation of tumor-specific CD8^+^ T cells^188^. In addition to IL4, emerging studies reveal that tumor-associated eosinophils and ILC2s can impede cancer progression and favorably contribute to ICB therapy response^189–197^. While early clinical trials of recombinant human IL4 showed only modest efficacy and undesired toxicity^198–202^, a new generation of targeted IL4-based therapies including 4-toxin conjugates, inducible IL4-expressing oncolytic adenoviruses, and tumor-targeting IL4 “immunocytokine” fusion proteins show promising results in preclinical, and in certain cases, early clinical, studies^203–209^.

Biologic scaffolds and accompanying type-2 inflammation may be leveraged to simultaneously promote tissue injury healing and inhibit cancer progression. Excitingly, recent studies show the feasibility and benefits of multi-functional biomaterial constructs engineered to deliver anti-cancer drugs (e.g., chemotherapeutics), stimulate resection wound repair, and even minimize the risk of postoperative infection^210–215^. Overall, while tissue injury may adversely impact cancer outcomes, tremendous potential exists in combining elements of regenerative medicine, biomaterial design, and cancer immunotherapy to develop novel strategies for improving postoperative cancer management.

## Materials and Methods

### Mice

All mice were housed at Johns Hopkins University animal facilities in compliance with Animal Care and Use Committee (ACUC) ethical guidelines. All procedures were performed in accordance with protocol MO21M80/MO24M66 approved by the Johns Hopkins University ACUC. Wildtype C57BL/6 and BALB/c mice were obtained from The Jackson Laboratory (strain # 000664 and 000651, respectively). IL-4/GFP-enhanced transcript (4Get) mice (C.129-Il4^tm1Lky^/J) were originally obtained from The Jackson Laboratory (strain # 004190) and subsequently bred in-house. All experiments were performed on female mice at 7-9 weeks of age.

### Murine Tissue Injury Models

#### Volumetric Muscle Loss (VML)

All surgical implements were sterilized and surfaces thoroughly disinfected. Mice were anesthetized via isoflurane inhalation, their flank and hindlimb fur was shaved, and the surgical site thoroughly disinfected. VML injury was performed as previously described^54,55,72^. In short, a 1.5cm longitudinal skin incision was made above the visible patellar ligament, followed by cutting of the overlaying fascia. A critical-sized tissue defect (3mm x 3mm x 3mm) was excised in the quadriceps muscles using sterile surgical scissors. The defect space was directly filled with either 50μL of sterile Dulbecco’s phosphate-buffered saline (dPBS, untreated groups), 50μL of ECM (preparation described below) or 75μL of rSEA (preparation described below). Immediately after treatment of defect site, the skin wound was closed using sterile veterinary nylon suture. All mice received a requisite dose (0.1mg/mouse) of non-steroidal anti-inflammatory drug (NSAID) carprofen (Rimadyl 54771-8507-1 or OstiFen 510510) administered via subcutaneous injection for pain relief. The VML injury was performed bilaterally for most experiments, except where unilateral injury (either ipsilateral or contralateral to tumor-bearing flank) is explicitly stated.

#### Skin Excision via Biopsy Punch

All surgical implements were sterilized and surfaces thoroughly disinfected. Mice were anesthetized via isoflurane inhalation, their dorsal and flank fur was shaved, and their surgical site was thoroughly disinfected. A sterile 6mm biopsy punch was used to outline a circular wound on the dorsal skin (towards the cranial end in alignment with the forelimbs over the latissimus dorsi muscle) that was fully excised using sterile serrated forceps and surgical scissors. Post-operative care and monitoring was performed as detailed above for the VML procedure.

### Preparation of Biologically-Derived Wound Healing Therapeutics

#### Porcine-Derived Decellularized Extracellular Matrix (ECM)

Multiple sources of decellularized ECM were utilized throughout the experimental studies as dictated by reagent availability. Decellularized ECM derived from porcine small intestine submucosa (SIS) was prepared in-house using a process previously outlined^164,216^. In short, fresh porcine small intestines were acquired from Wagner Meats (Mount Airy, Maryland), thoroughly washed with water, bisected longitudinally to form a flat sheet, and mechanically scraped (remove the mucosa, serosa, and muscularis externa layers). The remaining submucosal tissue was rinsed with Milli-Q water, transferred into clean autoclave-sterilized glass flasks or pyrogen-free plastics, and treated with 0.1% (w/v) peracetic acid and 4% (v/v) ethanol prepared in Milli-Q water for 2 hours with strong agitation via stir plate. The remaining SIS-ECM sheets were transferred into clean autoclave-sterilized metal sieves, extensively rinsed with Milli-Q water and sterile 1x dPBS until reaching neutral pH, manually strained to remove excess liquid, and flash frozen with liquid nitrogen prior to short-term storage at −80°C. The SIS-ECM sheets were lyophilized for approximately 3 days and then ground into particulate using a cryogenic mill. SIS-ECM particulate was stored at −20°C and UV sterilized prior to use. Alternatively, sterilized porcine urinary bladder matrix (UBM) particulate, both MicroMatrix Standard Particulate (MM1000) and MicroMatrix Fine Particulate (MM0200F), were obtained as lyophilized powders from ACell Inc. (Columbia Maryland, Integra LifeSciences Parent Organization) where they were produced with adherence to good manufacturing practices (GMP) approved for pre-clinical application. For both SIS and UBM, the ECM particulate was reconstituted in sterile 1x dPBS at 200mg/mL, loaded into a 1mL syringe (needle removed), and 50μL delivered directly into the VML defect site (10mg/defect site of ECM) at the time of surgery.

#### Helminth-Derived Regenerative Soluble Egg Antigen (rSEA)

All SEA isolation and rSEA formulation was performed in-house as previously outlined^72,216^. In short, *Schistosoma mansoni* (*S. mansoni*), reagents were provided by the NIAID Schistosomiasis Resource Center of the Biomedical Research Institute (Rockville, Maryland) facilitated by NIH-NIAID Contract HHSN272201700014I. After thawing on ice, the received eggs were re-suspended with sterile 1x dPBS (100,000 eggs/mL) and homogenized via motorized pestle or 2mL Dounce tissue homogenizer (Kimble, USA) with successful disruption visually assessed with a phase contrast microscope. The homogenized mixture was centrifuged (21,000g for 45 minutes at 4°C) and then the resulting supernatant was ultracentrifuged (100,000g for 90 minutes at 4°C). The insoluble lipid fraction (top layer) and top-half portion of the soluble fraction were sterile filtered. The two fractions were combined (9 parts of soluble fraction with 1 part of lipid fraction) prior to storage at −80°C. Protein concentration was measured via Qubit Protein Assay Kit (Invitrogen Q33211). For *in vivo* wound-healing application, rSEA protein concentration was determined to be 1.2mg/mL with 75μL delivered directly into the VML defect site (90μg/defect site of rSEA protein) at the time of surgery.

### Murine Syngeneic Cancer Models

#### In Vitro Cell Culture

Syngeneic murine cancer cell lines B16-F10 (CRL-6475) melanoma and CT26 (CRL-2638) colorectal carcinoma were obtained from the American Type Culture Collection (ATCC). B16-F10 cells were cultured in DMEM media (Gibco 11965-092) while CT26 cells were cultured in RPMI-1640 media (Gibco 11875-093). All cell culture media was supplemented with 10% (v/v) heat-inactivated fetal bovine serum (Gibco 26140-079), 100 units/mL penicillin, and 100 μg/mL streptomycin (Gibco 15140122) and vacuum-filtered (0.22μm pore size). All cell lines were handled using sterile cell culture technique and incubated at 37°C with 5% CO_2_.

#### Subcutaneous Tumor Inoculation and Monitoring

Syngeneic cancer cells were subcutaneously inoculated (B16-F10 in C57BL/6, CT26 in BALB/c) either prior to (3 or 14 days) or post (3, 7 or 10 days) tissue injury (VML ± ECM/rSEA or skin excision biopsy). Mice were anesthetized via isoflurane inhalation. Their right flanks were shaved, disinfected with 70% ethanol, and injected with 1×10^5^ cancer cells re-suspended in sterile 1x PBS using a 26-gauge syringe. External tumor dimensions were measured every 2-4 days (starting day 7 post-injection) using digital calipers. Tumor volume was calculated using the ellipsoid volume equation, volume=(π/6)*(L)*(W^2^), where L is tumor length (larger dimension) and W is tumor width (smaller dimension). Survival criteria consisted of tumor volume >1500mm^3^ (with no dimension exceeding 2cm), severe involuting ulceration, or impediment of essential bodily function (i.e., eating, urination, defecation, and mobility) as approved by JHU-ACUC (MO21M80/MO24M66). Upon meeting survival criteria, mice were euthanized by CO_2_ or isoflurane overdose asphyxiation followed by secondary cervical dislocation.

#### Statistical Analysis

All tumor growth curves are presented as mean±SEM. At the earliest timepoint at which survival criteria was met or pre-determined experimental endpoint, tumor volumes are presented using bar graphs depicting mean±SD. Pre-determined exclusion criteria included unexplained pre-mature death (with low tumor burden) or lack of palpable tumor development throughout experiment’s duration. Significant outliers as determined by ROUT statistical test (Q=2%) were excluded from analysis. If data was normally distributed (Shapiro-Wilk test, α=0.05), it was analyzed using an unpaired, two-tailed student t-test; otherwise, it was analyzed using a non-parametric two-tailed Mann-Whitney test. For experiments with more than two groups, data was analyzed using an ordinary one-way or two-way ANOVA with Tukey’s multiple comparisons test. For experiment compilation (tumor volume at day 14 post-tumor inoculation), the individual biological replicates within a given independent experiment were averaged and each data point represents the group’s experimental mean. Compiled data was analyzed using either a ratio-paired two-tailed T test or mixed-effects analysis. Survival criteria consisted of tumor volume >1500 mm^3^ (with no dimension exceeding 2 cm), severe involuting ulceration, or impediment of essential bodily function as approved by JHU-ACUC (MO21M80/MO24M66). Survival data are presented using Kaplan-Meier curves and analyzed using Log-Rank Mantel Cox test. Complete responder (CR) criteria consisted of a tumor-bearing mouse that exhibited complete and sustained tumor regression (no palpable or measurable tumor) following ICB therapy. Survival curves from two independent experiment replicates were combined for presentation in the main figures, with the individual experiments presented separately in supplement figures. All statistical analysis was performed using GraphPad Prism software (v 9-10). Not Significant (NS) p>0.05, * p<0.05, ** p<0.01, *** p<0.001, **** p<0.0001.

### *In Vivo* Antibody Treatment Regimens

#### Immune Checkpoint Blockade (ICB) Therapy

ICB therapy consisted of monoclonal antibodies blocking PD1 (clone RPM1-14, BioXCell BP0146) and CTLA4 (clone 9H10, BioXCell BP0131) delivered via intraperitoneal (IP) injection at 5mg/kg body weight each prepared in sterile dilution buffer (*InVivo*Pure pH 7.0, BioXCell IP0070). The corresponding isotype controls were Rat IgG2a (clone 2A3, BioXCell BP0089) and Polyclonal Syrian Hamster IgG (BioXCell BP0087), respectively. Early αPD1/αCTLA4 combination therapy dosing began 5 days post-tumor inoculation (CT26 and B16F10). Delayed αPD1 monotherapy dosing began 10 days post-tumor inoculation (CT26). For all regimens, ICB antibodies were delivered every 3 days for a total of 4 treatment doses.

#### CD8 T Cell Depletion

CD8^+^ T cells were depleted using αCD8β antibody (clone 53-5.8, BioXCell BE0223) delivered via IP injections at 5mg/kg body weight prepared in sterile dilution buffer (*InVivo*Pure pH 7.0, BioXCell IP0070). The corresponding isotype control was Rat IgG1 anti-horseradish peroxidase (clone HRPN, BioXCell BE0088). Mice were randomly split between injured (bilateral VML) and uninjured control groups with subcutaneous CT26 tumor cell inoculation 7 days post-injury. Antibody treatment for CD8^+^ T cells depletion began 6 days prior to tumor cell inoculation (1-day post VML injury) and dosing continued every 3 days for the entire duration of the survival experiment (10 total doses). Systemic depletion of CD8^+^ T cells was confirmed via flow cytometric (Cytek Aurora) profiling of peripheral blood using αCD45 (clone 30-F11), αCD3 (clone 17A2) and αCD8α (clone 53-6.7) fluorophore-conjugated antibodies upon meeting survival criteria.

#### IL4 Neutralization

IL4 was neutralized using αIL4 antibody (clone 11B11, BioXCell BE0045) delivered via IP injection prepared in sterile dilution buffer (*InVivo*Pure pH 7.0, BioXCell IP0070). The corresponding isotype control was Rat IgG1 anti-horseradish peroxidase (clone HRPN, BioXCell BE0088). Mice were randomly split between untreated and ECM (10mg/defect, MicroMatrix, ACell MM1000) treated bilateral VML injury groups with subcutaneous CT26 tumor cell inoculation 7 days post-injury. Antibody treatment for IL4 neutralization began with an initial high dose (1mg/mouse) 2 days prior to tumor cell inoculation (5-days post VML injury), followed by a lower maintenance dose (0.5mg/mouse) starting on the day of tumor cell inoculation (7-days post VML injury) and continued every 5 days for the entire duration of the survival experiment (9 total doses).

### Lymphocyte Egress Inhibition with Fingolimod Hydrochloride

Lymphocyte lymph node egress was inhibited using fingolimod hydrochloride (FTY720 HCl, MW 343.9, Selleckchem S5002) delivered via IP injection at 25μg/mouse prepared in sterile saline (Sodium Chloride 0.9%, Intermountain Life Sciences Z1376). The corresponding vehicle control was sterile saline. Mice were randomized between injured (ipsilateral VML) and uninjured control groups with subcutaneous CT26 tumor cell inoculation 7 days post-injury. FTY720 treatment began 2 days prior to tumor cell inoculation (5-days post VML injury) and dosing continued every 2-3 days (3 doses per week) for the entire duration of the survival experiment (15 total doses). Lymph node lymphocyte sequestration was confirmed via flow cytometric (Cytek Aurora) profiling of peripheral blood using αCD45 (clone 30-F11) and αCD3 (clone 17A2) fluorophore-conjugated antibodies upon meeting survival criteria.

### Murine Tissue Processing for Flow Cytometry

#### Tumor

Following euthanasia, subcutaneous tumors were excised with careful attention not to include surrounding inguinal adipose, lymph node or skin tissue. When possible, harvested tumors were weighed to collect tissue mass measurements. Tumors were mechanically dissociated via manual dicing followed by enzymatic digestion with Liberase TL (1.67 Wünsh U/mL, 0.5mg/mL of stock, Roche 05401020001) and DNAse I (0.2mg/mL, Roche 10104159001) in RMPI-1640 with L-Glutamine (Gibco 11875-093) and 25mM HEPES Buffer (Quality Biological 118-089-721) in an incubator shaker at 37°C for either 25 (B16-F10) or 45 (CT26) minutes. Enzymatic digestion was quenched by placing samples on ice and adding RMP1-1640 media supplemented with 1% (w/v) bovine serum albumin (Sigma-Aldrich A9647-100g). Digested samples were filtered through 70μm cell strainers (Falcon 352350) followed by either 40μm cell strainers (Falcon 352340) or a Percoll density gradient (layers of 80-40-20% Percoll) (GE Healthcare 17-0891-01) to remove necrotic cells and debris allowing for immune cell enrichment. Following centrifugation of the density gradient (2100g for 30 minutes at 37°C, acceleration and deceleration ramp set to 0), the interfacial layer between the 80% and 40% Percoll solutions was collected for flow cytometric analysis. Samples were plated into 96-well U-bottom plates and pelleted via centrifugation.

#### Tumor-Draining Lymph Node (LN)

Following euthanasia, the tumor-draining inguinal LNs were carefully separated from encapsulating adipose tissue. The harvested LNs were manually mashed through 70μm cell strainers (MACS SmartStrainers, Miltenyi Biotec 130-110-916) with blunt pestles and the strainers thoroughly rinsed with sterile dPBS.

#### Inguinal Fat Pad

The inguinal fat pad overlays the subcutaneous tumor and encapsulates the tumor-draining inguinal LN. Following euthanasia and removal of inguinal LNs, inguinal fat pads were harvested and processed using the same protocols as outlined above for tumor samples including mechanical dissociation, enzymatic digestion (20 minutes), and filtering through 70μm cell strainers.

#### Blood

Peripheral blood was collected into K2EDTA-coated tubes (BD 365974) using cheek-bleed approach in isoflurane anesthetized mice. Blood volume for analysis was standardized across all samples (typically 150μL/sample) and red blood cell lysis was performed with BD Pharm Lyse Lysing Buffer (BD 555899).

### Multi-Parameter Flow Cytometry

Harvested tissues were processed into single-cell suspensions as detailed above and plated into 96-well U-bottom plates in which all subsequent staining steps took place. Multiple flow cytometry panels were developed to achieve both breadth and depth in immune phenotyping including T cell checkpoint (**Table S1**), T cell effector memory (**Table S2**), T cell intracellular cytokine (**Table S3**), 4get (**Table S4**), T cell tetramer (**Table S5**), and macrophage phenotype (**Table S6**) panels. The FACS buffer used for all wash and staining steps (except viability) consisted of dPBS without CaCl_2_ or MgCl_2_ (Gibco 14190-144) supplemented with 1% (w/v) bovine serum albumin (Sigma-Aldrich A9647-100g) and 1mM EDTA (Invitrogen 15575-038).

#### Viability Staining

Wells were washed with dPBS without CaCl_2_ or MgCl_2_ (Gibco 14190-144) to remove residual media proteins prior to viability staining. Zombie NIR Fixable Viability Stain (BioLegend 423106) was used as the amine-reactive viability stain for all panels, except macrophage phenotype panel which used Fixable Viability Dye eFluor780 (eBioscience 65-0865-14). Samples were incubated in 100μL of viability dye for 30 minutes on ice in the dark. Following viability staining, samples were washed with FACS buffer at least twice.

#### Surface Marker Staining

All surface marker antibodies used were directly conjugated to fluorophores. All antibody stains were prepared in FACS buffer supplemented with TruStain FcX anti-mouse CD16/CD32 antibody (1:20, clone 93, BioLegend 101320), True-Stain Monocyte Blocker (1:50, BioLegend 426102), and Super Bright Complete Staining Buffer (1:50, eBioscience SB-4401-75). Antibodies conjugated to certain protein-based tandem dyes (i.e., PE-Cy5, PE-Cy7, APC-Cy7, APC-Fire750, and APC-eFluor780) were added to antibody cocktail immediately prior to staining to minimize tandem dye aggregation. Samples were incubated in 100μL of antibody stain for 45 minutes on ice in the dark. Following surface staining, samples were washed with FACS buffer at least twice. If no further intracellular staining was being performed, samples were fixed using FluoroFix buffer (BioLegend 422101) for 15 minutes at room temperature in the dark (macrophage phenotype panel was not fixed). Samples were washed with FACS buffer at least twice prior to running on cytometer.

#### CD8 T Cell Tetramer Staining

Class I MHC tetramers were utilized for identification and profiling of tumor-specific CD8^+^ T cells. The H2-Ld MuLV gp70 env_423-431_ tetramer (1:100, Peptide Sequence SPSYVYHQF referred to as AH1, NIH Tetramer Core Facility) conjugated to APC was used for CT26 tumor experiments (**Table S5, S7**). Irrelevant peptide-loaded tetramers and lymph nodes from non-tumor bearing mice served as negative controls. All tetramer staining was performed following viability staining and prior to surface marker staining. Tetramer stains were prepared in FACS buffer supplemented with TruStain FcX and True-Stain Monocyte Blocker. Samples were incubated in 100μL of tetramer stain for 30 minutes at room temperature in the dark. Following tetramer staining, samples were washed with FACS buffer at least twice prior to proceeding with surface marker staining as detailed above.

#### Nuclear Transcription Factor Staining

Samples underwent viability and surface marker staining as detailed above, followed by fixation and permeabilization using the True-Nuclear Transcription Factor buffer set (BioLegend 424401). Samples were incubated in 1x fixative for 60 minutes at room temperature in the dark, followed by three rounds of washing with 1x Perm buffer. Intracellular antibody stains (**Table S1, S2**) were prepared in 1x Perm buffer supplemented with TruStain FcX, True-Stain Monocyte Blocker, and Super Bright Complete Staining Buffer. Samples were incubated in 100μL of antibody stain for 45 minutes at room temperature in the dark. Following intracellular staining, samples were washed with 1x Perm buffer three times prior to storage in FACS buffer.

#### Ex Vivo T Cell Stimulation for Cytokine Production

*Ex vivo* cell stimulation and inhibition of protein transport was performed to capture T cell phenotype as assessed by intracellular cytokine production. Tumor (following Percoll density gradient) and LN samples were incubated in 200μL of 1x Cell Stimulation Cocktail plus protein transport inhibitors (contain phorbol 12-myristate 13-acetate (PMA), ionomycin, brefeldin A, and monensin) (500x stock, eBioscience 00-4975-93) diluted in Iscove’s Modified Dulbecco’s Media (IMDM) with L-Glutamine and 25mM HEPES without phenol red (Gibco 21056023) supplemented with 10% (v/v) heat-inactivated fetal bovine serum (Gibco 26140-079) for 4 hours at 37°C with 5% CO_2_.

#### Intracellular T Cell Cytokine Staining

Samples underwent *ex vivo* cell stimulation with protein transport inhibition, viability staining, and surface marker staining as detailed above followed by fixation and permeabilization using the Cyto-Fast Fix/Perm buffer set (BioLegend 426803). Samples were incubated in 100μL of Cyto-Fast Fix/Perm solution for 20 minutes at room temperature in the dark, followed by two rounds of washing with 1x Cyto-Fast Perm Wash solution. Intracellular cytokine antibody stains (**Table S3**) were prepared in 1x Cyto-Fast Perm Wash solution supplemented with TruStain FcX, True-Stain Monocyte Blocker, and Super Bright Complete Staining Buffer. In addition to single stains and FMOs, isotype controls were prepared for all intracellular antibodies. Samples were incubated in 100μL of antibody stain for 20 minutes at room temperature in the dark. Following intracellular staining, samples were washed with 1x Cyto-Fast Perm Wash solution at least twice prior to storage in FACS buffer.

#### Flow Cytometer and Analysis Software

All panels were run on the spectral 4-laser Cytek Aurora flow cytometer (Lasers: 405nm Violet, 488nm Blue, 561nm Yellow/Green, and 640nm Red) with automated sample loader except for the macrophage phenotype panel which was run on the conventional 4-laser Attune NxT flow cytometer (Lasers: 405nm Violet, 488nm Blue, 532nm Green, and 637nm Red). Cytek Aurora data acquisition and subsequent spectral unmixing was performed using SpectroFlow software.

Autofluorescence signatures were extracted from unstained samples to remove background signal and improve resolution. Attune NxT data acquisition was performed using the Attune Cytometric Software. Data analysis was performed using FlowJo software (Tree Star, Version 10.8). Manual gate placement for cell population delineation was guided by FMO and isotype controls. Estimated total cell counts were calculated using the following formula: EstTotalCount=(TotalVol/AcqVol)*(DilFac)*(Count), where TotalVol was total sample volume, AcqVol was sample volume acquired as recorded by the cytometer’s volumetric measurement, DilFac was the dilution factor by which the sample was split prior to staining, and Count was the number of events as recorded by cytometer within the desired cell population gate. For tumor samples, the estimated total cell count was normalized to tumor volume to report cell density.

#### Statistical Analysis

Flow cytometry results were presented as either proportions (percentage of larger parent population), calculated cell counts, or density (calculated cell counts/tumor volume) using bar graphs depicting mean±SD. If data was normally distributed (Shapiro-Wilk test, α=0.05), it was analyzed using an unpaired, two-tailed student t-test; otherwise, it was analyzed using a non-parametric two-tailed Mann-Whitney test. For experiments with more than two groups, data was analyzed using an ordinary one-way ANOVA or two-way ANOVA with Tukey’s multiple comparisons test. All statistical analysis was performed using GraphPad Prism software (v 9-10). Not Significant (NS) p>0.05, * p<0.05, ** p<0.01, *** p<0.001, **** p<0.0001.

### Single-Cell RNA-Sequencing

#### Sample Preparation

The transcriptomic phenotypes of CT26 tumor-infiltrating T cells were profiled using single-cell RNA-sequencing (scRNAseq). Mice were randomized between uninjured and VML injured groups with subcutaneous CT26 tumor inoculation 7 days post-injury. CT26 tumors (n=12/group) were harvested 16 days post tumor inoculation and processed using the flow cytometry protocol detailed above including a Percoll density gradient. Viability, CD8 class I tetramer, and surface marker staining were performed as detailed above (**Table S7**). During the last 30 minutes of surface staining, a unique TotalSeq-C hashtag oligo (HTO) antibody was added to the staining mixture for each sample within the group (**Table S7**). Prior to sorting, cells from 4 tumor samples within a group (each marked with a unique HTOs) were combined, resulting in 3 hashed-pooled samples per group. Cell sorting was performed by the JHU SKCCC High Parameter Flow Core on a 4-laser BD FACSAria Fusion Cell Sorter. CT26 tumor-specific CD8 T cells (SSC^lo^, Singlets, Live, CD45^+^, CD11b^-^, CD3^+^, CD8^+^, AH1-Tetramer^+^) and all other T cells (SSC^lo^, Singlets, Live, CD45^+^, CD11b^-^, CD3^+^, AH1-Tetramer^-^) were sorted separately into dPBS with 1% BSA, centrifuged, and re-suspended in dPBS with 0.01% BSA. Sorted T cells were counted using the Countess III (Thermo Fisher Scientific) and processed for scRNAseq with HTOs by Psomagen. A targeted 20,000 cells per sample were loaded to a single Gel Bead-in-Emulsion (GEM) reaction onto the Chromium X instrument (10X Genomics). Single-cell RNA-seq and feature barcode libraries were prepared using Chromium GEM-X Single Cell 5’ Reagent kits V3 (CG000734) according to manufacturer protocols. Libraries were profiled by Tapestaion High Sensitivity DNA kit (Agilent Technologies) and quantified. Libraries were sequenced on NovaSeq X Plus at 50,000 reads/cell for GEX library and 5,000 reads/cell for feature barcode library.

#### Single-Cell RNAseq Data Analysis

FASTQ files were aligned with 10X Genomics reference GRCh38-2020-A. Automatically called cells were further filtered to ensure usage of high-quality droplets with captured cells. RNA barcodes were filtered on UMI count (>500 UMIs), feature count (>250 features), and percentage of mitochondrial genes (<10%). Cell hashing demultiplexing was performed using Seurat’s HTODemux. Only cells labeled with a single hashing oligo (90.69% of cells) were kept for downstream analysis (doublets marked by 2+ hashing oligos and unlabeled cells were excluded). Most biological samples consisted of several thousand cells (4,833±3,447 cells/sample). To achieve even cell distribution across samples, one outlier sample (19,000 cells) was randomly down-sampled to 10,000 cells and samples with less than 1,000 cells were removed from subsequent analysis, resulting in a total of 21 final samples (uninjured: n=12, VML-injured n=9). Seurat^217^ (v4.3.0) was used for normalization, identification of variable genes, scaling, principal component analysis, UMAP dimensional reduction, and SNN generation followed by Leiden clustering for RNAseq. Clusters were identified using a combination of marker genes and differential expression comparing each cluster to all other cells in the data set. Small irrelevant clusters (e.g., contamination of myeloid and stromal cells) were removed. Differential expression analyses, such as comparing tumor-specific AH1-Tet^+^ CD8^+^ T cells between uninjured and VML-injured groups, were performed using Mann-Whitney U test. Gene set enrichment analysis (GSEA) was run using fgsea^218^ (v1.24.0) with gene sets obtained from the Molecular Signatures Database^219^, including Hallmark and a curated subset of T cell-rated gene sets. Features were ranked by -log(pvalue) * sign(foldchange).

### Bulk RNA-Sequencing

#### Sample Preparation

The transcriptomic phenotypes of CT26 tumor-infiltrating T cells and macrophages were profiled using bulk RNA-sequencing (RNAseq). Mice were randomized between uninjured, untreated VML injury, and ECM-treated VML injury groups with subcutaneous CT26 tumor inoculation 7 days post-injury. CT26 tumors were harvested 14 days post tumor cell inoculation (21 days post VML injury) and processed using the flow cytometry protocol detailed above including a Percoll density gradient. Viability and surface marker staining were performed as detailed above (**Table S8**). Three tumor samples were pooled together to form one “biological replicate” to achieve adequate cell yield, with a final number of three “biological replicates” per experimental group for sorting and bulk RNAseq analysis. Cell sorting was performed by the JHU SKCCC High Parameter Flow Core on a 4-laser BD FACSAria Fusion Cell Sorter. T cells (SSC^lo^, Singlets, Live, CD45^+^, NKp46^-^, CD3^+^) and macrophages (Singlets, Live, CD45^+^, CD11b^+^, Ly6G^-^, Ly6C^lo/mid^, F480^mid/hi^) were sorted directly into DNA LoBind tubes (Eppendorf 022431048) filled with of sterile Buffer RLT Plus (Qiagen 1030963) supplemented with 1x 2-Mercaptoethanol (Gibco 21985023). After sorting, tubes were briefly vortexed and left at room temperature for 1-2 minutes to allow for cell lysis, followed by snap freezing on dry ice and storage at −80°C.

#### RNA Isolation and Quality Control (QC)

DNA/RNA isolation and QC was performed by the Johns Hopkins Single Cell and Transcriptomics Core. DNA/RNA was isolated using the AllPrep DNA/RNA Mini Kit (Qiagen 80204) compatible with ≤5×10^5^ sorted cells. RNA yield was quantified using NanoDrop Spectrophotometer and quality assessed using the Agilent Fragment Analyzer. All RNA samples had acceptable RNA Quality Number (RQN) scores with values between 7.5-10.

#### Library Preparation and Bulk RNAseq

Library preparation and bulk RNAseq was performed by the Johns Hopkins Single Cell and Transcriptomics Core. Complementary DNA synthesis and stranded Total RNA Seq library preparation with unique dual indexes (UDI) was performed using Illumina Standard Total RNA Ligation Kit. QC was assessed with Qubit and Agilent Fragment Analyzer. Libraries were pooled and pair-end sequencing performed using the Illumina NovaSeq platform with a S1_100 flow cell at a targeted depth of 50 million unique reads per sample.

#### Bulk RNAseq Data Analysis

FASTQ files were aligned using STAR^220^ against the GENCODE release M27 (GRCm39) mouse genome assembly and annotations^221^. Normalized counts are scaled by the library size factor of the sample and presented as counts per million. Differential expression analysis was performed with DESeq2^222^ based on a model using the negative binomial distribution with shrinkage estimation of logarithmic fold change. Gene expressions were modeled separately for each sorted cell type (T cells and Macrophages) with experimental condition (uninjured, VML injury, and ECM-treated VML injury) as the only variable included in the design matrix. Reported p-values are Benjamini-Hochberg adjusted to account for multiple testing.

### Retrospective Clinical Cohort Analysis using TriNetX Database

#### Retrospective Cohort Study

We utilized the TriNetX database (https://trinetx.com) (accessed on October 14, 2025, with 158,799,917 patients receiving care across 112 Health Care organizations globally, at the time of analysis) to identify breast cancer patients who received ICB therapy. All patients included in our analysis were required to have a documented ICD-10 diagnosis code of C50 (malignant neoplasm of breast) and receive ICB treatment blocking either PD1 (Pembrolizumab: RxNorm 1547545, Nivolumab: RxNorm 1597876, Dostarlimab: RxNorm 2539967, Cemiplimab: RxNorm 2058826), PDL1 (Atezolizumab: RxNorm 1792776, Avelumab: RxNorm 1875534, Durvalumab: RxNorm 1919503), or CTLA4 (Ipilimumab: RxNorm 1094833, Tremelimumab: RxNorm 2619313). To evaluate whether placement of a biological implant (e.g., acellular dermal matrix scaffold) following mastectomy can impact the overall survival of breast cancer patients treated with ICB therapy, we constructed two cohorts. (1) The experimental cohort included breast cancer patients who both underwent ‘complete simple mastectomy’ (CPT 19303) and received ‘implantation of biologic implant for soft tissue reinforcement’ (CPT 15777) on or up three months after any instance of ICB treatment. (2) The control cohort consisted of breast cancer patients who underwent complete simple mastectomy (CPT 19303) on or up to three months after any instance of ICB treatment and never received a biological implant (CPT 15777).

#### Statistical Analysis

Overall survival (endpoint criteria: deceased) was compared between the two cohorts using a Kaplan-Meier curve and Log-Rank test (* p<0.05). Available categorical data (e.g., demographics, cancer co-treatment) were analyzed using Fisher’s Exact test and age was analyzed using unpaired, two-tailed T test (* p<0.05).

## Data and Code Availability

<u>Single-cell RNAseq</u> raw FASTQ files and aligned counts files for both RNA and hashing features have been made available for download through NCBI GEO under the access number **GSE311280**. Processed objects are available upon reasonable request.

Bulk RNAseq raw FASTQ files and aligned counts files have been made available for download through NCBI GEO under the access number **GSE308871**.

<u>Scripts</u> used for data analysis and visualization in this work were written in R. Scripts for all analysis of scRNAseq and bulk RNAseq are publicly accessible in the GitHub repository hosted at https://github.com/Elisseeff-Lab/vml_injury_and_tumors.

## Acknowledgements

We would like to thank the Johns Hopkins University (JHU) animal facilities and caretakers, the JHU SKCCC High Parameter Flow core facility for performing FACS, the NIH Tetramer Core Facility (NIH Contract 75N93020D00005 and RRID:SCR_026557) for providing the “H2-Ld | MuLV gp70 env 423-431 | SPSYVYHQF | APC” tetramer, Dr. Katlin Stivers for maintaining Cytek Auroa cytometer and providing flow cytometry expertise, the JHU Transcriptomics & Deep Sequencing core facility for performing bulk RNA sequencing, Psomagen and Linda Orzolek for performing single-cell RNA sequencing, ACell Integra LifeSciences for providing ECM scaffold materials, Dr. Sudipto Ganguly for sharing reagents and providing expertise in preclinical cancer models, and Dr. Lisa Jacobs and Dr. Gedge Rosson for their expertise and helpful discussions on clinical treatment of breast cancer patients and biological scaffold use for reconstruction. All schematic figures were created using BioRender.

This research was supported, in part, by the Intramural Research Program of the National Institutes of Health (NIH), National Cancer Institute, CCR, CIL. The contributions of the NIH author(s) were made as part of their official duties as NIH federal employees, are in compliance with agency policy requirements, and are considered Works of the United States Government. However, the findings and conclusions presented in this paper are those of the author(s) and do not necessarily reflect the views of the NIH or the U.S. Department of Health and Human Services

## Funding

This study was funded by the NIH Pioneer Award DP1AR076959 (J.H.E.), NSF Graduate Research Fellowship Program (A.R. DGE1746891), NIH T32 Postdoctoral Research Training Program 3T32DC000027-30S1 (J.A.G.), K99AG081564 (J.C.M), Mark Foundation for Cancer Research (K.N.S., D.M.P.), Bloomberg∼Kimmel Institute for Cancer Immunotherapy (K.N.S., D.M.P).

## Author Contributions

A.R. and J.H.E. conceptualized and designed the studies. A.R. performed experiments and interpreted results. E.F.G.G., J.A.G, L.D.H, J.C.M., D.R.M., and P.A. assisted with experimental work. Y.M. performed clinical cohort analysis. C.C., M.P., and K.K. performed computational analyses. M.T.W. and K.N.S. advised on the studies. A.R. and J.H.E. wrote the manuscript. D.M.P. and J.H.E. supervised the studies.

## Competing Interests

A.R. is an independent contractor for Aegeria Soft Tissue. C.C. is the owner of C M Cherry Consulting. K.N.S holds founders’ equity in Clasp Therapeutics. D.M.P. is a consultant at Aduro Biotech, Amgen, Astra Zeneca, Bayer, Compugen, DNAtrix, Dynavax Technologies Corporation, Ervaxx, FLX Bio, Immunomic, Janssen, Merck, and Rock Springs Capital. D.M.P. holds equity in Aduro Biotech, DNAtrix, Ervaxx, Five Prime therapeutics, Immunomic, Potenza, Trieza Therapeutics. D.M.P. is a member of the scientific advisory board for Bristol Myers Squibb, Camden Nexus II, Five Prime Therapeutics, and WindMil. D.M.P. is a member of the board of directors in Dracen Pharmaceuticals. J.H.E. holds equity in Unity Biotechnology and Aegeria Soft Tissue and is a consultant for Tessara.

## Notes

https://github.com/Elisseeff-Lab/vml_injury_and_tumors

