## Supplementary Figures and Tables for "Tissue Injury and Biomaterial Treatment Modulate Tumor Growth and Response to Immunotherapy"

1 Translational Tissue Engineering Center, Johns Hopkins School of Medicine; Baltimore, MD, USA.
2 Department of Biomedical Engineering, Johns Hopkins School of Medicine; Baltimore, MD, USA.
3 Department of Otolaryngology-Head and Neck Surgery, Vanderbilt University Medical Center; Nashville, TN, USA.
4 Department of Biomaterials & Applied Oral Sciences, School of Biomedical Engineering, Department of Microbiology & Immunology, Dalhousie University; Halifax, NS, Canada.
5 Department of Chemical and Biomolecular Engineering, Johns Hopkins University; Baltimore, MD, USA.
6 Wallace H. Coulter Department of Biomedical Engineering, Georgia Institute of Technology and Emory University; Atlanta, GA, USA.
7 Department of Bioengineering, Swanson School of Engineering, University of Pittsburgh; Pittsburgh, PA, USA.
8 Cancer Biomaterials Engineering Section, Cancer Innovation Laboratory Center for Cancer Research, National Cancer Institute; Frederick, MD, USA.
9 Bloomberg~Kimmel Institute for Cancer Immunotherapy, Sidney Kimmel Comprehensive Cancer Center, Johns Hopkins School of Medicine; Baltimore, MD, USA.
10 The Mark Foundation Center for Advanced Genomics and Imaging, Johns Hopkins University; Baltimore, MD, USA.

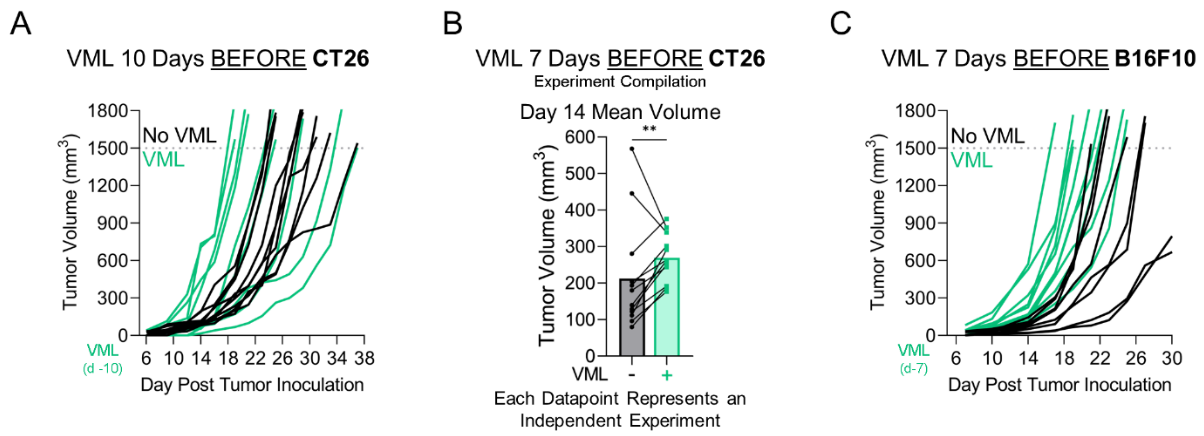

<u>**Suppl Figure 1**</u>**: VML Injury Prior to Tumor Cell Inoculation Accelerates Murine Tumor Growth Kinetics.**
**(A)** Individual CT26 tumor growth curves of uninjured and VML injured mice 10 days prior to tumor cell inoculation. **(B)** Experiment compilation of mean CT26 tumor volumes (day 14) of uninjured and VML injured groups (VML injury performed 7 days prior tumor cell inoculation) (n=12 independent experiments). **(C)** Individual B16F10 tumor growth curves of uninjured and VML injured mice 7 day BEFORE tumor inoculation. **(Statistics)** Experiment Compilation: Each datapoint represents group mean of an independent experiment analyzed using ratio-paired two-tailed T test (*B*). NS: Not significant p>0.05, * p<0.05, ** p<0.01.

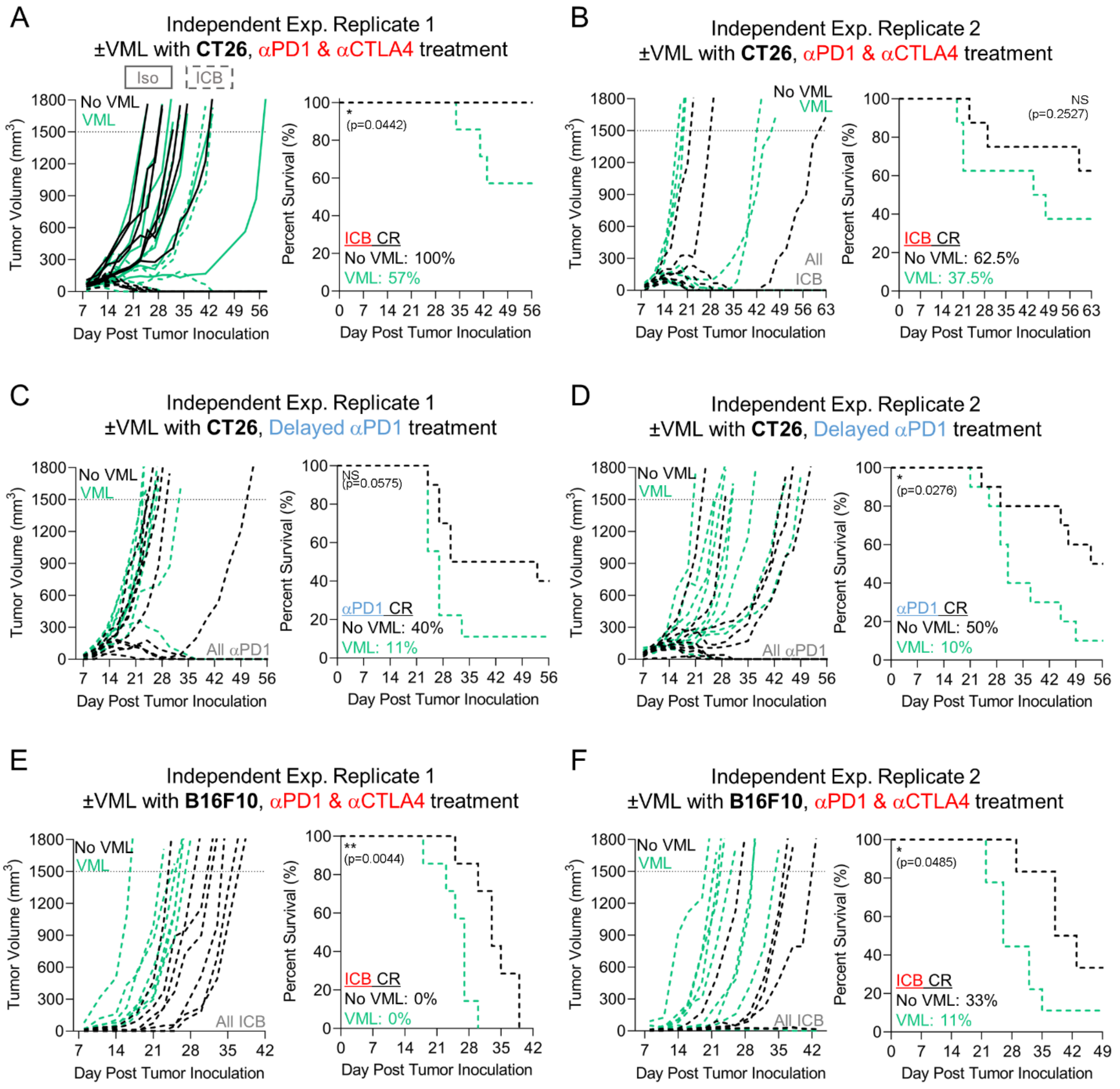

**<u>Suppl Figure 2</u>: VML Injury Prior to Tumor Cell Inoculation Impairs Murine Tumor Response to ICB Therapy.** **(A, B)** Individual CT26 tumor growth and survival curves of uninjured (black) and VML injured (green) mice treated with αPD1/αCTLA4 or **(C, D)** delayed αPD1 monotherapy from independent experiment replicates. **(E, F)** Individual B16F10 tumor growth and survival curves of uninjured (black) and VML injured (green) mice treated with αPD1/αCTLA4 from independent experiment replicates. **(Statistics)** Survival: Kaplan-Meier curve with Log-Rank Mantel-Cox test (*A-F*). Combined survival curves from the matched independent experiments are presented in Figure 1E-1F. NS: Not significant $p>0.05$, * $p<0.05$, ** $p<0.01$. CR: Complete Responder.

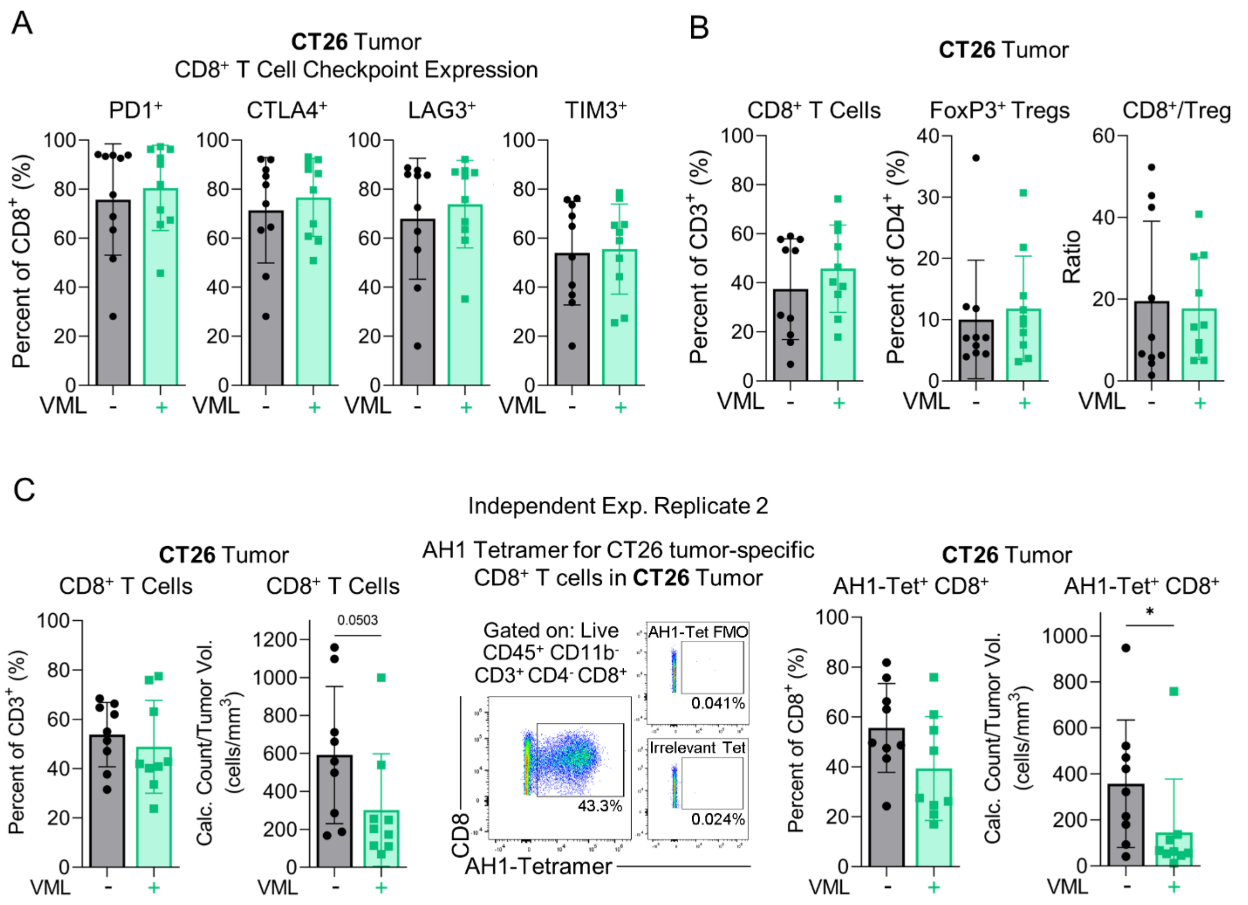

**<u>Suppl Figure 3</u>: VML Injury Decreased Intra-Tumoral Density of CD8$^+$ T Cells But Did Not Alter Their Co-Inhibitory Checkpoint Expression.**

**(A)** Flow cytometric profiling of co-inhibitory checkpoint expression (e.g., PD1, CTLA4, LAG3, TIM3) on CT26 tumor-infiltrating CD8$^+$ T cells of uninjured and VML injured mice. **(B)** Flow cytometric profiling of percentage and relative ratio of CD8$^+$ T cells and FoxP3$^+$ regulatory T cells (Tregs) in CT26 tumors of uninjured and VML injured mice. **(C)** Flow cytometric profiling of percentage and intra-tumoral density (cells/mm$^3$) of all CD8$^+$ T cells and CT26 tumor-specific CD8$^+$ T cells marked by AH1-loaded MHC class I tetramer in uninjured and VML injured mice. **(Statistics)** Bar graphs: mean±SD. Normally distributed data (Shapiro-Wilk test, $\alpha$=0.05) was analyzed using an unpaired two-tailed student t-test, otherwise a non-parametric two-tailed Mann-Whitney test was used (*A*-*C*). NS: Not significant $p>0.05$, * $p<0.05$.

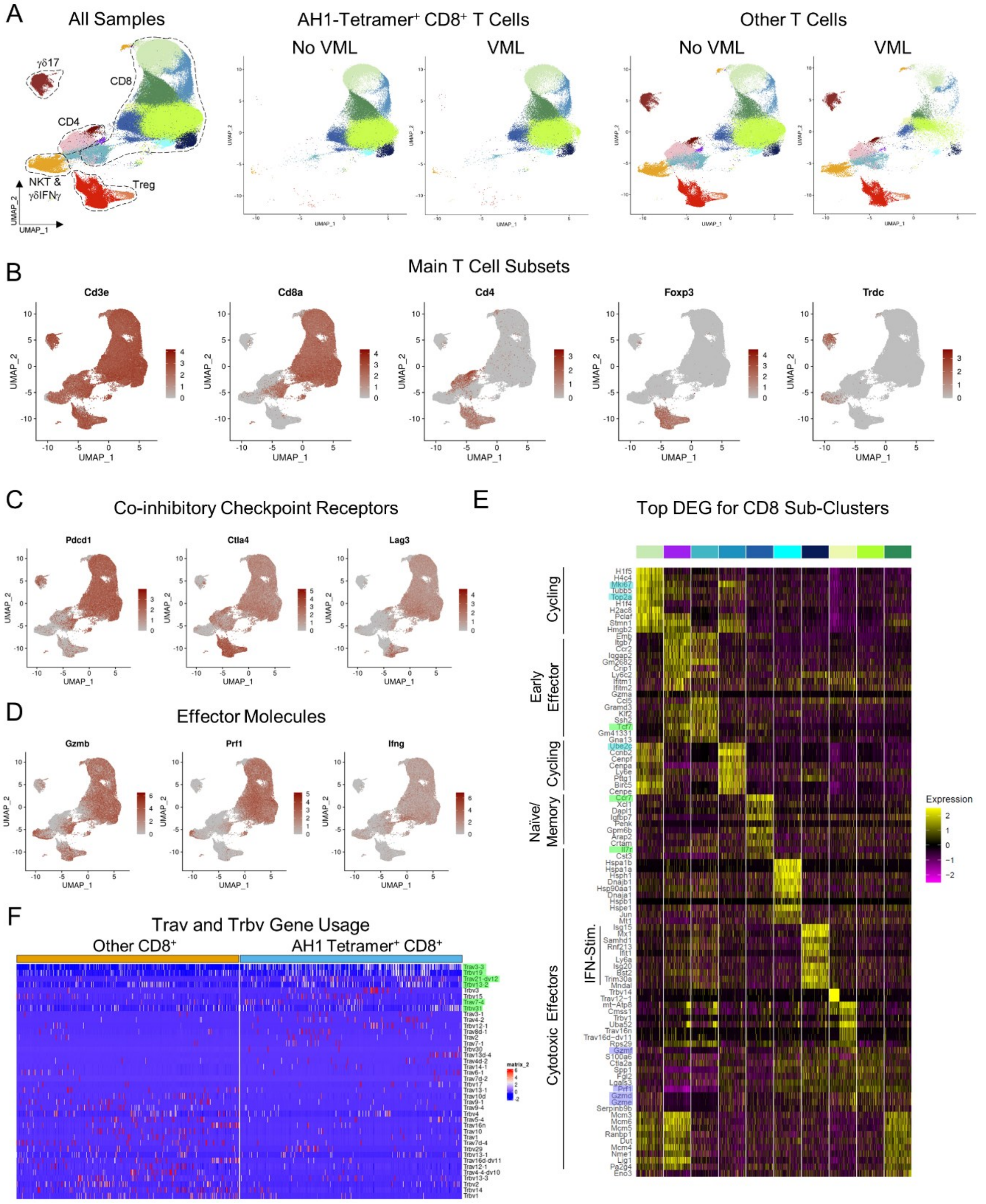

**Suppl Figure 4: Single-Cell RNA-Sequencing Revealed Distinct Subsets of CT26 Tumor-Infiltrating T Cells.** **(A)** Overall UMAP of CT26 tumor-infiltrating T cells separated by sorted cell type (AH1-Tetramer$^+$ CD8$^+$ T cells and Other T cells) and experimental group (No VML and VML). **(B)** Feature plots of select T cell subset marker genes (*Cd8a, Cd4, Foxp3, Trdc*), **(C)** co-inhibitory checkpoint receptors (*Pdcd1, Ctla4, Lag3*), and **(D)** effector molecules (*Gzmb, Prf1, Ifng*). **(E)** Row-scaled heatmap of top 10 differentially expressed genes for the CD8 T cell subclusters with select marker genes names for cycling (*Mki67, Top2a, Ube2c*), naïve/memory (*Tcf7, CCr7, Il7r*), and cytotoxicity (*Gzmd/e/f, Prf1*) highlighted. **(F)** Row-scaled expression heatmap of top 20 *Trav* and *Trbv* genes based on fold-change by group, CT26 tumor-reactive AH1-Tetramer$^+$ CD8$^+$ T cells and all other CD8$^+$ T cells. Previously reported variable region gene usage in AH1-Tetramer$^+$ CD8$^+$ T cells highlighted in green.

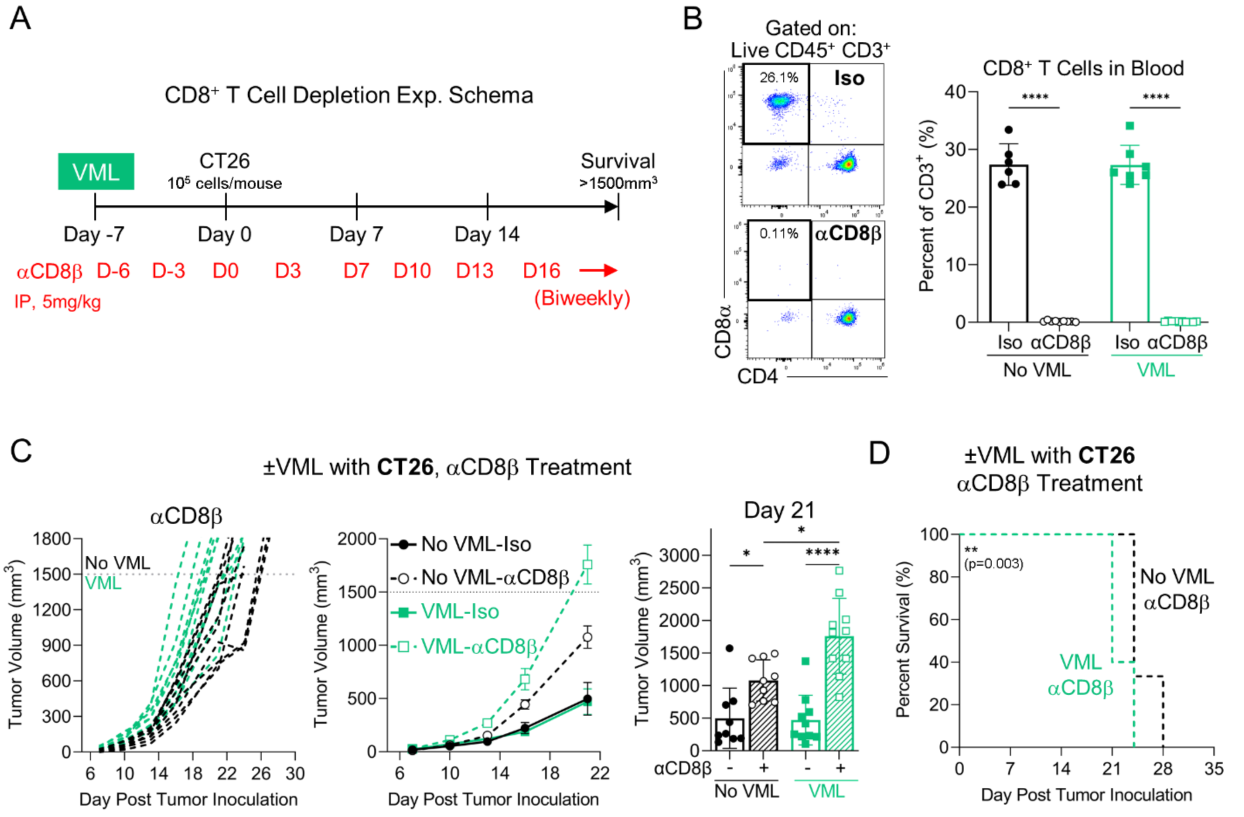

**Suppl Figure 5: CD8 T Cell Depletion Did Not Mitigate VML Injury-Induced Accelerated CT26 Tumor Growth.** **(A)** Experimental scheme of CD8+ T cell depletion with αCD8β antibody treatment. Bilateral VML injury was performed on quadriceps muscles 7 days prior to subcutaneous CT26 tumor inoculation. Isotype or αCD8β (5mg/kg, clone: 53-5.8) treatment began 1-day post-injury and was dosed biweekly until mice met survival criteria (volume >1500mm$^3$ or severe involuting ulceration). **(B)** Representative flow cytometry plots and quantification of CD8+ T cells in peripheral blood (collected at survival endpoint) to confirm successful antibody-mediated CD8+ T cell depletion. **(C)** Individual and mean CT26 tumor growth kinetics and **(D)** survival curves of uninjured (black, circle) and VML injured (green, square) mice treated with isotype (solid) or αCD8β antibody (dashed) (n=9-10). **(Statistics)** Tumor growth curves: mean±SEM. Bar graphs: mean±SD (displaying earliest survival timepoint). Data was analyzed using an ordinary two-way ANOVA with Tukey's multiple comparisons test (only relevant comparisons shown) (*B, C*). Survival: Kaplan-Meier curve with Log-Rank Mantel-Cox test (*D*). NS: Not significant p>0.05, * p<0.05, ** p<0.01, *** p<0.001, **** p<0.0001.

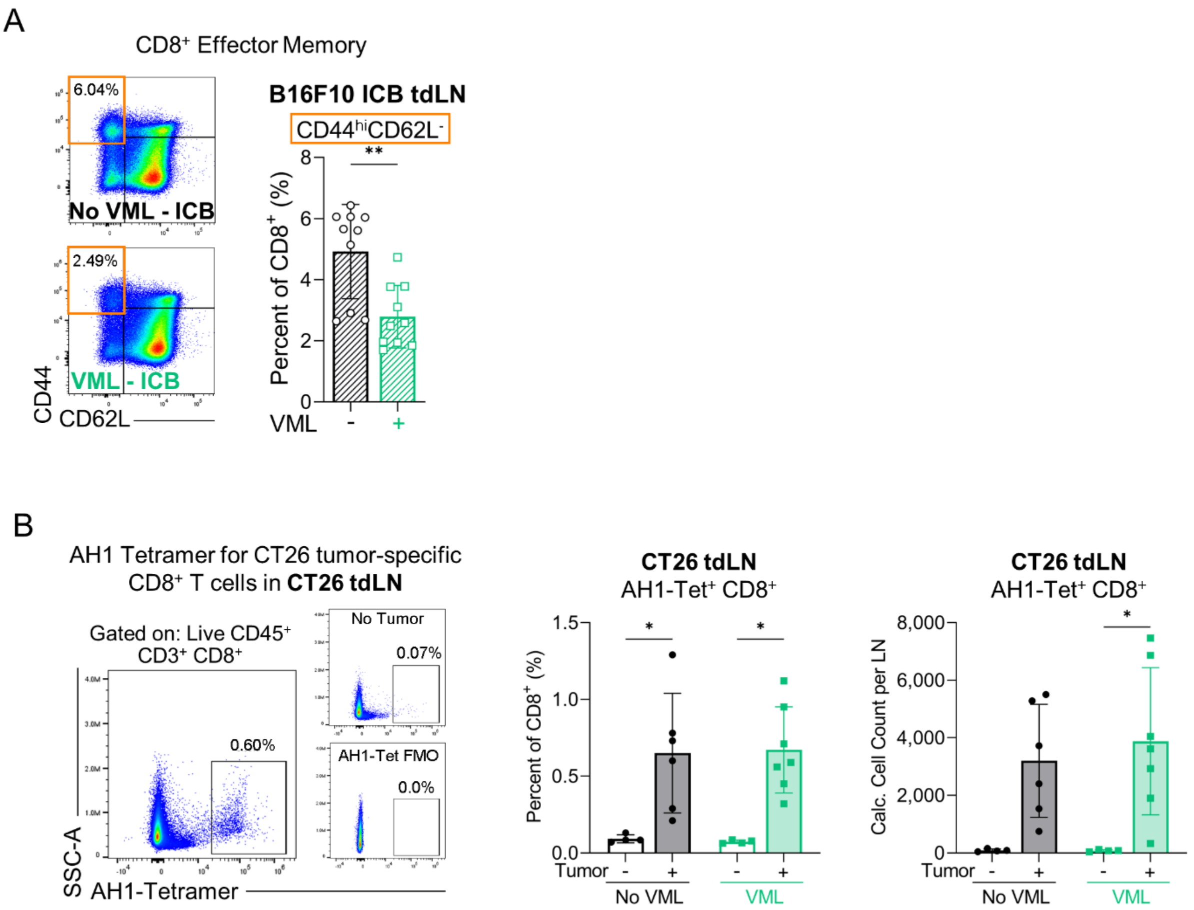

**<u>Suppl Figure 6</u>: CD8⁺ T Cell Profiling in Tumor-Draining Lymph Nodes (tdLNs) in VML-Injured Mice.**
**(A)** Flow cytometric profiling of $CD8^+$ T cell effector memory differentiation ($CD44^{hi}CD62L^-$) in B16F10 tdLNs of uninjured and VML-injured mice treated with αPD1/αCTLA4. **(B)** Flow cytometric profiling of CT26 tumor-specific $CD8^+$ T cells marked by AH1-loaded class I tetramer in inguinal LNs of uninjured and VML-injured mice with and without CT26 tumors. **(Statistics)** Bar graphs: mean±SD. For 2 groups, data was not normally distributed (Shapiro-Wilk test, $\alpha$=0.05) and analyzed a non-parametric two-tailed Mann-Whitney test (*A*). For >2 groups, data was analyzed using an ordinary two-way ANOVA with Tukey's multiple comparisons test (only relevant comparisons shown) (*B*). NS: Not significant p>0.05, * p<0.05, ** p<0.01.

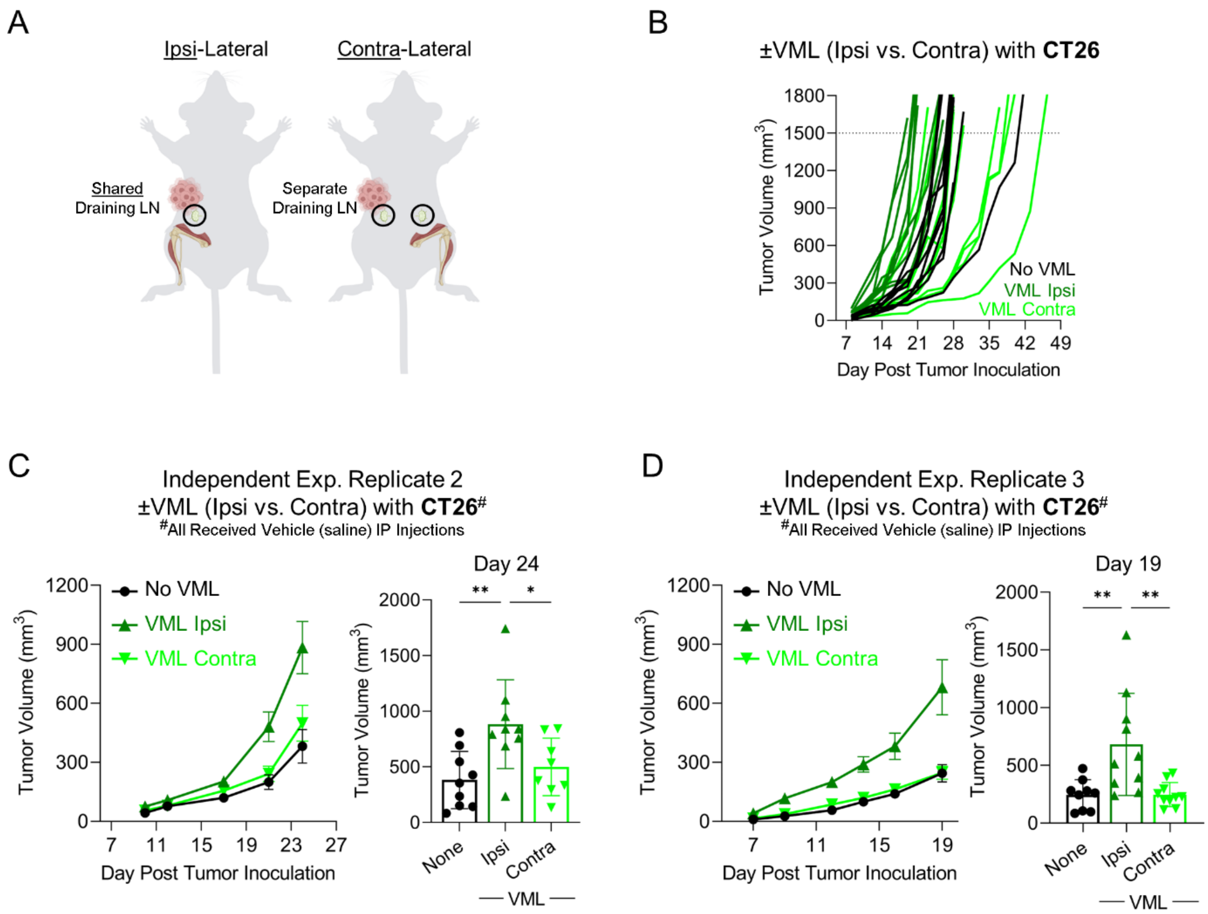

**<u>Suppl Figure 7</u>: Ipsi-lateral VML Injury Accelerates CT26 Tumor Growth Relative to Contra-lateral VML Injury.** **(A)** Schematic of bilateral VML injury modification into a unilateral procedure performed on either the right (ipsi-lateral) or left (contra-lateral) hindlimb quadriceps muscles 7 days prior to subcutaneous CT26 tumor inoculation on the right flank. **(B)** Individual CT26 tumor curves of uninjured (black), ipsi-lateral VML (dark green), or contra-lateral VML (neon green) mice (n=9-10). **(C, D)** Independent experiment replicates showing CT26 tumor growth kinetics of uninjured (black, circle), ipsi-lateral VML (dark green, triangle), and contra-lateral VML (neon green, inverted triangle) mice treated with vehicle (sterile saline) (n=8-10). These studies were performed in tandem with the ones presented in Figure 3G and Suppl Figure 9 (same vehicle group data). **(Statistics)** Tumor growth curves: mean±SEM. Bar graphs: mean±SD (displaying earliest survival timepoint). Data was analyzed using an ordinary one-way ANOVA with Tukey's multiple comparisons test (*C, D*). NS: Not significant $p>0.05$, * $p<0.05$, ** $p<0.01$.

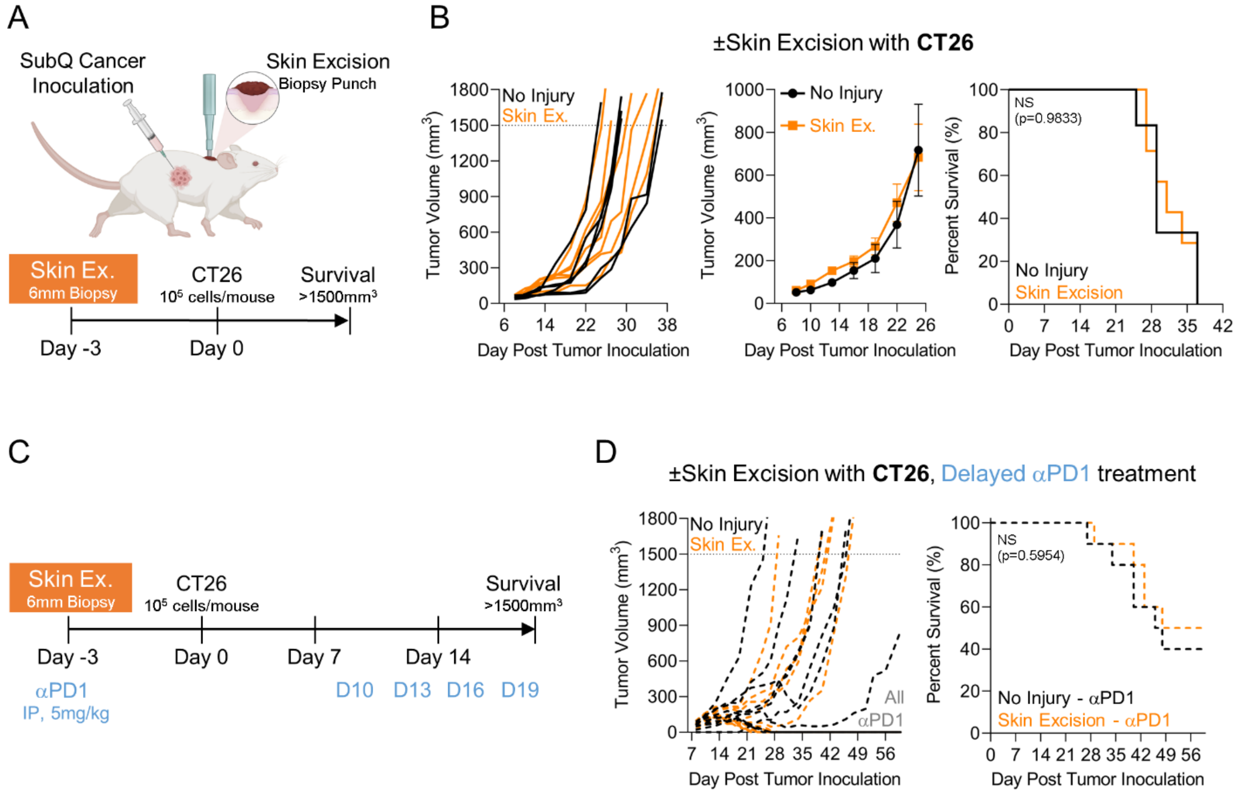

**<u>Suppl Figure 8</u>: Dorsal Skin Excision Injury Does Not Impact CT26 Tumor Growth Kinetics.**
**(A)** Experimental scheme of skin excision (ex.) injury performed via 6mm biopsy punch on mouse dorsum (in line with forelimbs over latissimus dorsi muscle) 3 days prior to subcutaneous CT26 tumor inoculation on the right flank. **(B)** CT26 tumor growth and survival (volume >1500mm$^3$ or severe involuting ulceration) curves of uninjured (black, circle) and skin ex. injured (orange, square) mice (n=6-7). **(C)** Experimental scheme for ICB treatment of concurrent skin ex. injury with CT26 tumor model. αPD1 (5mg/kg) treatment began 10 days post-tumor inoculation and was administered every 3 days for 4 total doses. **(D)** CT26 tumor growth and survival curves of uninjured (black) and skin ex. injured (orange) mice treated with αPD1 (dashed) (n=10). **(Statistics)** Tumor growth curves: mean±SEM. Bar graphs: mean±SD (displaying earliest survival timepoint). Normally distributed data (Shapiro-Wilk test, α=0.05) was analyzed using an unpaired, two-tailed student t-test (*B*). Survival: Kaplan-Meier curve with Log-Rank Mantel-Cox test (*B, D*). NS: Not significant p>0.05, * p<0.05.

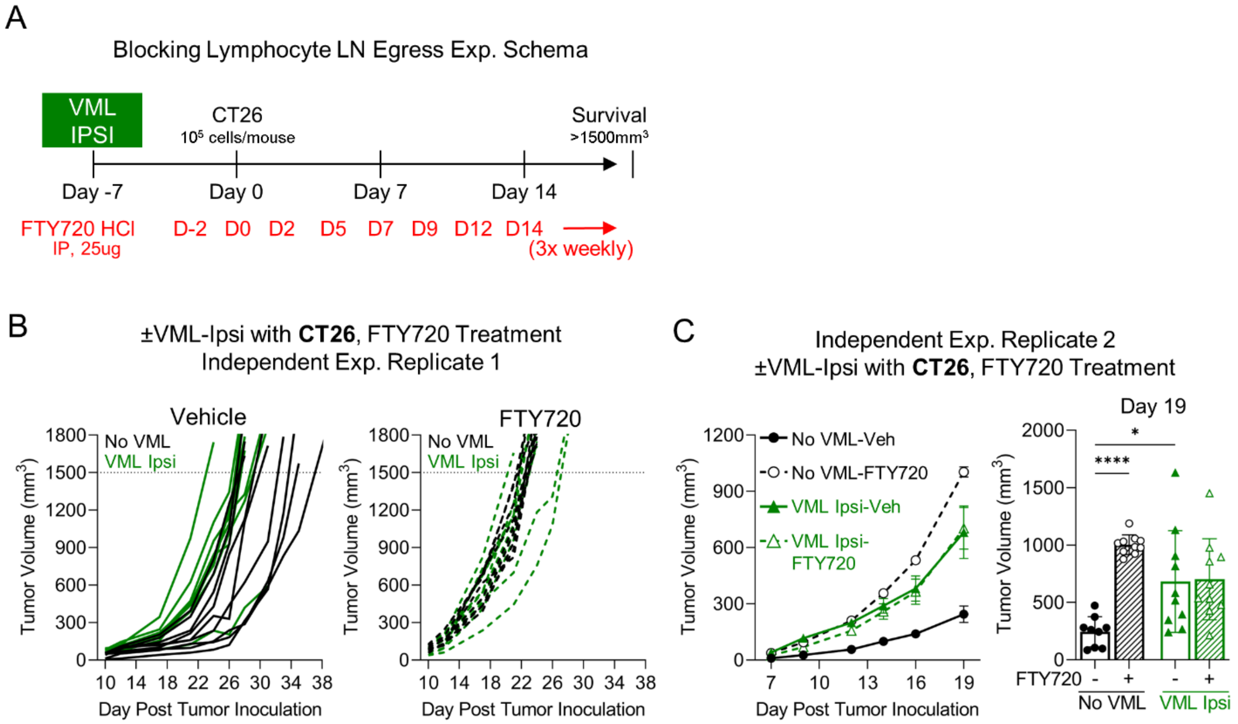

**Suppl Figure 9: T Cell LN Egress Blockade Abrogates VML Injury-Induced Accelerated CT26 Tumor Growth.** **(A)** Experimental scheme of pharmacological inhibition of lymphocyte LN egress using Fingolimod (FTY720) hydrochloride (HCl). Ipsi-lateral VML injury was performed on the right quadriceps muscles 7 days prior to subcutaneous CT26 tumor inoculation on the right flank. Vehicle (sterile saline) or FTY720 HCl (25ug/mouse) treatment began 5 days post-injury delivered via intraperitoneal injection and was dosed 3 times per week until mice met survival criteria (volume >1500mm$^3$ or severe involuting ulceration). **(B)** Individual tumor growth curves of uninjured (black) or ipsi-lateral VML injured (dark green) mice treated with vehicle (solid) or FTY720 HCl (dashed). **(C)** Independent experiment replicate showing CT26 tumor growth kinetics of uninjured or ipsi-lateral VML- injured treated with FTY720. These studies were performed in tandem with the ones presented in Suppl Figure 7C-D (same vehicle group data). **(Statistics)** Tumor growth curves: mean±SEM. Bar graphs: mean±SD (displaying earliest survival timepoint). Data was analyzed using a two-way ANOVA with Tukey's multiple comparisons test (*C*) (only relevant comparisons shown). NS: Not significant p>0.05, * p<0.05, ** p<0.01, *** p<0.001, **** p<0.0001.

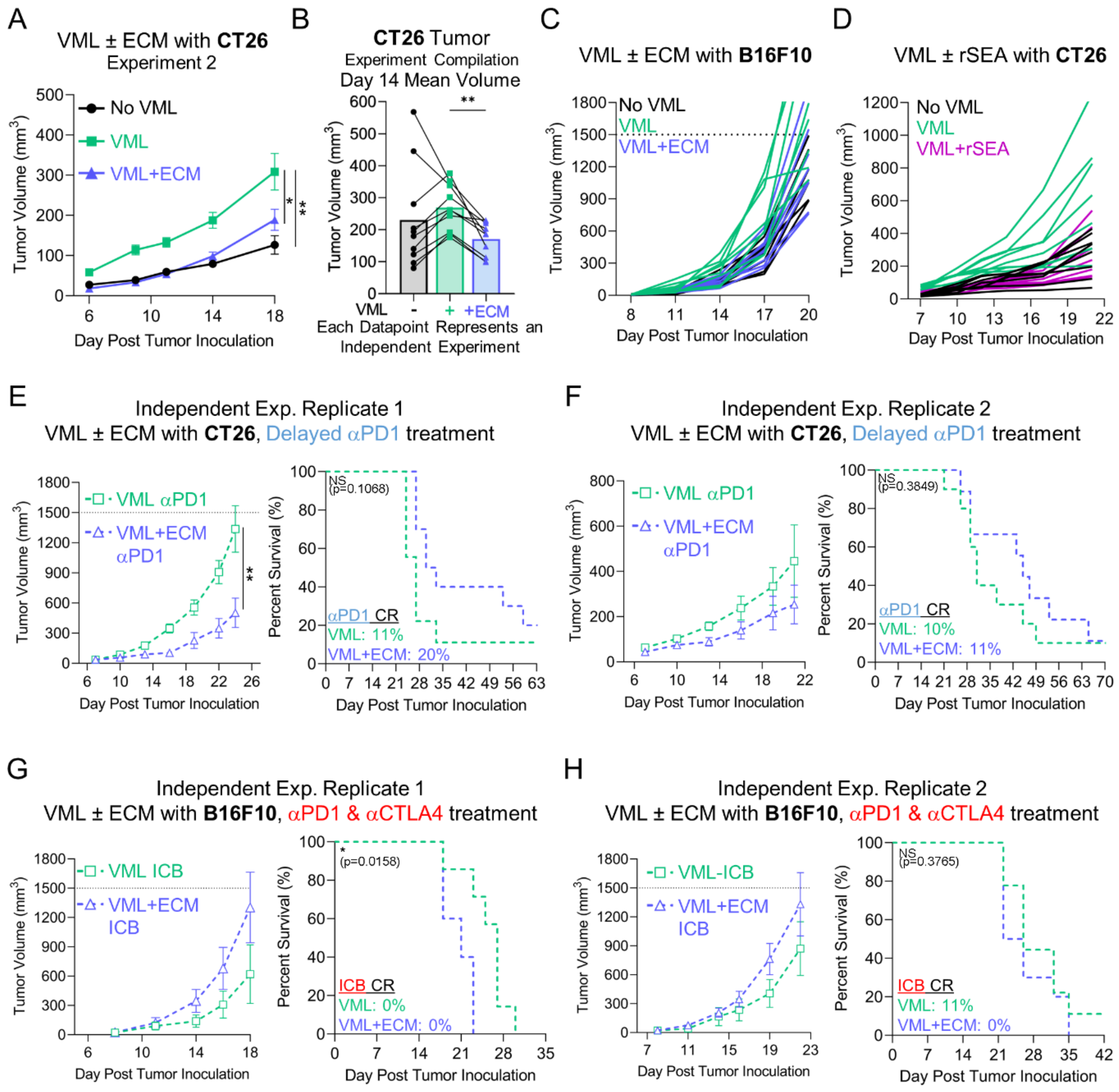

**Suppl Figure 10: Treatment of VML Injury with ECM Scaffolds Slows Injury-Induced Murine Tumor Growth.** **(A)** Independent experiment showing CT26 tumor growth kinetics of uninjured (black, circle), untreated VML-injured (green, square), and extracellular matrix (ECM) scaffold-treated VML-injured (blue, triangle) mice. **(B)** Experiment compilation of mean CT26 tumor volumes on day 14 post-inoculation (n=10 independent experiments). **(C)** Individual B16F10 tumor growth curves of uninjured, untreated VML-injured, and ECM-treated VML-injured mice. **(D)** Individual CT26 tumor growth curves of uninjured, untreated VML-injured, and *Schistosoma mansoni* regenerative soluble egg antigen (rSEA)-treated VML-injured mice. **(E, F)** CT26 tumor growth and survival curves of untreated and ECM-treated VML-injured mice treated with delayed αPD1 monotherapy from independent experiment replicates. These studies were performed in tandem with those presented in Suppl Figure 2C and 2D (same untreated VML injury data). **(G, H)** B16F10 tumor growth and survival curves of untreated and ECM-treated VML-injured mice treated with αPD1/αCTLA4 from independent experiment replicates. These studies were performed in tandem with those presented in Suppl Figure 2E and 2F (same untreated VML injury data). **(Statistics)** Tumor growth curve: mean±SEM. Normally distributed data (Shapiro-Wilk test, $\alpha=0.05$) was analyzed using an unpaired two-tailed student t-test, otherwise a non-parametric two-tailed Mann-Whitney test was used (*E-H, left*). For >2 groups, data was analyzed using ordinary one-way ANOVA with Tukey's multiple comparisons test at final timepoint (*A*). Experiment Compilation: Each datapoint represents an independent experiment with lines connecting datapoints within the same experiment. Data was analyzed using a mixed-effects analysis with Tukey's multiple comparisons test (*B*). Survival: Kaplan-Meier curve with Log-Rank Mantel-Cox test (*E-F, right*). Combined survival curves from the matched independent experiments are presented in Figure 4E and 4F. NS: Not significant p>0.05, * p<0.05, ** p<0.01. CR: Complete Responder.

A

Demographics Table

| | Mastectomy Only | Mastectomy with ECM | p-value |
|---|---|---|---|
| **Total Patients (n)** | 1,018 | 613 | |
| **Sex** | | | >0.9999 |
| Female | 99.7% (1,015) | 100% (613) | |
| **Age** | | | <0.0001 |
| Age at Treatment Initiation (mean ± SD) | 55 ± 13.8 | 46.5 ± 11.7 | |
| **Race** | | | 0.8417 |
| White | 64.2% (654) | 70.6% (433) | |
| Black or African American | 19.3% (196) | 14.8% (91) | |
| Asian | 5.6% (57) | 4.9% (30) | |
| Other | 10.9% (111) | 9.6% (59) | |

**Suppl Fig 11: Demographics of ICB-Treated Breast Cancer Patients With and Without ECM Scaffolds.**
**(A)** Summary table comparing common demographics (e.g., sex, age, race) between breast cancer patients with complete mastectomy, with versus without biological implant placement, on or up to 3 months after ICB treatment. **(Statistics)** Categorical data was analyzed using Fisher's exact test and age using an unpaired, two-tailed t-test (*A*). NS: Not significant p>0.05, * p<0.05.

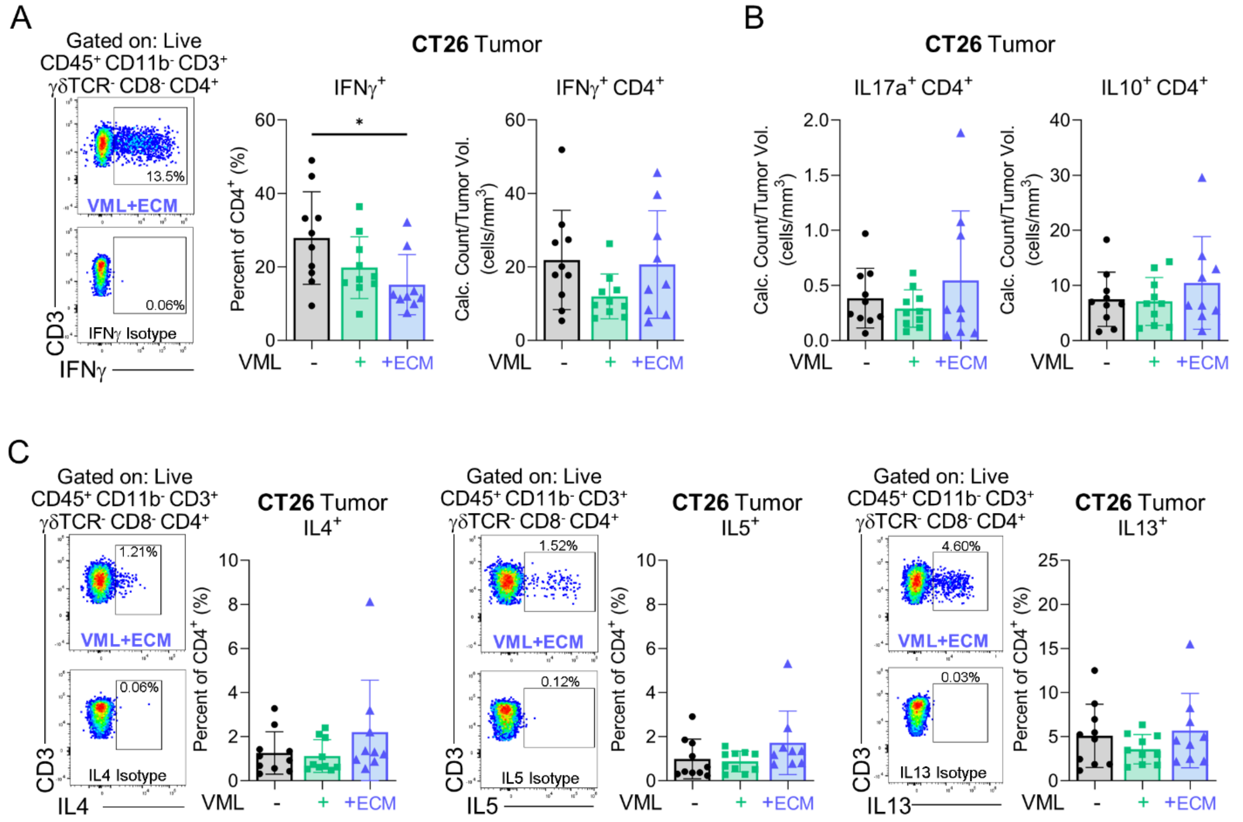

**<u>Suppl Fig 12</u>: CD4$^+$ T Cell Cytokine Production in CT26 Tumors of Untreated and ECM-treated Injured Mice.** **(A, B)** Frequency and/or intra-tumoral density of $T_H1$ (IFNγ+), $T_H17$ (IL17a+) or Tregs (IL10+) from CT26 tumors of uninjured and VML injured (untreated and ECM-treated) mice following *ex vivo* cell stimulation. This study was performed in tandem with the one presented in Figure 2J (same IFNγ+ CD4+ data for uninjured and untreated VML injured groups). **(C)** Representative flow cytometry plots with isotype controls and frequency of type-2 cytokine (IL4, IL5, IL13)-producing $T_H2$ cells from uninjured and VML injured (untreated and ECM-treated) mice following *ex vivo* cell stimulation. **(Statistics)** Bar graphs: mean±SD. Data was analyzed using an ordinary one-way ANOVA with Tukey's multiple comparisons test (*A-C*). NS: Not significant $p>0.05$, * $p<0.05$.

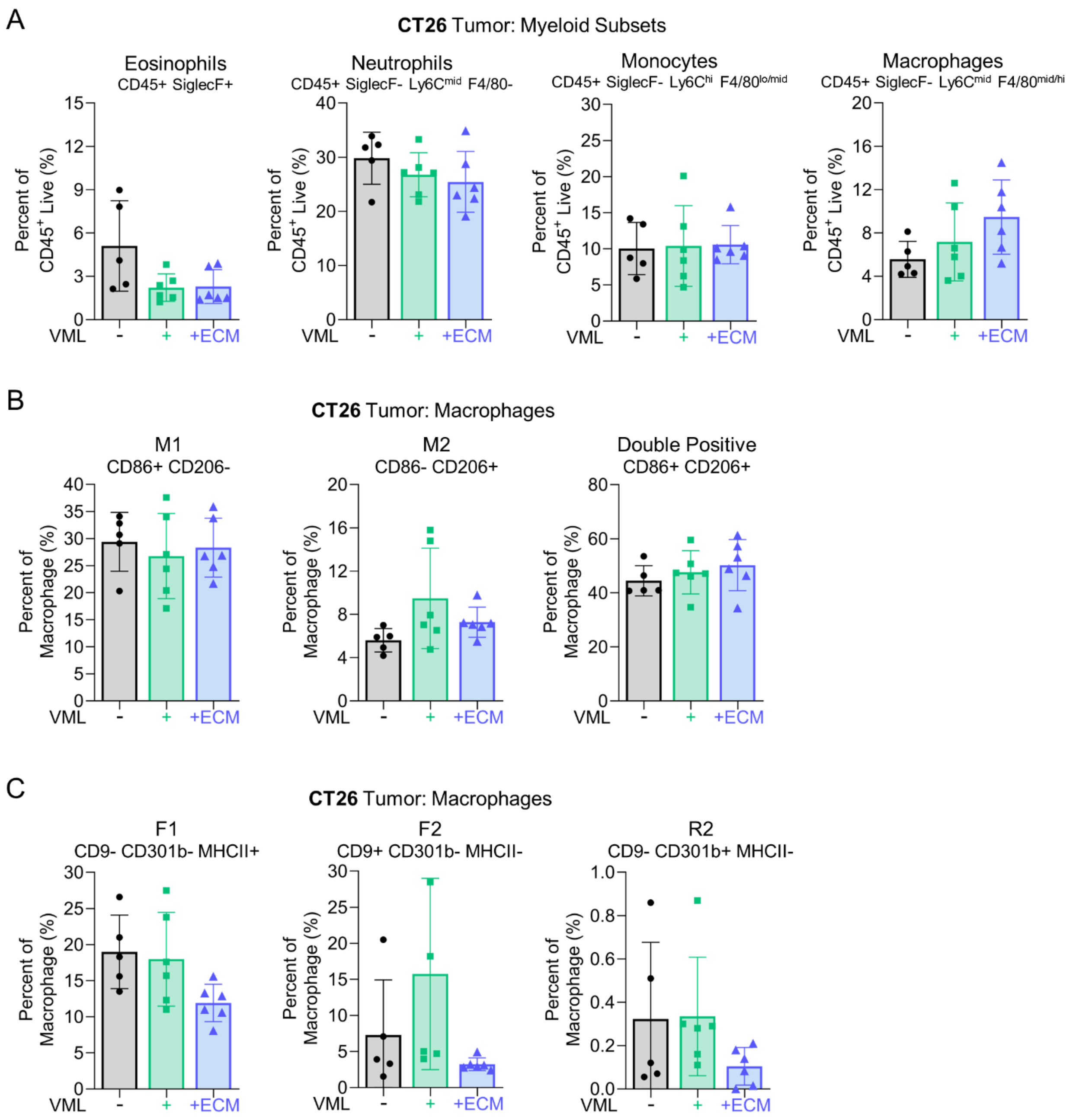

**<u>Suppl Fig 13</u>: ECM Treatment of VML Injury Does Not Alter Conventional Myeloid Subsets in CT26 Tumors.** **(A)** Flow cytometric profiling of broad myeloid subsets (e.g., eosinophils, neutrophils, monocytes, and macrophages) in CT26 tumors of uninjured and VML-injured (untreated and ECM-treated) mice. **(B)** Flow cytometric profiling of macrophage polarization towards conventional M1 (CD86+ CD206-) and M2 (CD86- CD206+) subsets or **(C)** pro-fibrotic and pro-regenerative subsets marked by CD9, CD301b, and MCHII in CT26 tumors of uninjured and VML-injured (untreated and ECM-treated) mice. **(Statistics)** Bar graphs: mean±SD. Data was analyzed using an ordinary one-way ANOVA with Tukey's multiple comparisons test (*A*-*C*). NS: Not significant p>0.05, * p<0.05.

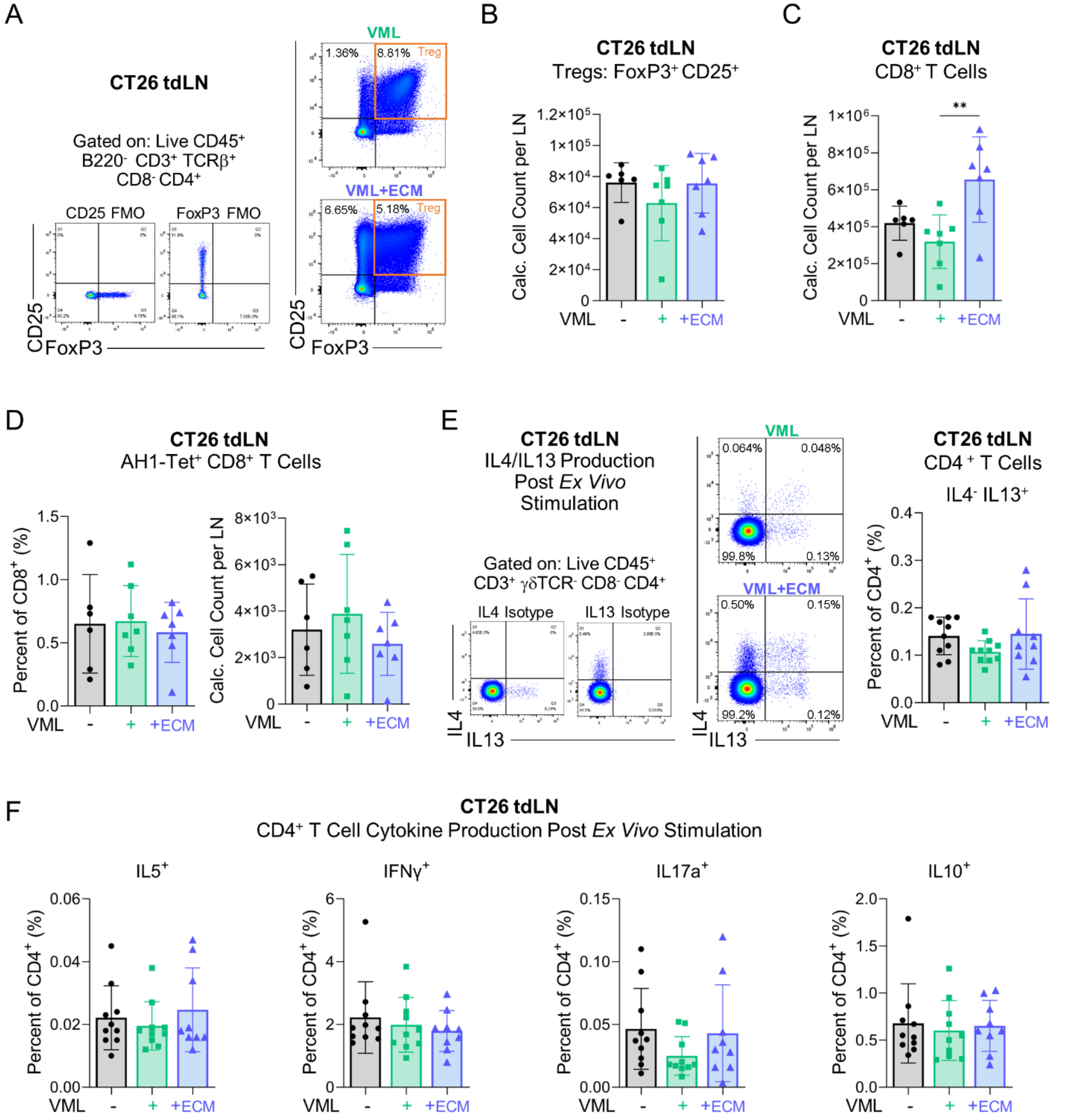

**<u>Suppl Fig 14</u>: T Cell Response in LNs of VML Injured Mice (Untreated and ECM-treated) with CT26 Tumors.** **(A)** Representative flow cytometry gating and **(B)** quantification of Tregs (CD4+ CD25+ FoxP3+) in inguinal LNs of uninjured and VML-injured (untreated and ECM-treated) mice with CT26 tumors. **(C)** Quantification of $CD8^+$ T cells and **(D)** tumor-specific $CD8^+$ T cells marked by AH1-loaded class I tetramer in inguinal LNs of uninjured and VML-injured (untreated and ECM-treated) mice with CT26 tumors. **(E)** Representative gating and quantification of IL4 and IL13 production by $T_H2$ cells or **(F)** other $CD4^+$ T cell cytokines (e.g., IFNγ, IL17a, IL10) following *ex vivo* cell stimulation in inguinal LNs of uninjured and VML-injured (untreated and ECM-treated) mice with CT26 tumors. **(Statistics)** Bar graphs: mean±SD. Data was analyzed using an ordinary one-way ANOVA with Tukey's multiple comparisons test (*A-C*). NS: Not significant p>0.05, * p<0.05, ** p<0.01.

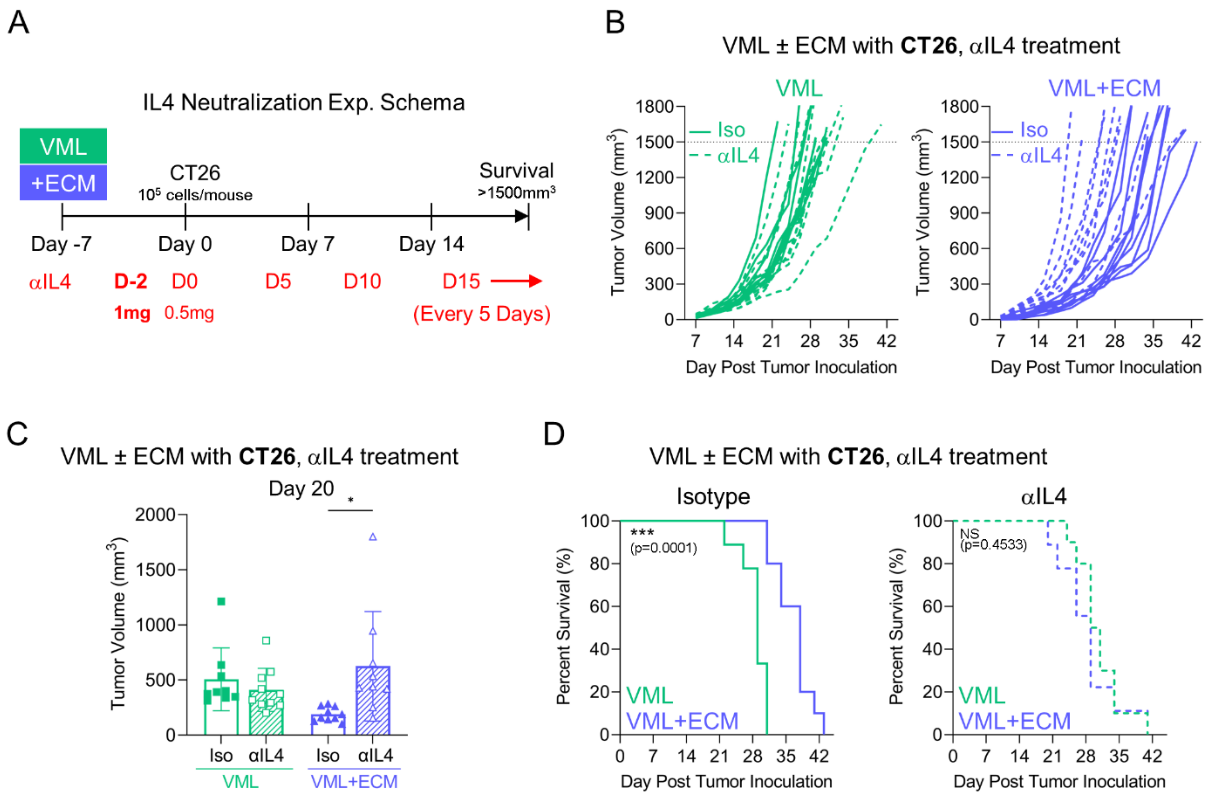

**Suppl Fig 15: IL4 Neutralization Abrogated CT26 Tumor Growth Control in ECM-Treated VML-Injured Mice.**
**(A)** Experimental scheme for disrupting type-2 signaling with an IL4 neutralizing antibody. Bilateral VML injury was performed on quadriceps muscles, either untreated or ECM-treated, 7 days prior to subcutaneous CT26 tumor inoculation on the right flank. Isotype or αIL4 treatment began with an initial high dose (1 mg/mouse) administered 5 days post-injury and continued with a maintenance dose (0.5 mg/mouse) administered 7 days post-injury and dosed every 5 days until survival criteria was met (volume >1500mm$^3$ or severe involuting ulceration). **(B)** Individual CT26 tumor growth curves, **(C)** tumor volume measurements on day 20 (earliest survival timepoint), and **(D)** overall survival curves for untreated (green) and ECM-treated (blue) VML-injured mice receiving isotype (solid) or αIL4 (dashed). **(Statistics)** Bar graphs: mean±SD. Data was analyzed using an ordinary two-way ANOVA with Tukey's multiple comparisons test (*C*). Survival: Kaplan-Meier curve with Log-Rank Mantel-Cox test (*D*). NS: Not significant p>0.05, * p<0.05, ** p<0.01, *** p<0.001.

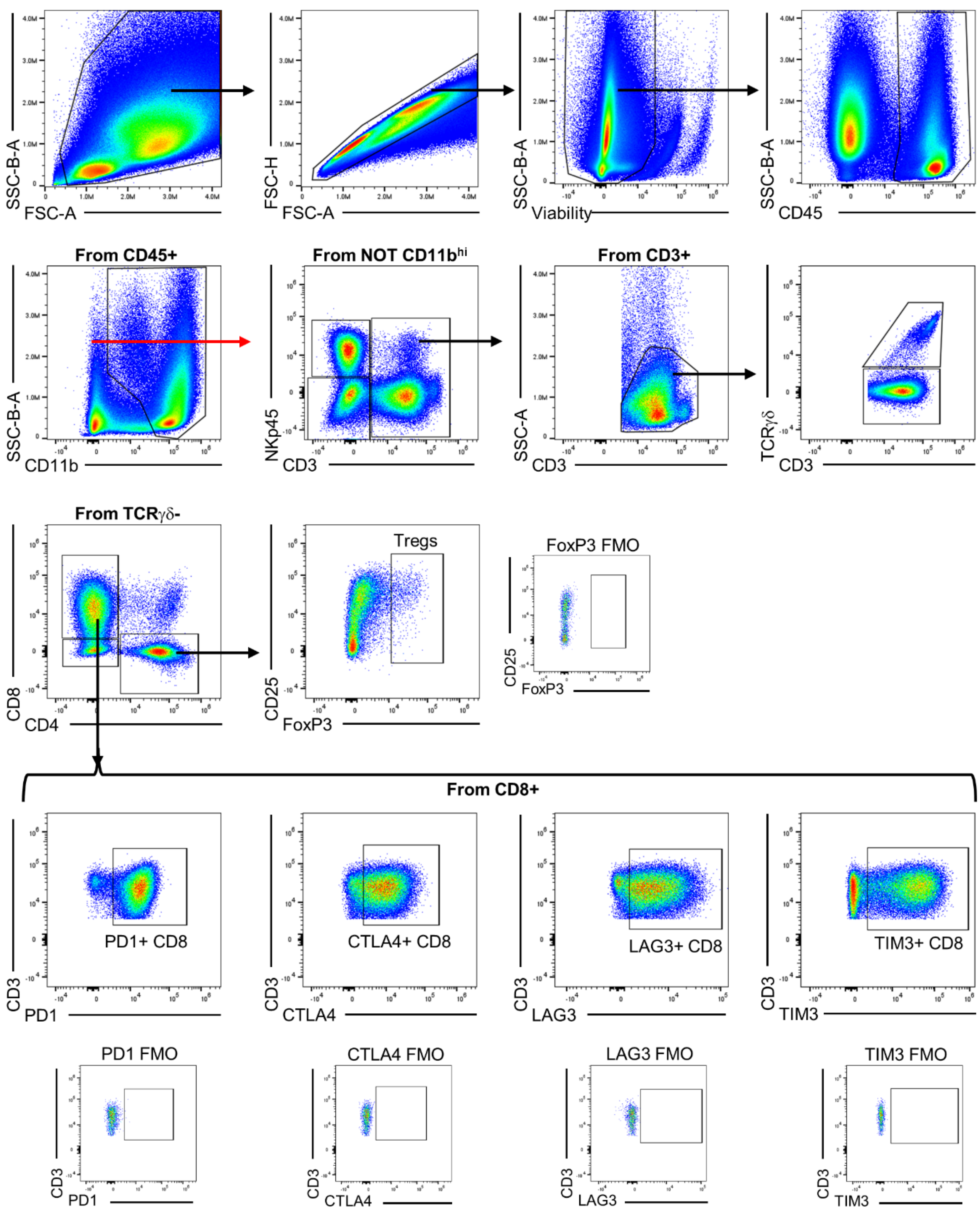

**<u>Suppl Fig 16</u>: Flow cytometry gating scheme for T Cell Checkpoint panel.**

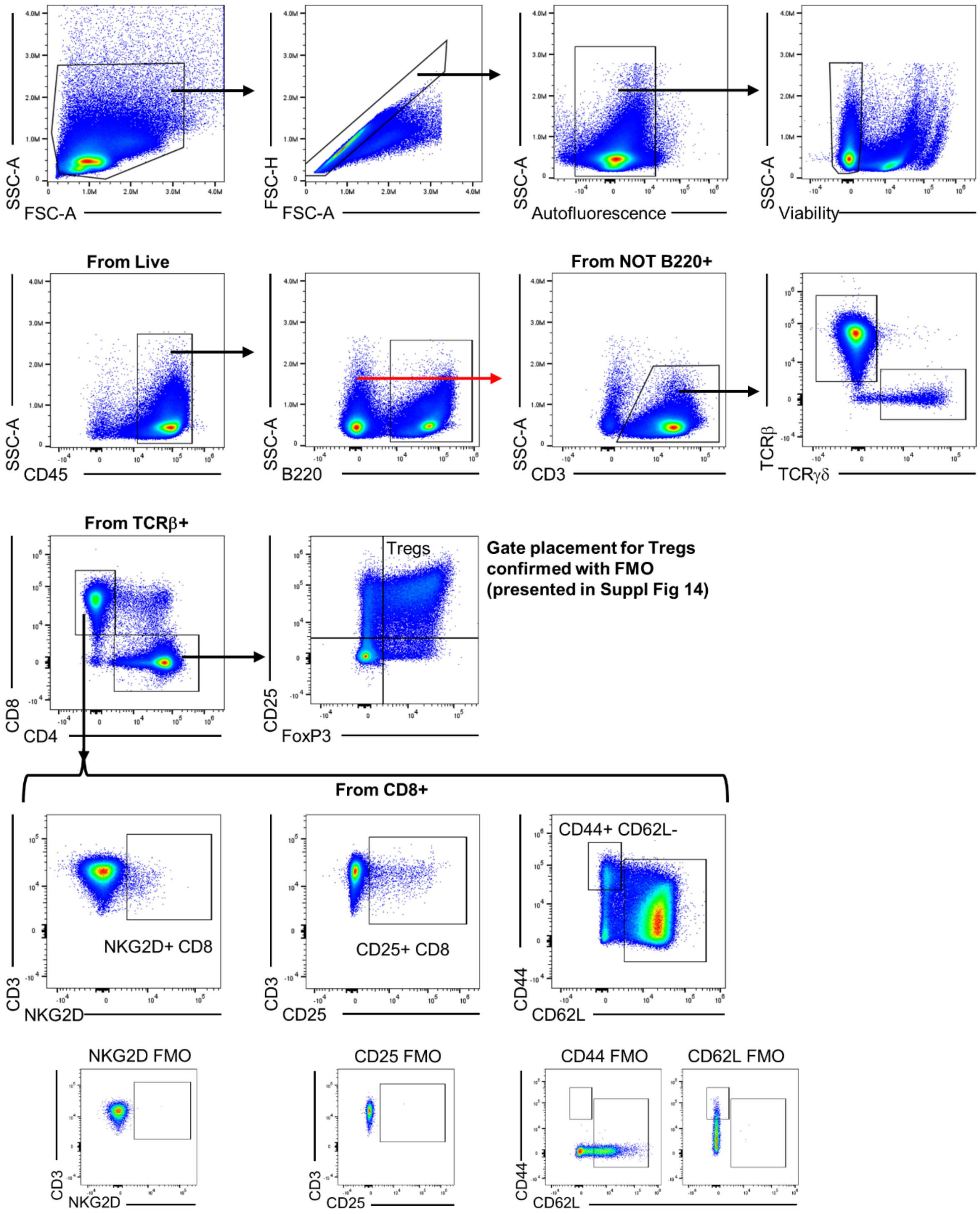

<u>Suppl Fig 17</u>: **Flow cytometry gating scheme for T Cell Effector Memory panel.**

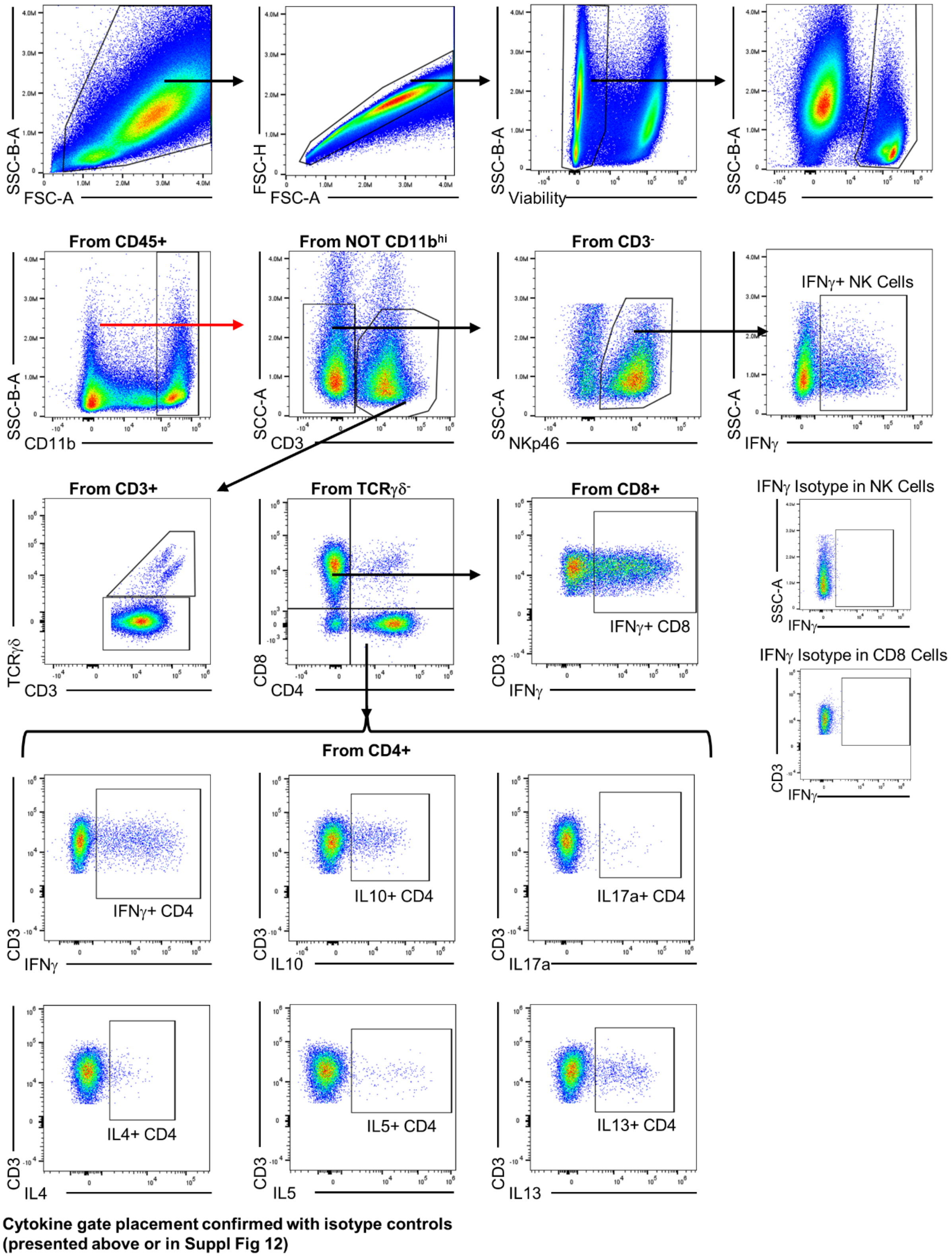

**<u>Suppl Fig 18</u>: Flow cytometry gating scheme for T Cell Intracellular Cytokine panel.**

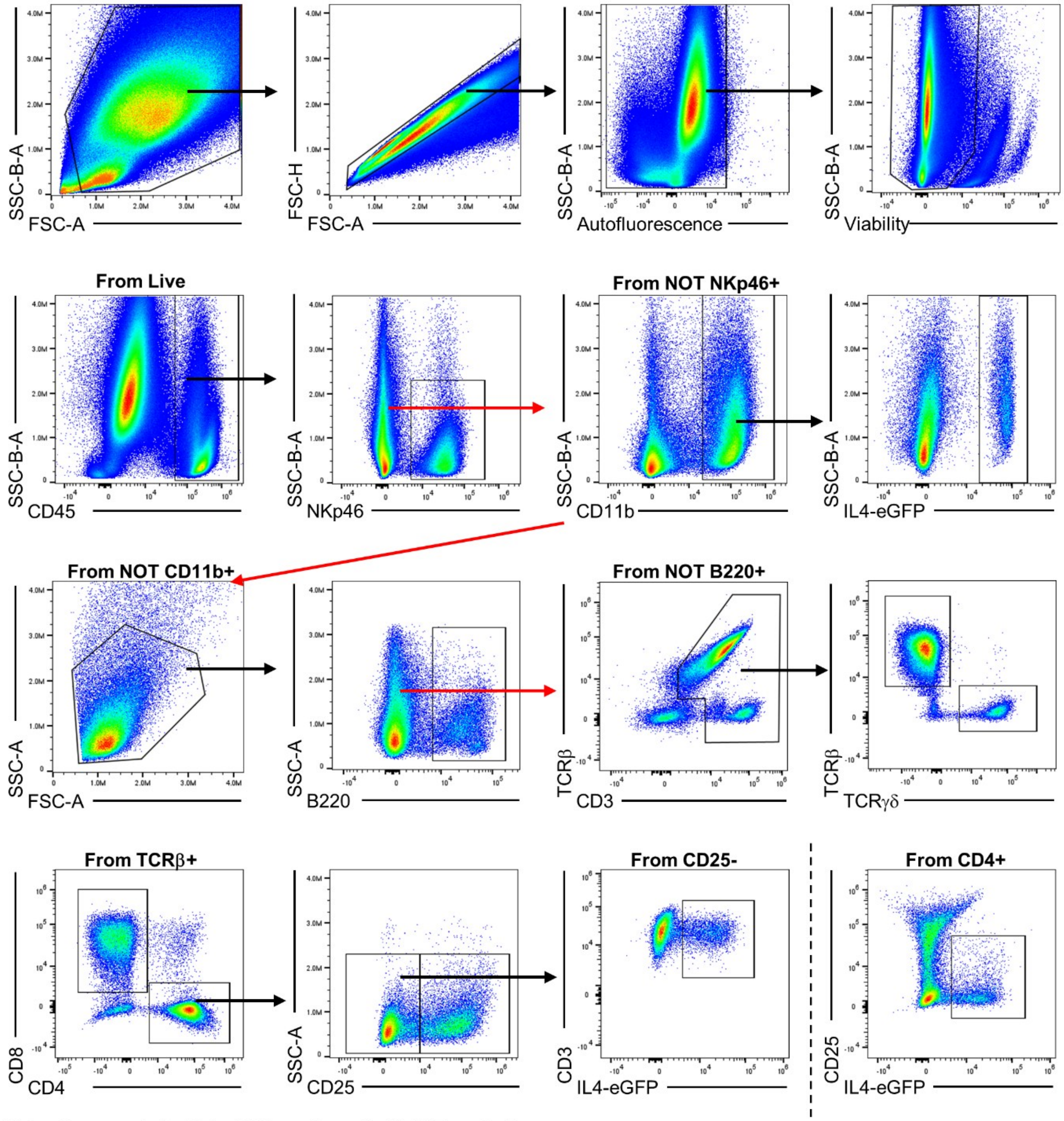

**Gate placements for IL4-eGFP confirmed with WT control (presented in Fig 5 and 6)**

**<u>Suppl Fig 19</u>: Flow cytometry gating scheme for 4get Mouse Strain panel.**

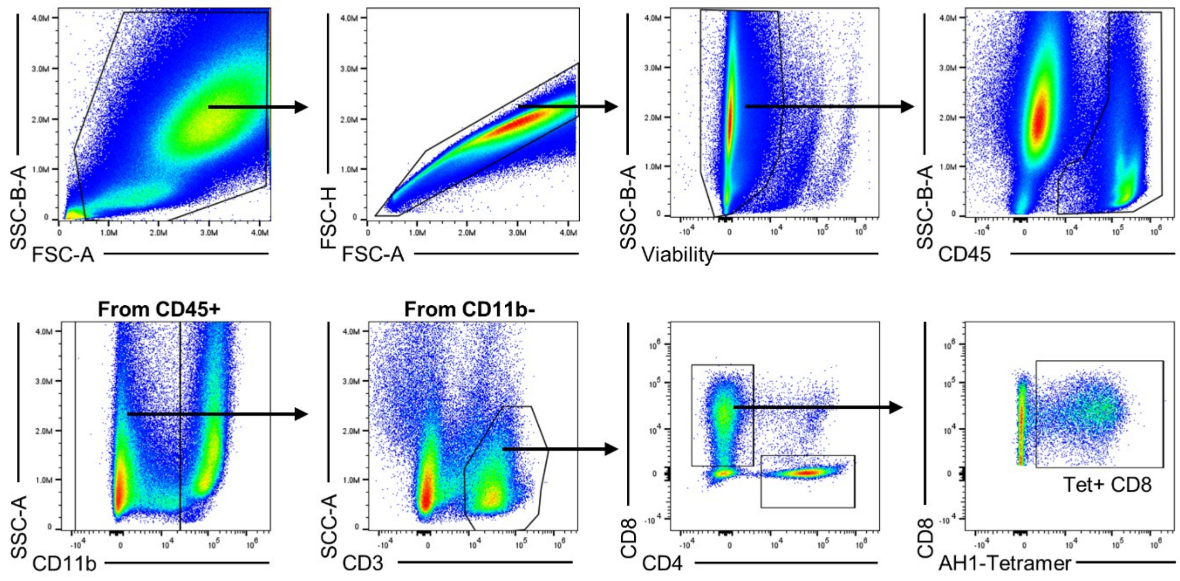

**Gate placement for AH1-Tetramer+ CD8 confirmed with Irrelevant-Tetramer and FMO (presented in Suppl Fig 3)**

**<u>Suppl Fig 20</u>: Flow cytometry gating scheme for T Cell AH1-Tetramer panel.**

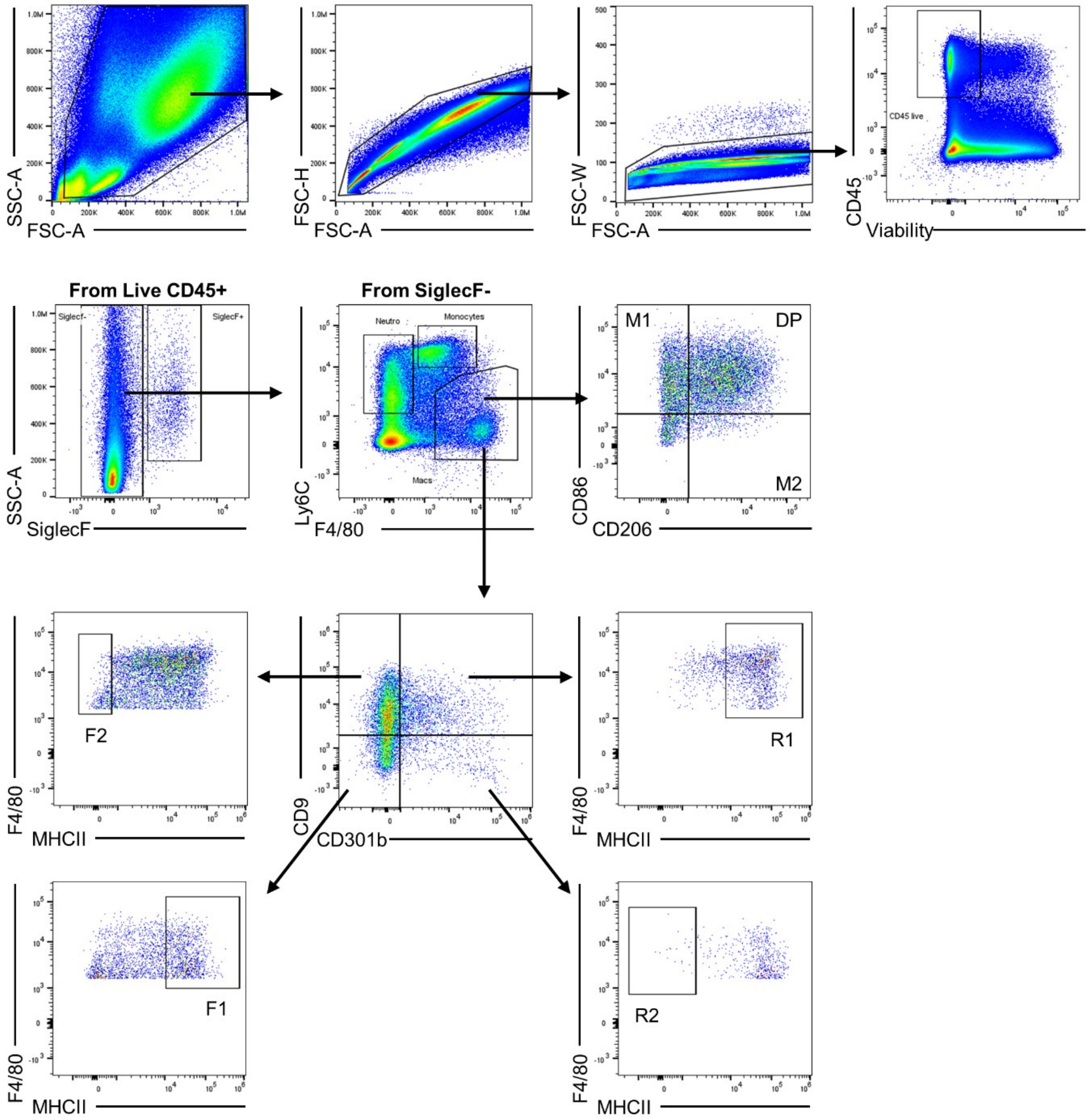

**<u>Suppl Fig 21</u>: Flow cytometry gating scheme for Macrophage Phenotype panel.**

### Supplementary Tables

**Table 1**: T Cell Checkpoint Flow Cytometry Panel

| Fluorophore | Antigen | Clone | Dilution | Manufacturer | Catalog # |
|---|---|---|---|---|---|
| BV421* | NKp46 | 29A1.4 | 50 | BioLegend | 137612 |
| BV421* | NK1.1 | PK136 | 50 | BioLegend | 108732 |
| BV480 | GITR | DTA-1 | 200 | BD Biosciences | 746745 |
| BV510 | CD45 | 30-F11 | 200 | BioLegend | 103138 |
| BV605 | CD4 | GK1.5 | 300 | BioLegend | 100451 |
| BV711 | TCR γ/δ | GL3 | 200 | BD Biosciences | 563994 |
| BV750 | B220 | RA3-6B2 | 200 | BioLegend | 103261 |
| BV786 | PD1 | RPM1-30 | 200 | BD Biosciences | 748264 |
| AF488 | TCF1/7 | C63D9 | 100 | Cell Signaling Technology | 6444S |
| Spark Blue 550 | CD3 | 17A2 | 100 | BioLegend | 100260 |
| BB700 | CD8α | 53-6.7 | 200 | BD Biosciences | 566409 |
| PerCP-eFluor710 | TIGIT | GIGD7 | 100 | Thermo Fisher | 46-9501-82 |
| PE | CD25 | PC61 | 200 | BioLegend | 102008 |
| PE/Dazzle 594 | CTLA4 | UC10-4B9 | 200 | BioLegend | 106318 |
| PE-Cy7 | TIM3 | RMT3-23 | 150 | BioLegend | 119716 |
| APC | Tox | REA473 | 150 | Miltenyi Biotec | 130-118-335 |
| eFluor 660 | FoxP3 | FJK-16s | 100 | Thermo Fisher | 50-5773-82 |
| AF700 | CD11b | M1/70 | 400 | BioLegend | 101222 |
| Zombie NIR | Viability | - | 5000 | BioLegend | 423106 |
| APC-eFluor780 | LAG3 | C9B7W | 100 | Thermo Fisher | 47-2231-82 |

* Depending on mouse strain. BALB/c used NKp46, C57BL/6 used NK1.1
Red text indicates intracellular antibody staining

**Table 2**: T Cell Effector Memory Flow Cytometry Panel

| Fluorophore | Antigen | Clone | Dilution | Manufacturer | Catalog # |
|---|---|---|---|---|---|
| BV421 | TCRβ | H57-197 | 100 | BioLegend | 109230 |
| V450^ | CD27 | LG.3A10 | 100 | BD Biosciences | 561245 |
| BV480 | KLRG1 | 2F1 | 200 | BD Biosciences | 746353 |
| BV510 | CD45 | 30-F11 | 200 | BioLegend | 103138 |
| BV605 | TCR γ/δ | GL3 | 100 | BioLegend | 118129 |
| BV650 | CD44 | IM7 | 300 | BioLegend | 103049 |
| BV711 | NKG2D | CX5 | 100 | BD Biosciences | 563694 |
| BV750 | B220 | RA3-6B2 | 200 | BioLegend | 103261 |
| BV786 | CXCR3 | CXCR3-173 | 100 | BD Biosciences | 741032 |
| KIRAVIA Blue 520 | CD69 | H1.2F3 | 100 | BioLegend | 104554 |
| Spark Blue 550 | CD3 | 17A2 | 100 | BioLegend | 100260 |
| PerCP | Ly6C | HK1.4 | 300 | BioLegend | 128028 |
| BB700 | CD8α | 53-6.7 | 200 | BD Biosciences | 566409 |
| PE | CD25 | PC61 | 200 | BioLegend | 102008 |
| PE/Dazzle 594 | CD62L | MEL-14 | 250 | BioLegend | 104448 |
| PE-Cy5.5 | FoxP3 | FJK-16s | 200 | Thermo Fisher | 35-5773-82 |
| AF647* | NKp46 | 29A1.4 | 200 | BioLegend | 137628 |
| AF647* | NK1.1 | PK136 | 200 | BioLegend | 108720 |
| AF700 | CD11b | M1/70 | 400 | BioLegend | 101222 |
| Zombie NIR | Viability | - | 5000 | BioLegend | 423106 |
| APC-eFluor780 | CD127 | A7R34 | 100 | Thermo Fisher | 47-1271-82 |
| APC-Fire810 | CD4 | GK1.5 | 150 | BioLegend | 100480 |

^ Product has since been discontinued by manufacturer
* Depending on mouse strain. BALB/c used NKp46, C57BL/6 used NK1.1
Red text indicates intracellular antibody staining

**Table 3**: T Cell Intra-Cellular Cytokine Flow Cytometry Panel

| Fluorophore | Antigen | Clone | Dilution | Manufacturer | Catalog # |
|---|---|---|---|---|---|
| BV421 | NKp46 | 29A1.4 | 50 | BioLegend | 137612 |
| BV510 | CD45 | 30-F11 | 200 | BioLegend | 103138 |
| BV605 | CD4 | GK1.5 | 300 | BioLegend | 100451 |
| BV650 | CD11b | M1/70 | 600 | BioLegend | 101259 |
| BV711 | TCR γ/δ | GL3 | 200 | BD Biosciences | 563994 |
| BV786 | IL4 | 11B11 | 100 | BD Biosciences | 564006 |
| Vio B515 | IL10 | REA1008 | 100 | Miltenyi Biotec | 130-116-848 |
| Spark Blue 550 | CD3 | 17A2 | 100 | BioLegend | 100260 |
| BB700 | CD8α | 53-6.7 | 200 | BD Biosciences | 566409 |
| PE | IL5 | TRFK5 | 150 | BioLegend | 504303 |
| PE-eFluor610 | IL13 | eBio13A | 150 | Thermo Fisher | 61-7133-82 |
| APC | IFNγ | XMG1.2 | 150 | BioLegend | 505810 |
| AF700 | IL17A | TC11-18H10.1 | 200 | BioLegend | 506914 |
| Zombie NIR | Viability | - | 5000 | BioLegend | 423106 |
| BV786 | Rat IgG1 | R3-34 | 100 | BD Biosciences | 563847 |
| Vio B515 | Human 1gG1 | REA293 | 100 | Miltenyi Biotec | 130-114-556 |
| PE | Rat IgG1 | RTK2071 | 150 | BioLegend | 400407 |
| PE-eFluor610 | Rat IgG1 | eBRG1 | 150 | Thermo Fisher | 61-4301-82 |
| APC | Rat IgG1 | RTK2071 | 150 | BioLegend | 400411 |
| AF700 | Rat IgG1 | RTK2071 | 200 | BioLegend | 400420 |

Red text indicates intracellular antibody staining

**Table 4**: 4get Mouse Strain Flow Cytometry Panel

| Fluorophore | Antigen | Clone | Dilution | Manufacturer | Catalog # |
|---|---|---|---|---|---|
| BV421* | TCRβ | H57-197 | 100 | BioLegend | 109230 |
| BV421* | Siglec F | E502440 | 150 | BD Biosciences | 562681 |
| BV510 | CD45 | 30-F11 | 200 | BioLegend | 103138 |
| BV605 | CD4 | GK1.5 | 300 | BioLegend | 100451 |
| BV711 | TCR γ/δ | GL3 | 200 | BD Biosciences | 563994 |
| BV750 | B220 | RA3-6B2 | 200 | BioLegend | 103261 |
| BV786 | PD1 | RPM1-30 | 200 | BD Biosciences | 748264 |
| GFP | IL4 | - | - | - | - |
| Spark Blue 550 | CD3 | 17A2 | 100 | BioLegend | 100260 |
| BB700 | CD8α | 53-6.7 | 200 | BD Biosciences | 566409 |
| PerCP-eFluor710 | TIGIT | GIGD7 | 100 | Thermo Fisher | 46-9501-82 |
| PE | CD25 | PC61 | 200 | BioLegend | 102008 |
| PE-Cy7 | TIM3 | RMT3-23 | 150 | BioLegend | 119716 |
| AF647 | NKp46 | 29A1.4 | 200 | BioLegend | 137628 |
| AF700 | CD11b | M1/70 | 400 | BioLegend | 101222 |
| Zombie NIR | Viability | - | 5000 | BioLegend | 423106 |
| APC-eFluor780 | LAG3 | C9B7W | 100 | Thermo Fisher | 47-2231-82 |

* TCRβ for tumors, quads and iLN. SiglecF for adipose tissue.

Red text indicates transgenic reporter

**Table 5**: T Cell AH1-Tetramer Flow Cytometry Panel

| Fluorophore | Antigen | Clone | Dilution | Manufacturer | Catalog # |
|---|---|---|---|---|---|
| BV510 | CD45 | 30-F11 | 200 | BioLegend | 103138 |
| BV605 | CD4 | GK1.5 | 300 | BioLegend | 100451 |
| BV786 | PD1 | RPM1-30 | 200 | BD Biosciences | 748264 |
| Spark Blue 550 | CD3 | 17A2 | 100 | BioLegend | 100260 |
| BB700 | CD8α | 53-6.7 | 200 | BD Biosciences | 566409 |
| APC | AH1-Tet | H2L(d) SPSYVYHQF | 100 | NIH Tetramer Core | |
| Zombie NIR | Viability | - | 5000 | BioLegend | 423106 |

**Table 6**: Macrophage Phenotype Flow Cytometry Panel

| Fluorophore | Antigen | Clone | Dilution | Manufacturer | Catalog # |
|---|---|---|---|---|---|
| BV421 | CD86 | GL1 | 200 | BioLegend | 105032 |
| BV510 | Ly6C | HK1.4 | 300 | BioLegend | 128033 |
| BV605 | CD45 | 30-F11 | 125 | BioLegend | 103140 |
| AF488 | CD9 | MZ3 | 200 | BioLegend | 124808 |
| PE | CD301b | URA-1 | 250 | BioLegend | 146804 |
| PE/Dazzle 594 | I-A/I-E | M5/114.15.2 | 300 | BioLegend | 107647 |
| PE-Cy7 | F4/80 | BM8 | 250 | BioLegend | 123114 |
| APC | CD206 | C068C2 | 250 | BioLegend | 141708 |
| AF700 | SiglecF | S17007L | 200 | BioLegend | 155507 |
| eFluor780 | Viability | - | 1000 | Thermo Fisher | 65-0865-14 |

**Table 7**: FACS Panel for Single-Cell RNA-seq

| Fluorophore | Antigen | Clone | Dilution | Manufacturer | Catalog # |
|---|---|---|---|---|---|
| BV421 | CD8a | 53-6.7 | 100 | BD Biosciences | 563898 |
| BV605 | CD45 | 30-F11 | 200 | BioLegend | 103140 |
| BV711 | CD11b | M1/70 | 600 | BioLegend | 101242 |
| PE | CD3 | 17A2 | 150 | BioLegend | 100206 |
| APC | AH1-Tet | H2L(d) SPSYVYHQF | 100 | NIH Tetramer Core | |
| Zombie NIR | Viability | - | 5000 | BioLegend | 123114 |

| Hashing Oligo. | Antigen | Clone | Dilution | Manufacturer | Catalog # |
|---|---|---|---|---|---|
| TotalSeq-C0301 | MHCI; CD45 | M1/42; 30-F11 | 100 | BioLegend | 115861 |
| TotalSeq-C0302 | MHCI; CD45 | M1/42; 30-F11 | 100 | BioLegend | 115863 |
| TotalSeq-C0303 | MHCI; CD45 | M1/42; 30-F11 | 100 | BioLegend | 115865 |
| TotalSeq-C0304 | MHCI; CD45 | M1/42; 30-F11 | 100 | BioLegend | 115867 |
| TotalSeq-C0305 | MHCI; CD45 | M1/42; 30-F11 | 100 | BioLegend | 115869 |
| TotalSeq-C0306 | MHCI; CD45 | M1/42; 30-F11 | 100 | BioLegend | 115871 |
| TotalSeq-C0307 | MHCI; CD45 | M1/42; 30-F11 | 100 | BioLegend | 115873 |
| TotalSeq-C0308 | MHCI; CD45 | M1/42; 30-F11 | 100 | BioLegend | 115875 |
| TotalSeq-C0309 | MHCI; CD45 | M1/42; 30-F11 | 100 | BioLegend | 115877 |
| TotalSeq-C0310 | MHCI; CD45 | M1/42; 30-F11 | 100 | BioLegend | 115879 |
| TotalSeq-C0311 | MHCI; CD45 | M1/42; 30-F11 | 100 | BioLegend | 115881 |
| TotalSeq-C0312 | MHCI; CD45 | M1/42; 30-F11 | 100 | BioLegend | 115883 |

**Table 8**: FACS Panel for Bulk RNA-seq

| Fluorophore | Antigen | Clone | Dilution | Manufacturer | Catalog # |
|---|---|---|---|---|---|
| BV421 | Ly6C | HK1.4 | 200 | BioLegend | 128032 |
| BV605 | CD45 | 30-F11 | 250 | BioLegend | 103140 |
| BV711 | CD11b | M1/70 | 600 | BioLegend | 101241 |
| AF488 | Ly6G | 1A8 | 300 | BioLegend | 127626 |
| PE | CD3 | 17A2 | 150 | BioLegend | 100206 |
| PE-Cy7 | F4/80 | BM8 | 250 | BioLegend | 123114 |
| AF647 | NKp46 | 29A1.4 | 100 | BioLegend | 137628 |
| Zombie NIR | Viability | - | 5000 | BioLegend | 423106 |
